# The whale shark genome reveals patterns of vertebrate gene family evolution

**DOI:** 10.1101/685743

**Authors:** Milton Tan, Anthony K. Redmond, Helen Dooley, Ryo Nozu, Keiichi Sato, Shigehiro Kuraku, Sergey Koren, Adam M. Phillippy, Alistair D.M. Dove, Timothy D. Read

## Abstract

Chondrichthyes (cartilaginous fishes) are fundamental for understanding vertebrate evolution, yet their genomes are understudied. We report long-read sequencing of the whale shark genome to generate the best gapless chondrichthyan genome assembly yet with higher contig contiguity than all other cartilaginous fish genomes, and studied vertebrate genomic evolution of ancestral gene families, immunity, and gigantism. We found a major increase in gene families at the origin of gnathostomes (jawed vertebrates) independent of their genome duplication. We studied vertebrate pathogen recognition receptors (PRRs), which are key in initiating innate immune defense, and found diverse patterns of gene family evolution, demonstrating that adaptive immunity in gnathostomes did not fully displace germline-encoded PRR innovation. We also discovered a new Toll-like receptor (TLR29) and three NOD1 copies in the whale shark. We found chondrichthyan and giant vertebrate genomes had decreased substitution rates compared to other vertebrates, but gene family expansion rates varied among vertebrate giants, suggesting substitution and expansion rates of gene families are decoupled in vertebrate genomes. Finally, we found gene families that shifted in expansion rate in vertebrate giants were enriched for human cancer-related genes, consistent with gigantism requiring adaptations to suppress cancer.

## Introduction

Jawed vertebrates (Gnathostomata) comprise two extant major groups, the cartilaginous fishes (Chondrichthyes) and the bony vertebrates (Osteichthyes, including Tetrapoda) (Venkatesh et al., 2014). Comparison of genomes between these two groups not only provides insight into early gnathostome evolution and the emergence of various biological features, but also enables inference of ancestral jawed vertebrate traits (Venkatesh et al., 2014). The availability of sequence data from many species across vertebrate lineages is key to the success of such studies. Until very recently, genomic data from cartilaginous fishes was significantly underrepresented compared to other vertebrate lineages. The first cartilaginous fish genome, that of *Callorhinchus milii* (known colloquially as ghost shark, elephant shark, or elephant fish), was used to study the early evolution of genes related to bone development and emergence of the adaptive immune system (Venkatesh et al., 2014). As a member of the Holocephali (chimaeras, ratfishes), one of the two major groups of cartilaginous fishes, *C. milii* separated from the Elasmobranchii (sharks, rays, and skates) approximately 420 million years ago, shortly after the divergence from bony vertebrates. Sampling other elasmobranch genomes for comparison is therefore critically important to our understanding of vertebrate genome evolution (Redmond et al., 2018).

Until recently, few genetic resources have been available for elasmobranchs in general, and for the whale shark (*Rhincodon typus*) in particular. The first draft elasmobranch genome published was for a male whale shark of Taiwanese origin by Read *et al*. (2017). Famously representing one of Earth’s ocean giants, the whale shark is by far the largest of all extant fishes, reaching a maximum confirmed length of nearly 19 meters (McClain et al., 2015). Due to its phylogenetic position relative to other vertebrates, the scarcity of shark genomes, and its unique biology, the previous whale shark genome was used to address questions related to vertebrate genome evolution (Hara et al., 2018; Marra et al., 2019), the relationship of gene evolution in sharks and unique shark traits (Hara et al., 2018; Marra et al., 2019), as well as the evolution of gigantism (Weber et al., 2020). A toll-like receptor (TLR) similar to TLR21 was also found in this first whale shark genome draft assembly, suggesting that TLR21 was derived in the most recent common ancestor of jawed vertebrates. While this represented an important step forward for elasmobranch genomics, the genome was fragmentary, and substantial improvements to the genome contiguity and annotation were expected from reassembling the genome using PacBio long-read sequences (Read et al., 2017).

Despite the relative lack of genomic information prior, much recent work has focused upon further sequencing, assembling, and analyzing of the whale shark nuclear genome (Hara et al., 2018; Read et al., 2017; Weber et al., 2020). Hara *et al*. reassembled the published whale shark genome data, and sequenced transcriptome data from blood cells sampled from a different individual for annotation (Hara et al., 2018). Alongside the work on the whale shark genome, genomes have also been assembled for the bamboo shark (*Chiloscyllium punctatum*), cloudy catshark (*Scyliorhinhus torazame*), white shark (*Carharodon carcharias*), and white- spotted bamboo shark (*Chiloscyllium plagiosum*). Comparative analyses of shark genomes supported numerous evolutionary implications of shark genome evolution, including a slow rate of shark genome evolution, a reduction in olfactory gene diversity, positive selection of wound healing genes, proliferation of CR1-like LINEs within introns related to their larger genomes, and rapid evolution in immune-related genes (Hara et al., 2018; Marra et al., 2019; Weber et al., 2020; Zhang et al., 2020).

Long read sequencing is an important factor in assembling longer contigs to resolve repetitive regions which comprise the majority of vertebrate genomes (Koren et al., 2017). Herein we report on the best gapless assembly of the whale shark genome to date, based on *de novo* assembly of long reads obtained with the PacBio single molecule real-time sequencing platform. We used this assembly and new annotation in a comparative genomic approach to investigate the origins and losses of gene families, aiming to identify patterns of gene family evolution associated with major early vertebrate evolutionary transitions. Building upon our previous finding of a putative TLR21 in the initial draft whale shark genome assembly, we performed a detailed examination of the evolution of jawed vertebrate pathogen recognition receptors (PRRs), which are innate immune molecules that play a vital role in the detection of pathogens. Despite their clear functional importance, PRRs (and innate immune molecules in general), have been poorly studied in cartilaginous fishes until now. Given that cartilaginous fishes are the most distant evolutionarily lineage relative to humans to possess both an adaptive and innate immune system, the study of their PRR repertoire is important to understanding the integration of the two systems in early jawed vertebrates. For example, previous work has shown several deuterostome invertebrate genomes possess greater expanded PRR repertoires when compared to relatively conserved repertoires found in bony vertebrates, which suggests that adaptive immunity may have negated the need for many new PRRs in jawed vertebrates (Huang et al., 2008; Rast et al., 2006; Smith et al., 2013), although this hypothesis has not been formally tested. Using the new whale shark genome assembly, we therefore investigated the repertoires of three major PRR families: NOD-like receptors, RIG-like receptors, and Toll-like receptors. Next, we compared the rates of functional genomic evolution in multiple independent lineages of vertebrates in which gigantism has evolved, including the whale shark, to test for relationships between gigantism and genomic evolution among vertebrates. Finally, we studied whether gene families that have shifted in gene duplication rates were enriched for orthologs of known cancer-related genes. Larger-bodied organisms tend to have lower cancer rates than expected given their increased numbers of cells relative to smaller-bodied organisms (Peto et al., 1975), suggesting genes involved in cancer suppression may evolve differently in vertebrate giants. Recent research in giant mammals such as elephants and whales supports this hypothesis and has identified selection or duplication of various gene families that are related to suppressing cancer in humans (Abegglen et al., 2015; Sulak et al., 2016; Tollis et al., 2019). Hence, we studied whether gene families that have shifted in gene duplication rates were enriched for orthologs of known cancer-related genes.

## Results and Discussion

### Gapless Genome Assembly

We added to our previously sequenced short-read Illumina data (∼30X coverage) by generating 61.8 GB of long-read PacBio sequences; relative to a non- sequencing-based estimate of genome size of 3.73 Gbp (Hara et al., 2018), this is an expected coverage of long read sequences of about 16X coverage, for a total of ∼46X coverage. The new whale shark genome assembly represented the best gapless assembly to date for the whale shark (Appendix, Supplementary table 1). The total length of the new assembly was 2.96 Gbp.

The total size of the assembly was very similar to the genome size estimated from the *k*-mer based approach GenomeScope of approximately 2.79 Gbp, suggesting the genome is fairly complete. On the other hand, it was smaller than a non-sequencing based estimate of the whale shark genome size of 3.73 Gbp by Hara *et al*. (2018), which suggests that sections of the genome, potentially comprising primarily repetitive elements, are still missing. Repetitive elements were annotated to comprise roughly 50.34% of the genome assembly (Appendix 1A).

The new assembly had 57,333 contigs with a contig N50 of 144,422 bp, or fewer contigs than the number of scaffolds of previous assemblies, and a higher contig N50, representing a dramatic improvement in gapless contiguity compared to the existing whale shark genome assemblies (Supplementary table 1). This higher contiguity at the contig level (vs. scaffold level) was also better than the published *Callorhinchus*, brownbanded bamboo shark, cloudy catshark, and white shark genomes (Hara et al., 2018; Marra et al., 2019; Venkatesh et al., 2014). The scaffolded genome had 39,176 scaffolds and a scaffold N50 of 344,460 bp. Relative to the previously published most contiguous assembly by Weber *et al*. (2020) while they reported a far higher scaffold N50 of ∼2.56 Gb (3.13 Gb ≥ 200 bp), our scaffolded assembly has far fewer scaffolds (3.3 M sequences, 139,611 sequences ≥ 200 bp) (Supplementary table 1)

Based on GenomeScope, the whale shark genome had an estimated level of heterozygosity of approximately 0.0797–0.0828%, consistent with the k-mer coverage plot showing a unimodal distribution (Supplementary figure 1). Comparison of the k-mer profile with the presences of k-mers in the assembly revealed that they were relatively concordant, with no indication that there were many k-mers that were represented twice in the assembly (which could be due to a diploid individual having phased haplotypes assembled into separate contigs) (Supplementary figure 2). Mapping the reads to the genome assembly and calling SNPs using freebayes provided a similar estimate of 2,189,244 SNPs, a rate of 0.0739% heterozygosity, or an average of a SNP every 1353 bases. This suggests that the heterozygosity of the whale shark genome is relatively low.

To assess gene completeness, we first used BUSCO v2 (Simão et al., 2015). Of 2,586 orthologs conserved among vertebrates searched by BUSCO (Simão et al., 2015), we found 2,033 complete orthologs in the whale shark genome, of which 1,967 were single-copy and 66 had duplicates. 323 orthologs were detected as fragments, while 230 were not detected by BUSCO. With 78.7% complete genes, this represents a marked improvement over the previous whale shark genome assembly, which only had 15% complete BUSCO genes (Hara et al., 2018). We also evaluated gene completeness using a rigid 1-to-1 ortholog core vertebrate gene (CVG) set that is better tuned for finding gene families in elasmobranchs (Hara et al., 2015) implemented in gVolante server version 1.2.1 (Nishimura et al., 2017; accessed 2019 Apr 23) (Nishimura et al., 2017), we found that 85% of core vertebrate genes were complete and found that 97.4% of core vertebrate genes included partial genes, which compares favorably to completeness statistics in other shark assemblies (Hara et al., 2018) (Supplementary table 3). The gene content of this whale shark assembly was thus quite complete and informative for questions regarding vertebrate gene evolution.

#### Ancestral Vertebrate Genome Evolution

We sought to use the new whale shark genome assembly to infer the evolutionary history of protein-coding gene families (i.e. orthogroups)

across vertebrate phylogeny, aiming to provide insight into the evolution of biological innovations during major transitions in vertebrate evolution. Orthogroups are defined as all genes descended from a single gene in the common ancestor of the species considered (Emms and Kelly, 2015); hence, they are dependent to a degree on the phylogenetic breadth of species included in the analysis. Genes from the proteomes for 37 chordate species deriving primarily from Ensembl and RefSeq (O’Leary et al., 2016; Yates et al., 2020) (Supplementary table 4), including 35 representative vertebrates, a sea squirt (*Ciona*), and a lancelet (*Branchiostoma*) were assigned to 18,435 orthologous gene families using OrthoFinder, which were assigned to a mean of 10,688 orthogroups per genome. We then inferred the history of gene family origin and loss by comparing the presence and absence of gene families across species. Although accurate inference of gene family size evolution in vertebrates may benefit from further taxon sampling among invertebrates, *Branchiostoma* is notable among animals for retaining a relatively high number of gene families present in the animal stem branch (Richter et al., 2018), hence it is one of the best outgroup species to vertebrates to study the origins of novel genes within the vertebrate clade.

We inferred a consistent increase in the total number of gene families from the root to the MRCA (most recent common ancestor) of Gnathostomata, but only slight increases following this in the MRCAs of bony fishes and cartilaginous fishes (black numbers, Figure 1, Appendix). We also found numbers of novel gene families increased from the root to a peak in the MRCA of gnathostomes, and then novelty decreased precipitously towards the bony and cartilaginous fish descendants (numbers indicated by + symbol, Figure 1). Gene families conserved in all members of a clade may be considered core genes. There was also an increase in the number of core genes of each ancestor from the most inclusive to the least inclusive clades, as expected with decreasing phylogenetic breadth of the less inclusive clades (black parenthetical numbers, Figure 1). We found that a decreasing number of novel core gene families in vertebrate ancestors were retained between the MRCA of Olfactores (tunicates + vertebrates) and the MRCA of Gnathostomata (parenthetical numbers indicated by +, Figure 1), in contrast to the general pattern of increasing numbers of novel gene families along these same branches. Overall, this implied that the origin of jawed vertebrates established a large proportion of novel gene families of both bony vertebrates and cartilaginous fishes, with variable retention among descendant lineages.

**Figure 1.**
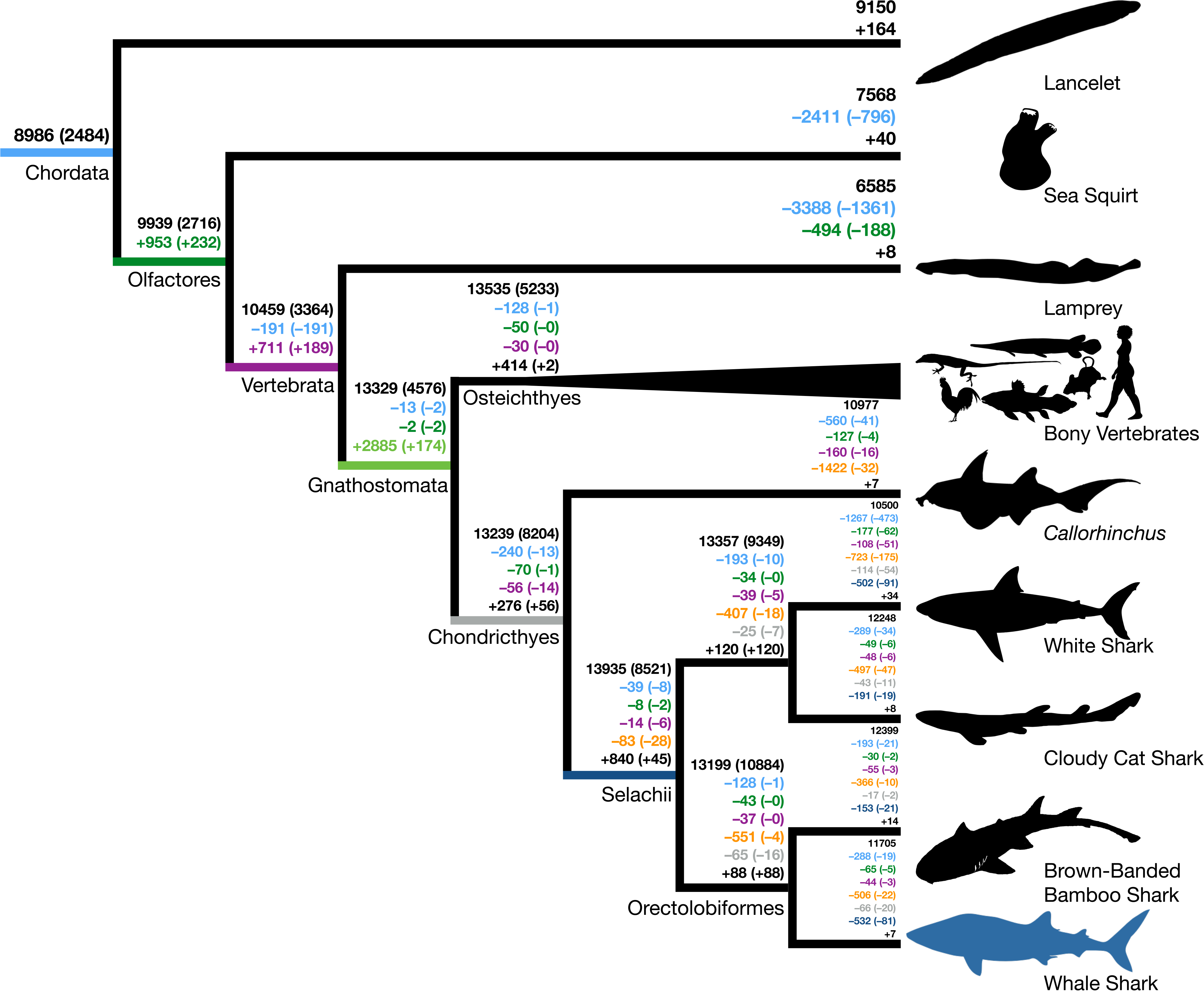
Origins and losses of vertebrate gene families. Above the branch in black is the total number of gene families inferred to be present in the most recent common ancestor at that branch; the number in parentheses indicates the number of gene families conserved in all descendants of that branch. Numbers preceded by + and – indicate the number of gene families inferred to be gained or lost along that branch. Gains and losses are color-coded based on the branch where these gene families originated. Colors indicate gene families present in the most recent common ancestor of chordates, green indicates gene families that originated in the most recent common ancestor of tunicates and vertebrates (Olfactores), purple indicates vertebrate- derived gene families, orange indicates gnathostome-derived gene families, grey indicates chondrichthyan-derived gene families, while dark blue indicates shark-derived gene families. Negative numbers within parentheses indicate gene family losses that are unique to that branch (as opposed to gene families that were also lost along other branches). Positive colored numbers within parentheses indicate novel gene families conserved in all descendants ("core" gene families).

The inclusion of multiple chondrichthyan lineages was important in the inference of gnathostome-derived gene families. The selachians (true sharks) lost fewer gene families than *Callorhinchus* (144 vs. 2,269 overall gene families, 83 vs. 1,422 gnathostome-derived gene families), but future improved taxon sampling may further increase this estimate. Additional high-quality genomes of holocephalans may demonstrate that some of these losses in *Callorhinchus* are lineage-specific or due to assembly or annotation errors, and the addition of batoids (skates and rays) could recover gene families independently lost in holocephalans and selachians. Thus, increasing chondrichthyan taxon sampling allowed for more confidence in the origin and loss of gene families in vertebrate history and assignment of hundreds of genes as having originated prior to the MRCA of gnathostomes.

The burst in emergence of novel gene families that we observed along the ancestral jawed vertebrate branch coincides with the two rounds (2R) of whole genome duplication that occurred early in vertebrate evolution, resulting in gene duplicates referred to as ohnologs (Braasch et al., 2016; Ohno et al., 1968; Singh et al., 2015). Hypothetically, divergent ohnologs may be erroneously assigned to novel gene families and artefactually inflate our estimate for gene family birth along the ancestral jawed vertebrate branch. To estimate the potential extent of ohnolog family splitting, we compared the 2,885 gene families inferred as novel at the base of jawed vertebrates to ohnolog families previously inferred by Singh *et al*. (2015). Generally, most ohnologs had all their copies assigned to single gene families (1131–1609 ohnologs per species). We found that only 157 (5.4% of 2,885 gene families) of gene families that we inferred to have originated in the MRCA of jawed vertebrates corresponded to split ohnologs. Hence, the split ohnologs are not a large proportion of the novel gene families in the jawed vertebrate ancestor. In addition, we also found that only 3 gene families were inferred to be derived at the MRCA of teleosts, coinciding with the teleost-specific genome duplication, which also supports that gene family inference is robust to genome duplication. This finding reinforces the importance of this evolutionary transition for genomic novelty, not just due to the vertebrate 2R whole genome duplication, but also through the addition of novel gene families.

Next, we tested whether gene families that were gained or lost during vertebrate evolution were enriched for certain GO (Gene Ontology) or Pfam annotations (Appendix; Supplementary table 5), potentially indicating functional genomic shifts preceding the origin of these clades. Functional annotations were annotated using InterProScan 5.32-71.0 and Kinfin 1.0 (Laetsch and Blaxter, 2017), and functional enrichment was determined using the Fisher’s exact test. Overall, there were 8,700 gene families (47.2%) annotated for Gene Ontology (GO) functional terms and 14,727 gene families (79.9%) annotated for Pfam protein domains. For example, for the 711 novel gene families in the MRCA of Olfactores, we found an enrichment of connexin function (Supplementary table 6). This is consistent with prior work that determined that the origin of connexin gap junction proteins among chordates was in the MRCA in Olfactores (Alexopoulos et al., 2004). We also found enrichment of ankyrin repeat domains, a motif found widely across eukaryotes which has diverse functions in mediation of protein– protein interactions (Li et al., 2006), and hence may be involved in the evolution of novel protein-complexes in Olfactores. Also, specific to the evolution of the whale shark, we inferred 7 novel gene families and a loss of 1,501 gene families. Neither of these sets of gene families were enriched for any functional terms, suggesting whale shark-specific traits are not attributed to functional genomic shifts due to the origins or losses of gene families.

Among the novel proteins in the MRCA of vertebrates, we found enrichment of several protein domain types including rhodopsin family 7-transmembrane (7-TM) receptor domains, immunoglobulin V-set domain, collagen triple helix repeats, zona pellucida domain, and C2H2- type zinc finger domain (Supplementary table 7; Appendix). The enrichment of collagen function is consistent with the importance of these collagens at the origin of vertebrates and their potential involvement in origin of vertebrate traits, such as bone and teeth (Boot-Handford and Tuckwell, 2003). The enrichment of the zona pellucida domain at the origin of vertebrates is consistent with previous evidence showing that zona pellucida proteins likely originated in vertebrates (Litscher and Wassarman, 2014). Inner-ear proteins also contain the zona pellucida domain, making its appearance in the vertebrate ancestor coincident with the origin of inner ears (Popper et al., 1992). The 7-TM domain proteins include a wide variety of receptors but were not enriched for any particular GO term. Some example receptors include those involved in binding a variety of ligands (e.g. fatty acids, neuropeptides, and hormones), and receptors with immune relevance (e.g. chemokine, bradykinin, and protease-activated receptors). The immunoglobulin V-set domain was found in several proteins, most which had roles in cell adhesion and other functions (Appendix). We also found enrichment among novel vertebrate genes for the C2H2-type zing finger domain, a well-characterized zinc finger domain primarily responsible for nucleotide–protein, as well as protein–protein interactions (Brayer and Segal, 2008; Wolfe et al., 2000). These novel genes were also not enriched for any particular GO term, but play a role in a variety of developmental signaling pathways and cell cycle regulation (Supplementary table 7; Appendix). The enrichment of these varied functional protein domains in the MRCA of vertebrates demonstrate their importance in the origin of diverse vertebrate traits, including responding to stimuli, fertilization, immunity, and signaling. Although the origins of some of these gene families were in the vertebrate ancestor, subsequent gene diversification in jawed vertebrates continued to increase the functional diversity of these gene families, such as in the collagens which were duplicated in the jawed vertebrate genome duplication (Haq et al., 2019; Wada et al., 2006).

In the MRCA of jawed vertebrates, we found enrichment of a variety of immune-related protein domains including immunoglobulin V-set domain, immunoglobulin C1-set domain, and interleukin-8-like small cytokines, with functional enrichment of immune response and hormone activity. Immunoglobulin-domain containing gene families included many immunoglobulins, interleukins, interleukin receptors, T-cell receptors, sialic acid-binding immunoglobulin-type lectins (Siglec proteins), chemokines, cluster of differentiation (CD) proteins, and MHC proteins (Supplementary table 8), consistent with the evolution of immunoglobulin-/T-cell receptor-based adaptive immunity in gnathostomes (Boehm, 2012; Kaiser et al., 2004). We also found enrichment for hormone activity, related to the origin of genes for many hormones at the origin of vertebrates (Supplementary table 8). This finding complemented previous work that identified hormones with a role in mammal homeostasis originating in the MRCA of jawed vertebrates, but it emphasizes that hormone activity is also a predominant function among earlier novel vertebrate gene families (Hara et al., 2018).

Differences between bony vertebrates and cartilaginous fishes might be due to functional differences in gene families specific to each lineage. For gene families exclusive to bony vertebrates (including the 414 gene families derived in the MRCA of bony vertebrates and 366 gene families lost in cartilaginous fishes), we found several enriched sets of protein domains and functions including GCPR domain, lectin C-type domain, and C2H2-type zinc- finger proteins (Supplementary Tables 9–10). GPCRs are 7-TM proteins that transmit signals in response to extracellular stimuli to G proteins (Pierce et al., 2002). This enrichment of GPCR protein function is consistent with the relative paucity of these receptors in cartilaginous fishes, noted previously (Marra et al., 2019). We found many of the GPCRs gained in the MRCA of bony vertebrates were olfactory receptors, which is also consistent with the relatively low number of olfactory receptors noted in cartilaginous fishes previously (Hara et al., 2018; Marra et al., 2019). We also found that one of the GPCR gene families included MAS1 and its relatives. MAS1 is important in response to angiotensin and regulating blood pressure, and although sharks produce angiotensin I (Takei et al., 1993), the precursor to angiotensin-(1-7), the lack of MAS1 and related receptors in cartilaginous fishes suggests that such responses are mediated by alternative receptors and that blood pressure regulation is distinct between cartilaginous fishes and bony vertebrates. Among the lectin C-type domain proteins, we found no orthologs of the NK gene cluster in cartilaginous fishes (e.g. CD69, KLRC), a conserved complex of genes found across bony vertebrates, which implies potential differences in the natural killer complex in cartilaginous fishes (Kelley et al., 2005). Gene families lost in cartilaginous fishes are also enriched for loss of KRAB box domain, which play a role in transcription repression factors (Margolin et al., 1994). There was also enrichment for genes including the C-type lectin domain, which bind a variety of ligands and have functions including playing roles in immunity (Brown et al., 2018). By contrast, we did not find enrichment in domains or functions among the gene families derived in cartilaginous fishes, which may in part be due to fewer annotations among gene families that are not present in bony vertebrates (Appendix). In summary, functional genomic differences between bony vertebrates and cartilaginous fishes are due to differences in the presence of gene families in bony vertebrates, with some related to immunity, chemosensation, and signaling.

Our analyses imply a dynamic history of gene family gain and loss across early vertebrate evolution. Of particular importance was the number of gene families gained in the MRCA of jawed vertebrates in establishing the gene families that are present in bony vertebrates and cartilaginous fishes, with these novel gene families being enriched for immune- related functions. The whale shark genome provided an important additional resource to study the origins of gene families in early vertebrate evolution.

#### Evolution of jawed vertebrate innate immune receptors

Cartilaginous fishes are the most distant human relatives to possess an adaptive immune system based on immunoglobulin antibodies and T cell receptors (Dooley, 2014; Flajnik and Kasahara, 2010). This has driven extensive functional study of their adaptive immune system and an in-depth, although controversial, analysis of the evolution of adaptive immune genes in the elephant shark genome (Dooley, 2014; Redmond et al., 2018; Venkatesh et al., 2014). By comparison, the cartilaginous fish innate immune system has been overlooked (Krishnaswamy Gopalan et al., 2014), despite its importance to understanding the impact that the emergence of adaptive immunity had on innate immune innovation. For example, some previous analyses of deuterostome invertebrate genomes identified greatly expanded PRR repertoires. Yet vertebrate PRR repertoires are considered to be highly conserved, leading to the suggestion that the need for vast PRR repertoires in vertebrates was superseded by the presence of adaptive immunity in vertebrates (Huang et al., 2008; Rast et al., 2006; Smith et al., 2013). Notable exceptions to this include an expansion of TLRs in codfishes due to a proposed loss of CD4 and MHC class II (Solbakken et al., 2017), and expansion of fish-specific NOD-like receptors (NLRs) in some other teleosts (Howe et al., 2016; Stein et al., 2007). As such, we sought to use the whale shark genome to determine whether cartilaginous fishes have a similar PRR set to bony vertebrates, with which they share an adaptive immune system, and also search for evidence of PRR expansions aiming to better understand vertebrate innate immune evolution. To this end, we used BLAST to identify whale shark sequences corresponding to three major PRR families –NLRs, RIG-like receptors (RLRs), and Toll-like receptors (TLRs) – and reconstructed their phylogeny among published, curated vertebrated PRR gene datasets.

NLRs are intracellular receptors that detect a wide array of pathogen- and damage- associated molecular patterns (e.g. flagellin, extracellular ATP, glucose) (Fritz and Kufer, 2015). We identified 43 putative NLRs in the whale shark. We found direct orthologs of almost all human NLRs (UFBOOT=100 for all; Table 1), of which 23 contained a clearly identifiable NACHT domain (a signature of NLRs) (NOD1, NOD2, CIITA, NLRC5, NLRC3) while other putative orthologs did not (NLRX1, NLRC4, BIRC1, NWD1, TEP-1, and NLRP). While inclusion of these sequences lacking an apparent NACHT might seem questionable, the false-negative rate for NACHT domain detection is high, even for some human NLRs. The presence of these orthologs in whale shark indicates the presence of a conserved core NLR repertoire in jawed vertebrates (Supplementary table 13, Figure 2 and Figure 2–figure supplement 1, Appendix). Surprisingly, we found three orthologs of NOD1 in the whale shark, which is a key receptor for detection of intracellular bacteria, rather than a single copy as in humans (ultrafast bootstrap support, UFBOOT=100; Figure 2 and Figure 2–figure supplement 2 4). Further analyses intimate that the three NOD1 copies resulted from tandem duplication events in the ancestor of cartilaginous fishes (Appendix). Sequence characterization suggests that all three of the whale shark NOD1s possess a canonical NACHT domain and so should retain a NOD1-like binding- mechanism, but may have unique recognition specificity (Figure 2–figure supplement 2, Appendix). Thus, we hypothesize that the three NOD1s present in cartilaginous fishes potentiate broader bacterial recognition or more nuanced responses to intracellular pathogens. In contrast to the scenario observed for NOD1, we did not find NACHT domain-containing orthologs of any of the 14 human NLRP genes, many of which activate inflammatory responses (Schroder and Tschopp, 2010), in whale shark, and only a single sequence lacking a detectable NACHT domain (Supplementary table 13, Figure 2 and Figure 2–figure supplement 1, Appendix). However, we did identify an apparently novel jawed vertebrate NLR gene family that appears to be closely related to the NLRPs (UFBOOT=67; Figure 2 and Figure 2–figure supplement 1). This family has undergone significant expansion in the whale shark (UFBOOT=100; Figure 2 and Figure 2–figure supplement 1), and we tentatively suggest that this may compensate for the paucity of true NLRPs in whale shark. Nonetheless, these results imply that the NLR-based inflammasomes in humans and whale sharks are not directly orthologous, and hence that NLR-based induction of inflammation and inflammation-induced programmed cell death (Schroder and Tschopp, 2010) are functionally distinct in human and whale shark. Interestingly, each of the vertebrate species we examined (human, zebrafish, and whale shark) has independently expanded different NLR subfamilies relative to the other species included in the analysis, with NLRP genes expanded in human (clade UFBOOT=99; Figure 2 and Figure 2–figure supplement 1) and the previously-identified ‘fish-specific’ FISNA in zebrafish (clade UFBOOT=86; Figure 2 and Figure 2–figure supplement 1). For the latter, we unexpectedly found a whale shark ortholog (UFBOOT=74), suggesting this gene was present in the jawed vertebrate ancestor and is not a teleost novelty. In all, while we found evidence for a core set of NLRs in jawed vertebrates, our analyses also show that multiple, independent NLR repertoire expansions, with probable immunological-relevance, have occurred during jawed vertebrate evolution despite the presence of the adaptive immune system.

**Table 1.**
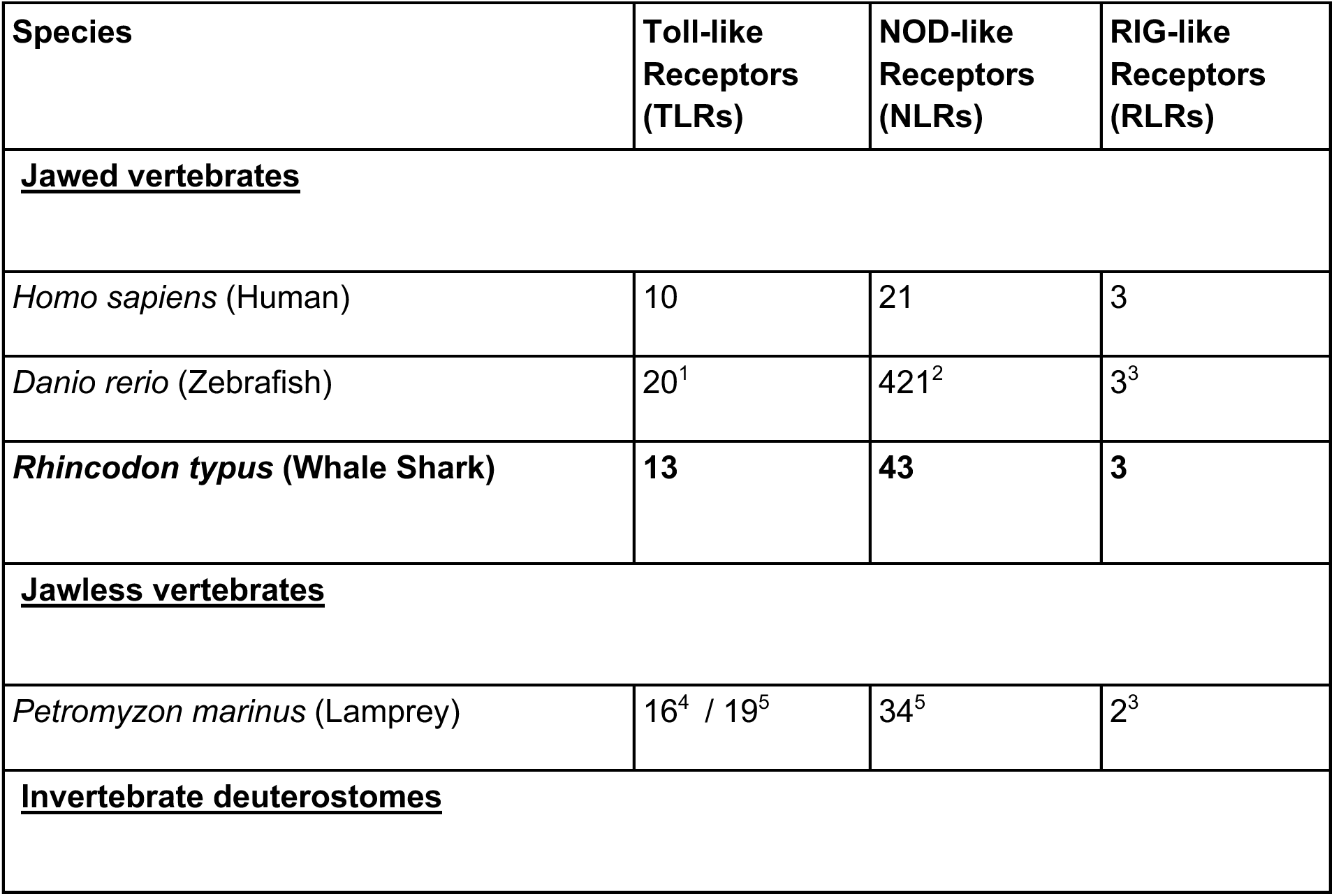
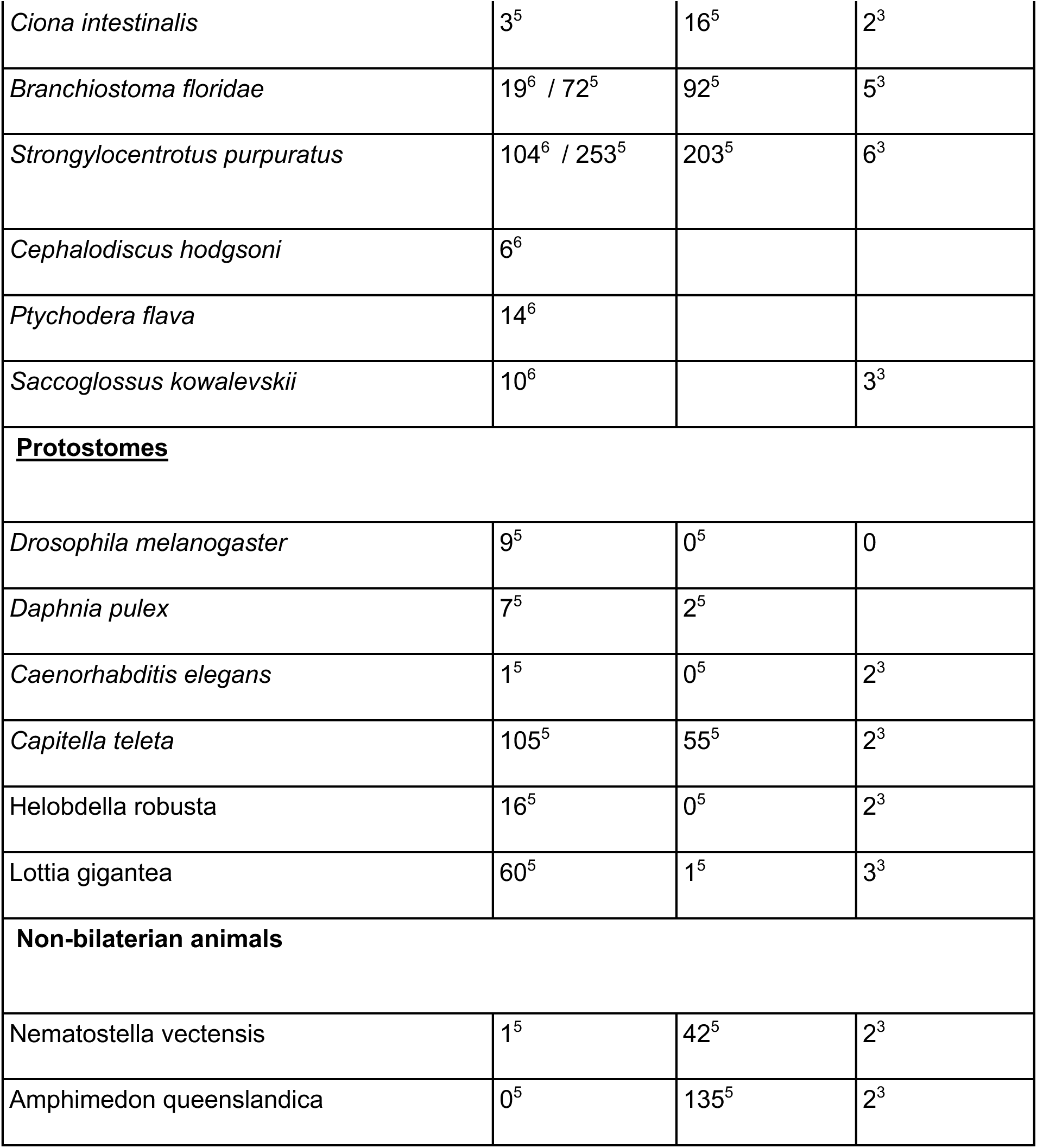
Vertebrate and invertebrate PRR repertoires. Superscripts indicate these citations: 1Chen *et al*. (2021), 2Howe et al. (2016), 3Mukherjee et al. (2014), 4Kasamatsu et al. (2010), 5Buckley & Rast (2015), 6Tassia et al. (2017).

**Figure 2.**
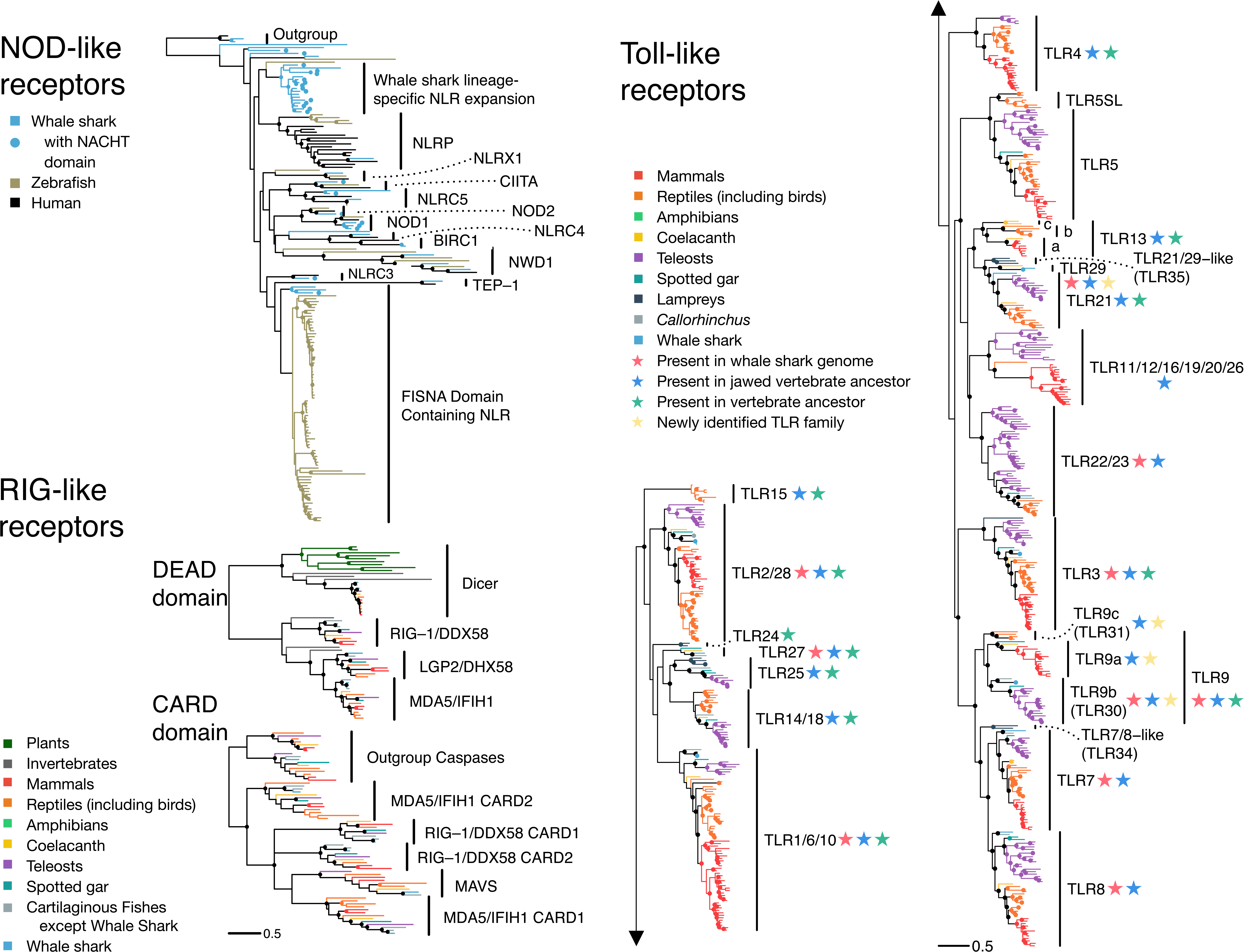
The PRR repertoire of whale shark. Nodes supporetd ≥95% UFBOOT indicated with a dot. For NOD-like receptors, NLRs in whale shark with a NACHT domain are indicated by a dot at the tip. See also Figure2-figure supplement 1-2. For RIG-like receptors, branches are colored by gene, except for RLRs in whale shark which are colored distinctly and labeled by a dot at each tip. See also, Figure2-figure supplement 3. For Toll-like receptors, each clade represents a seperate TLR except families found within TLR13 are also labeled a (TLR13a), b (TLR32), and c (TLR33). TLR families are also labeled by stars indicating whether they were present in the whale shark genome, present in jawed vertebrated ancestor, present in the vertebrate ancestor, and novel to this study. See also Figure2-supplement 4.

RIG-like receptors (RLRs) are intracellular receptors that detect viral nucleic acid and initiate immune responses through Mitochondrial Antiviral Signaling Protein (MAVS) (Mukherjee et al., 2014). Bony vertebrates have three RLR: RIG-1 (encoded by DDX58), MDA5 (IFIH1), and LGP2 (DHX58). Structurally, these are all DEAD-Helicase domain containing family proteins with a viral RNA binding C-terminal RD (RNA recognition domain), and an N-terminal CARD domain pair that mediates interaction with MAVS (Loo and Gale, 2011; Mukherjee et al., 2014). Previous phylogenetic studies either did not included RLRs from cartilaginous fishes or have failed to definitively identify each of the three canonical vertebrate RLRs in this lineage, meaning that the ancestral jawed vertebrate RLR repertoire remained unknown (Krishnaswamy Gopalan et al., 2014; Mukherjee et al., 2014). Our phylogenetic analyses of DEAD-Helicase and CARD domains indicate that orthologs of each of these genes exist in whale shark, revealing that all three RLRs had already diverged in the last common ancestor of extant jawed vertebrates (UFBOOT values all 100; Table 1, Figure 2 and Figure 2–figure supplement 2, Supplementary table 13). Further, and consistent with past findings (Mukherjee et al., 2014), we found that MDA5 and LGP2 are the result of a vertebrate-specific duplication, while RIG-1 split from these genes much earlier in animal evolution (Figure 2 and Figure 2–figure supplement 2, Appendix). We also identified MAVS orthologs in whale shark, elephant shark, and despite difficulties identifying a sequence previously (Boudinot et al., 2014), coelacanth (UFBOOT=100; Figure 2 and Figure 2–figure supplement 2, Supplementary table 13, Appendix). These results show that the mammalian RLR repertoire (and MAVS) was established prior to the emergence of extant jawed vertebrates and has been highly conserved since, consistent with a lack of evidence for large RLR expansions in invertebrates.

Toll-like receptors (TLRs) recognize a wide variety of PAMPs and are probably the best known of all innate immune receptors. While large expansions have been observed in several invertebrate lineages (Huang et al., 2008; Rast et al., 2006), many studies suggest that the vertebrate TLR repertoire is largely conserved (Boudinot et al., 2014; Braasch et al., 2016; Wang et al., 2016). Some teleosts appear to be an exception to this rule; however, this is likely due to the teleost-specific whole-genome duplication, and loss of CD4 and MHC class II in codfishes (Solbakken et al., 2017). We identified 13 putative TLRs in whale shark (Table 1, Supplementary table 13; Appendix) 11 of which are orthologous tp TLR 1/6/10, TLR2/28 (x2) TLR3, TLR7, TLR8, TLR9 (x2), TLR21, TLR22/23, and TLR27 (UFBOOT values all ≥99; Figure 2 and Figure 2–figure supplement 3; Appendix). The remaining two, along with a coelacanth sequence, represent a novel ancestral jawed vertebrate TLR gene family related to TLR21, for which we propose the name TLR29 (UFBOOT = 99; Figure 2 and Figure 2–figure supplement 3). Thus, the whale shark TLR repertoire is a unique combination when compared to all other vertebrates previously studied, being formed from a mix of classic mammalian and teleost TLRs, supplemented with TLR27 and the new TLR29. Our rooted phylogenies indicate that the ancestor of extant vertebrates possessed at least 15 TLRs, while the ancestor of jawed vertebrates possessed at least 19 TLRs (including three distinct TLR9 lineages), both of which are larger repertoires than possessed by modern 2R species (Figure 2 and Figure 2–figure supplement 3; Appendix). Unlike invertebrates, where both loss and expansion of TLRs have been extensive, our data suggest that many jawed vertebrate TLRs existed in the jawed vertebrate ancestor, with lineage-specific diversification of jawed vertebrate TLRs primarily resulting from differential loss (as well as genome duplications), supplemented by occasional gene duplication events.

Overall, our findings imply that the jawed vertebrate ancestor possessed a core set of PRRs that has largely shaped the PRR repertoires of modern jawed vertebrates. We propose the budding adaptive immune system formed alongside this core set of PRRs, with concomitant genome duplication-driven expansion of immunoregulatory genes leading these PRRs to become embedded within new, combined innate-adaptive immunity networks. Our results suggest that the impact of this on the propensity for large expansions is PRR type-specific, with expansion of NLRs being recurrent during jawed vertebrate evolution, and massive expansion of TLRs constrained without degeneration of the adaptive immune system. Although reliance upon innate immune receptors is offset in vertebrates due to the presence of the adaptive immune system, our results suggest that differences in PRR repertoires between vertebrates and invertebrates are driven by specific functional needs on a case-by-case basis. Thus, rather than a simple replacement scenario, the interaction with the adaptive immune system, and associated regulatory complexity, is likely a major factor restraining the proliferation of certain vertebrate PRRs. Although there is clear evidence that PRR expansions can and do occur in jawed vertebrates despite the presence of an adaptive immune system, further in-depth analyses are needed to help better tease out the changes in tempo of PRR diversification across animal phylogeny and whether this associates in any way with the emergence of adaptive immunity.

#### Rates of functional genomic evolution and gigantism

Rates of genomic evolution vary considerably across vertebrates, either across clades or in relationship to other biological factors, including body size. We compared rates in two different aspects of genomic evolution with potential functional relationship to gigantism in the whale shark, to those of other vertebrates: rates of amino acid substitution in protein-coding genes, and rates of evolution in gene family size.

Substitution rates across a set of single-copy orthologs varied across vertebrate genomes, and these rates were relatively low in the whale shark compared to most other vertebrates (Figure 3). We tested for different rates of substitution among vertebrate clades using the two-cluster test implemented by LINTRE (Takezaki et al., 1995). Previous use of this test to compare the elephant shark (*Callorhinchus milii*) genome to other vertebrates determined that *C. milii* has a slower substitution rate than the coelacanth, teleosts, and tetrapods (Venkatesh et al., 2014), and that cartilaginous fishes are slower than bony vertebrates (Hara et al., 2018). We also found that cartilaginous fishes have a significantly slower rate of substitution (p = 0.0004). In addition, *C. milii* (p = 0.0004) was found to be significantly slower than sharks, consistent with prior work (Weber et al., 2020). The whale shark was not significantly different in rate compared to the brownbanded bamboo shark (p = 0.1802), its closest relative included in the analysis. We also found that cartilaginous fishes had a lower rate of molecular evolution compared to subsets of bony vertebrates including ray-finned fishes (p = 0.0004), sarcopterygians (p = 0.001), the coelacanth (p = 0.0004), and tetrapods (p = 0.0088), but not significantly slower than the spotted gar (p = 0.1416). In addition, we found some patterns among other vertebrates consistent with previous studies on rates of molecular substitution, including that ray-finned fishes evolved more rapidly than sarcopterygians (p = 0.0004) and that spotted gar evolved more slowly than teleosts (p = 0.0004).

**Figure 3.**
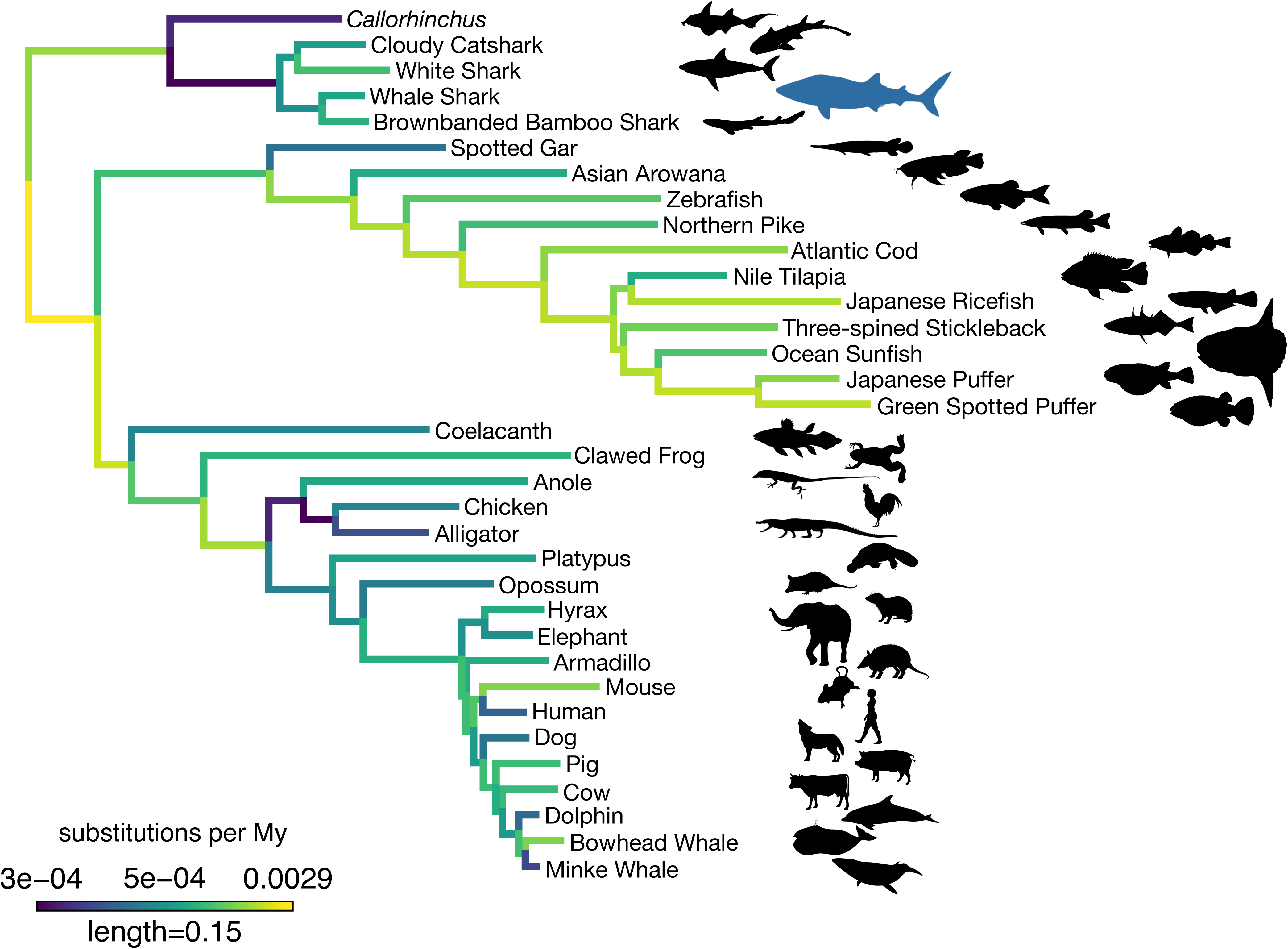
Amino acid substitution rate variation among jawed vertebrates. Branches are colored based on rates quantified by substitutions per site per million years of the maximum likelihood tree compared to a time-calibrated tree. Together, sharks have a slower rate of molecular evolution than *Callorhinchus* (see text on two-cluster test). However, sharks do not have a significantly slower rate of molecular evolution than spotted gar. Furthermore, vertebrate giants– including the whale shark, ocean sunfish, elephant, and whales – have significantly lower rates of molecular evolution than other vertebrates. Note, color scale is on normalized reciprocal-transformed data, which emphasizes changes between smaller values of substitution per My.

We then tested whether rates of molecular substitution differed on those branches leading to gigantism among vertebrates. The origins of gigantism in elephants, whales, and the whale shark have previously been shown to correspond to shifts in the rate or mode of body size evolution (Pimiento et al., 2019; Puttick and Thomas, 2015; Slater et al., 2017). We estimated time-varying rates of body size evolution in cartilaginous fishes using BAMM (Rabosky et al., 2013). Consistent with previous research (Pimiento et al., 2019), we found that gigantism in whale shark corresponds to a discrete shift in the rate of body size evolution to five times the background rate in cartilaginous fishes (Appendix; Supplementary figure 3) (Pimiento et al., 2019). Using PAML to fit models where the rates of amino acid substitution leading to vertebrate giants to other vertebrates differed, we found this model was significantly different than the strict clock (log-likelihood ratio test p = 1.76 x 10^-56^), indicating a significantly different rate of molecular substitution in vertebrate giants. This finding is consistent with earlier evidence that larger-bodied taxa have lower rates of protein evolution (Martin and Palumbi, 1993). However, given that the whale shark genome did not appear to evolve significantly more slowly than the brownbanded bamboo shark genome (noted above), or other small-bodied sharks as found previously when focusing on fourfold degenerate sites (Hara et al., 2018), there may not be an additive effect on substitution rates in the whale shark genome as both a vertebrate giant and a cartilaginous fish. This implied that substitution rates and body size may have less effect in cartilaginous fishes, which are already overall slowly evolving, in contrast to the pattern seen in other vertebrates.

Rates of change in gene family sizes, due to gain and loss of gene copies within gene families, can also vary across species (Han et al., 2013). This represents another potential axis of genomic evolution that is potentially independent from substitution rates. We estimated rates for gene family size evolution for 10,258 gene families present in the MRCA of vertebrates using CAFE 4.2.1 (Han et al., 2013). Average global rates of gene gain and loss in vertebrates were estimated to be 0.0006092 gains/losses per million years. We found that the rate of gene family size evolution in giant vertebrates was significantly faster than in the remaining branches, roughly double the rate in non-giant lineages (p < 0.002). Mean change in gene family size shows that rates of gene family size evolution vary across all taxa including giant lineages, and that an increase in gene family size evolution is not a consistent result of gigantism. However, more complex models in which giant lineages were allowed to vary in rate did not converge. These results suggested that the relationship of gigantism on gene substitution rates do not necessarily predict other forms of genomic evolution, including rates of gene family size evolution.

Replicated shifts in gene gain or loss for specific gene families in independent giant lineages might indicate the consistent effect of selection related to gigantism in particular functional genes. We inferred that 1,387 gene families had a rate shift in gene family size evolution on at least one branch in the vertebrate phylogeny (at p < 0.05, Supplementary file 9). For those gene families that had a rate shift, on average, around 7 independent rate shifts occurred among the vertebrate species considered. Gene families with any shift across the vertebrate phylogeny were enriched for ribosomal genes as well as a few gene families that were enriched for dynein heavy chain genes (Supplementary table 11). No gene families independently shifted in all giant taxa exclusive of other vertebrates, and only five gene families independently shifted in any giant taxa exclusive of other branches in the vertebrate phylogeny (Supplementary file 9). This indicates no consistent signal of selection for a rate shift in gene family size evolution for any particular gene family in the evolution of vertebrate giants.

Interestingly, the gene families that shifted in rates of gene gain and loss anywhere in vertebrate phylogeny were also enriched for human orthologs in the Cancer Gene Census (Fisher’s exact test, odds ratio = 1.43, p = 0.00414) (Sondka et al., 2018), suggesting that these cancer-relevant gene families were also more likely to shift in their expansion/contraction rate across vertebrates. However, this analysis did not consider which branches the shifts occurred on. We hence explored whether gene families that shifted in rate of gene family size evolution across any branch leading to gigantism were enriched for cancer genes. Cancer suppression has evolved by different mechanisms across mammalian lineages, including gene family expansions (Tollis et al., 2017); for example, the duplication of tumor suppressor protein TP53 has been implicated in reduced cancer rates in proboscideans (elephants and relatives) relative to other mammals (Sulak et al., 2016). By contrast, this same gene family is not expanded in baleen whales (Tollis et al., 2017), thus it is already known the same gene families do not expand in all mammalian giants, so we should not expect this to be the case when including fishes. We confirmed a rate shift in gene family size evolution in TP53 in the lineage leading to elephant, but this gene family also had a rate shift for TP53 along the branch leading to minke whale (but not bowhead whale), as well as the non-giant bamboo shark. Therefore, we also tested if all gene families that shifted along a branch leading to a vertebrate giant were enriched for cancer-related genes, including gene families that shifted along non-giant branches. Here, we found that all gene families that shifted in gene gain and loss rate along vertebrate giant lineages (including gene families that shifted along other branches) were enriched for cancer genes relative to all gene families that shifted in any branch in the vertebrate phylogeny (odds ratio = 2.66, p = 0.00199), with over twice as many cancer-related gene families estimated to have shifted in rate among all vertebrate lineages than the null expectation. That these gene families were not enriched for any particular GO function or protein domain implies that cancer- suppression can evolve through various mechanisms. For comparison, we also did the same test but focusing on any gene families that shifted along branches leading to the non-giant vertebrates sister to the giant vertebrates (i.e. brownbanded bamboo shark, pufferfishes, hyrax, bottlenose dolphin), and found that there was no significant enrichment for cancer gene families along those branches (odds ratio = 0.88, p = 0.624). Furthermore, we confirmed the significance of the observed effect size by randomly drawing sets of branches across the vertebrate phylogeny to test for enrichment of cancer genes along random sets of branches, and found that the observed odds ratio of 2.66 was more extreme than 98% of random odds ratios (i.e. p = 0.02). This reinforces that the finding of gene family size evolution shfits along giant branches is significantly enriched for cancer genes.

For the 1,387 gene families that had a significant rate shift in gene family size evolution, we then studied if the rates were significantly greater in cancer genes versus non-cancer genes, depending on whether or not branches led to giant taxa or not. By fitting a linear mixed model using lme4, we found that there was a significantly higher rate of gene family size evolution along branches leading to giant taxa versus other branches (coefficient 0.0203, p = 5.95e^-6^), no effect of whether a gene was related to cancer on rates (coefficient -0.00245, p = 0.219), but a significant interaction of cancer-suppression function and gigantism (coefficient 0.0102, p = 8.51e^-5^), such that rates of gene family size evolution in genes related to cancer leading to giant taxa are even higher than expected relative to the effect of being on a giant branch alone (Figure 4). The significantly higher rate of gene family size evolution in vertebrate giants is consistent with the genome-wide patterns estimated above. In these gene families where a rate shift occurred, we found the mean rate of gene family size evolution along branches leading to giant taxa was 3.32 times greater than the mean rate along other branches in cancer genes, but branches leading to giant taxa had only an average rate that was 2.60 times greater than other branches in genes not related to cancer. Overall, this is suggestive that dynamics of vertebrate evolution in cancer-related gene family size among the sampled taxa are driven by the evolution of gigantism.

**Figure 4.**
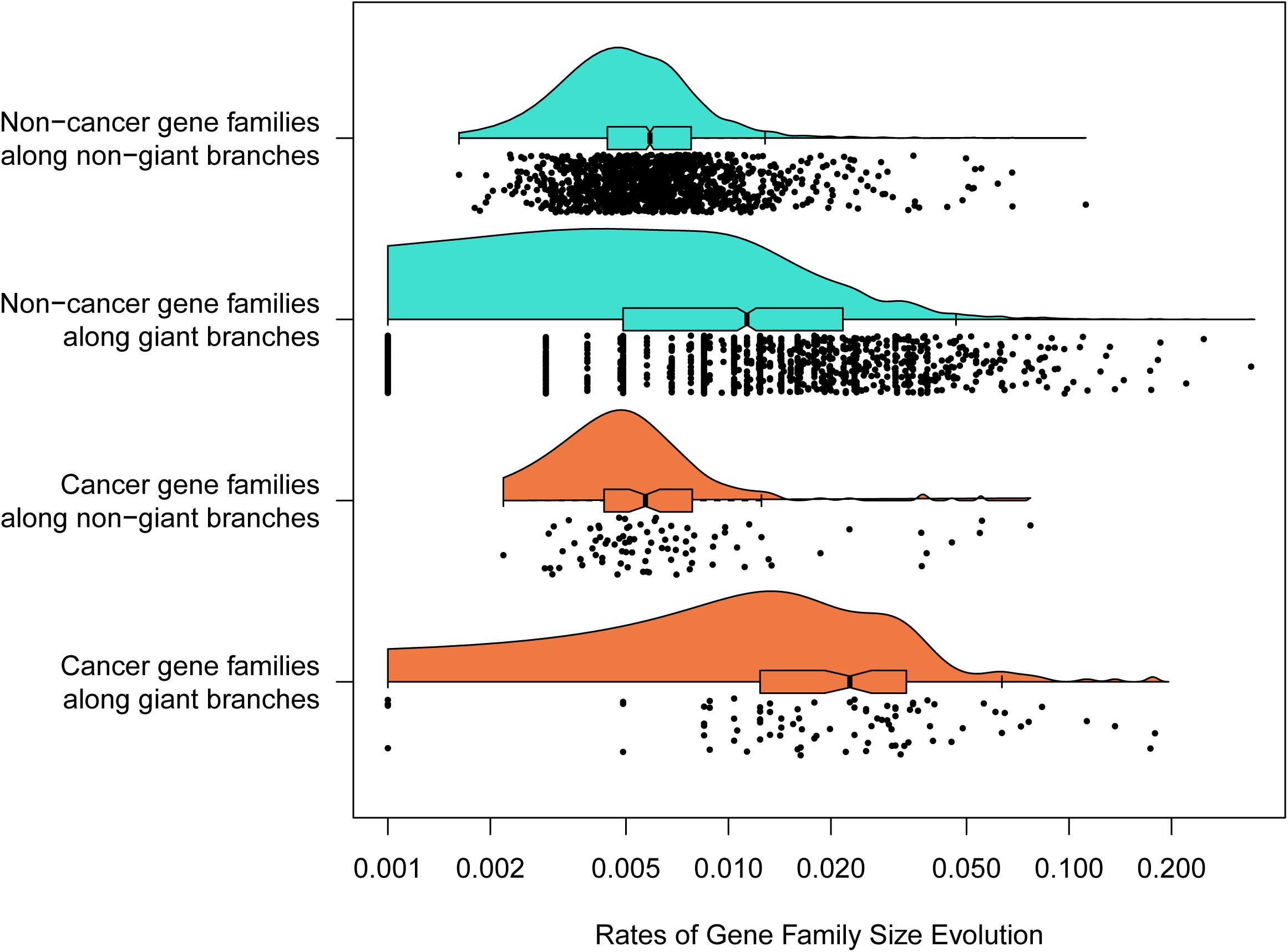
Among 1,387 gene families with a significant rate shift, branch-specific rates of gene family size evolution for branches leading to giant taxa were significantly higher than in branches leading to other taxa, and additionally the rate of gene family size evolution was even greater in cancer-related gene families related to other gene families specifically in branches leading to giant taxa.

## Conclusions

As a representative of cartilaginous fishes, a lineage for which only few genomes have thus far been sequenced, the whale shark genome provides an important resource for vertebrate comparative genomics. The new genome assembly based on long reads we reported in this paper represents the best gapless genome assembly thus far among cartilaginous fishes. Comparison of the whale shark to other vertebrates not only expands the number of shared gene families that were ancestral to jawed vertebrates but demonstrates that the early vertebrate genome duplications were also accompanied by a burst in the evolution of novel genes. These early gene families are involved in a diversity of functions including reproduction, metabolism, development, and adaptive immunity. We also find differences in gene families implying functional genomic differences between bony vertebrates and cartilaginous fishes, MAS1 and the NK gene cluster. With specific respect to genes involved in innate immune protection, we found divergent patterns of gene gain and loss between NLRs, RLRs, and TLRs, which provide insight into their repertoires in the jawed vertebrate ancestor. These results rejected a scenario where the importance of PRRs is muted in vertebrates by the presence of adaptive immunity, instead indicating the ongoing necessity of ancient PRRs, which were integrated with the new adaptive immune system in the jawed vertebrate ancestor. Finally, we demonstrated that the relationship between rates of gene family size evolution and rates of substitution to gigantism are decoupled, and that gene families that shifted in gene expansion and contraction rate leading to vertebrate giants were enriched for genes with cancer relevance. The whale shark genome helps to build a foundation in shark and vertebrate comparative genomics, which is useful to answer questions of broader vertebrate evolution and convergent evolution of distinctive traits. Further sequencing of high-quality elasmobranch genomes will continue to enhance research from finding unique, whale shark-specific evolutionary change to illuminating broader patterns of vertebrate evolution.

## Acknowledgements

The sequencing service was provided by the Norwegian Sequencing Centre (www.sequencing.uio.no), a national technology platform hosted by the University of Oslo and supported by the "Functional Genomics" and "Infrastructure" programs of the Research Council of Norway and the Southeastern Regional Health Authorities". We thank F. Thibaud-Nissen for assistance with genome annotation through NCBI RefSeq. We thank B. Morgan and the High Performance Computing oversight committee for access and assistance with the Center for Advanced Science Innovation and Commerce (CASIC) supercomputer at Auburn University, R. A. Petit III for assistance with computing at Emory University, and High Performance Computing at George Washington University for assistance with Colonial One and Pegasus. We thank the staff of Laboratory of Phyloinformatics in RIKEN BDR for transcriptome sequencing. Inference of gigantism in the whale shark was made possible by body size data kindly provided by C. Mull. We are thankful for funding provided from the Georgia Aquarium and the Emory School of Medicine Development. S. Koren and A.M. Phillippy were supported by the Intramural Research Program of the National Human Genome Research Institute, National Institutes of Health. Silhouettes used throughout via PhyloPic: *Callorhinchus*, CC BY-SA by Milton Tan, originally by Tony Ayling; whale shark, CC BY-SA by Scarlet23 (vectorized by T. Michael Keesey); spotted gar, tilapia, stickleback, CC BY-NC-SA by Milton Tan; Asian arowana, coelacanth, CC BY-NC- SA by Maija Karala; clawed frog, anole, platypus, opossum, elephant, CC BY by Sarah Werning; alligator, CC BY-NC-SA by Scott Hartman; mouse, CC BY-SA by David Liao; dolphin, CC BY-SA by Chris Huh; catshark, brownbanded bamboo shark, white shark, zebrafish, pike, cod, ricefish, ocean sunfish, chicken, armadillo, hyrax, human, dog, pig, cow, minke whale, bowhead whale, public domain.

## Author Contributions

M.T. contributed writing of the manuscript, experimental design, and implementing analyses of genome polishing, phylogenomics, orthology determination, gene family evolution, RNAseq mapping, and submission of genomic data to NCBI for accessioning and annotation by RefSeq. A.K.R. and H.D. performed analyses and wrote the results, supplementary text, and methods sections on innate immune genes. S.Koren and A.P. assembled the genome using Canu. R.N., K.S., and S.Kuraku generated whale shark blood transcriptome data. T.D.R. and A.D.M.D. initiated and managed the genome project. All authors provided feedback on the manuscript.

## Methods

### Genome sequence assembly and assessment

To improve on our earlier efforts to sequence and assemble the whale shark genome (Read et al., 2017), we generated PacBio long read sequences from the same DNA sample. These sequences are available on NCBI SRA under the accession SRX3471980. This resulted in 61.8 Gbp of sequences, equivalent to ∼20x fold coverage. The initial assembly was performed using Canu 1.2 (Koren et al., 2017) with adjusted parameters to account for the lower input coverage:

canu -p asm -d shark genomeSize=3.5g corMhapSensitivity=high corMinCoverage=2 errorRate=0.035

Illumina reads from all paired end read libraries from Read *et al*. were trimmed using Trimmomatic v0.39 with the following settings: ILLUMINACLIP:adapters.fa:2:30:10 LEADING:5 TRAILING:5 SLIDINGWINDOW:4:15 MINLEN:31 where adapters.fa is a fasta file containing all Illumina sequence adapters packaged with Trimmomatic (Bolger et al., 2014). Illumina reads from the single mate pair library from Read *et al*. were trimmed using NxTrim v0.4.3 using default settings (O’Connell et al., 2015). All Illumina reads were aligned to the genome using BWA-MEM (Li, 2013) v0.7.12-r1039 with default parameters and alignments were used as input into Pilon v1.18 under default settings (Walker et al., 2014) to correct errors in the draft assembly. All reads were then used to scaffold the genome using Platanus 1.2.4 (Kajitani et al., 2014). The runs for each library were provided as separate input libraries to platanus scaffold such that the insert sizes will be considered to be different for each library, and the resulting scaffolded assembly was passed to platanus gap_close, both with default settings. Genome size and quality statistics were computed using QUAST v5.0.2 on default settings (Gurevich et al., 2013) and compared to the published values from prior studies. The white shark genome paper did not report contig N50; we decomposed the scaffolds into contigs and determined N50 using seqkit v0.10.1 (Shen et al., 2016).

We performed k-mer analysis using Jellyfish version 2.2.6 on all Illumina reads using the-C setting to count canonical k-mers, a k-mer size (-m) of 21 and hash size (-s) of 100M (Marçais and Kingsford, 2011). Then, we used GenomeScope to fit a model that allows for assembly-free estimation of genome size, heterozygosity, and repeat content (Vurture et al., 2017), providing the k-mer size of 21, read length of 100, and using the default maximum *k*-mer coverage of 1000 (GenomeScope accessed 2017 June 05). We used KAT v2.2.0 to plot the k- mers and visualize the copy number in the genome of k-mers in the raw read Illumina data (Mapleson et al., 2016). We first used kat comp to compare the Jellyfish *k*-mer counts to the genome assembly, and plotted the results using kat plot spectra-cn.

We assessed gene completeness with conserved vertebrate orthologs using BUSCO v2 (Simão et al., 2015) and CVG orthologs using gVolante (version 1.2.1; accessed 2019 April 23) (Hara et al., 2015), and by mapping RNA-seq reads (Appendix). The trimmed reads were then re-used to call SNPs to assess heterozygosity using freebayes under default settings (Garrison and Marth, 2012). We then used vcflib packages vcffilter to filter the results for a minimum quality of 20 (-f "QUAL >20") and vcfstats to count the number of SNPs (Garrison et al., 2021).

### Transcriptome sequencing

Approximately 30 million short read pairs for whale shark transcripts were obtained with paired-end 127 cycles from blood cells of a male and a female by the Illumina HiSeq 1500, as described previously (Hara et al., 2018). Animal handling and sample collections at Okinawa Churaumi Aquarium were conducted by veterinary staff without restraining the individuals, in accordance with the Husbandry Guidelines approved by the Ethics and Welfare Committee of Japanese Association of Zoos and Aquariums. Downstream handling of nucleic acids was conducted in accordance with the Guideline of the Institutional Animal Care and Use Committee (IACUC) of RIKEN Kobe Branch (Approval ID: H16-11). Transcriptome sequence data are available at NCBI BioProject ID PRJDB8472 and DDBJ DRA ID DRA008572.

### Gene prediction

Genes predictions were provided to us by RefSeq using their genome annotation pipeline version 7.3 (Warren et al., 2017), details of the resulting annotation are publicly available (“Rhincodon typus Annotation Report,” 2018). This annotation included alignments of RNAseq data from grey bambooshark *Chiloscyllium griseum* kidney and spleen, nurse shark *Ginglymostoma cirratum* spleen and thymus, and brownbanded bambooshark *Chiloscyllium punctatum* retina, as well as protein alignments from Actinopterygii, and RefSeq protein sequences for Asian arowana *Scleropages formosus*, coelacanth, spotted gar, zebrafish, clawed frog, and human. After preliminary orthology determination, we determined additional genes absent in whale shark conserved among vertebrates, which we annotated by aligning protein sequences from these genes from human, gar, coelacanth, and mouse (Table S3) to whale shark using genBLAST v1.39 (She et al., 2011) with these settings: -p genblastg -e 1e-5 -g T -gff -cdna -pro -pid (Appendix, Supplementary file 3).

### Orthology inference

We identified orthologs from the whale shark genome by comparison to publicly available chordate genomes. We compiled chordate proteomes for 32 species representing major vertebrate clades, the sea squirt *Ciona intestinalis*, and lancelet *Branchiostoma floridae* (Supplementary table 4). In selecting representative vertebrates, we specifically included the ocean sunfish, African elephant, and two baleen whale genomes (minke whale, bowhead whale), and the most closely-related genomes available for these taxa (*Takifugu rubripes* and *Dichotomyctere nigroviridis*, rock hyrax, and bottlenose dolphin). These ortholog clusters were used for the identification of origins of gene families in chordate evolution and genes that originated in the most recent common ancestor of jawed vertebrates, studying enrichment or changes in functional annotation associated with these orthogroups, phylogenomics, estimation of rates of molecular substitution, and estimation of rates of gene duplication and loss.

Ortholog clusters from proteomes were determined using OrthoFinder v2.2.6 with default settings (Emms and Kelly, 2015). With the resulting hits, OrthoFinder adjust scores for reciprocal best hits while accounting for gene length bias and phylogenetic distance, then proceeds with clustering genes into orthogroups. Preliminary orthology determination suggested many potential missing orthologs in the elephant shark and whale shark genomes. We thus performed orthology-based annotation using genBLAST (She et al., 2011) as noted under Gene prediction methods, added newly identified proteins to the proteomes of whale shark and *Callorhinchus*, and reran the OrthoFinder pipeline including these proteins.

All proteins were then annotated for Gene Ontology (GO) and Pfam terms using InterProScan 5.32-71.0 (Jones et al., 2014), and representative annotations were assigned to each chordate orthogroup using KinFin 1.0 with the --infer-singletons option on to interpret gene families absent from clusters as singletons, then running the functional_annotation_of_clusters.py script packaged with KinFin under default settings, which assigns an annotation to a gene family if at least 75% of proteins in the gene family has that annotation, and at least 75% of taxa within the cluster have a protein with that annotation (Laetsch and Blaxter, 2017) (Supplementary file 6).

### Gene Family Origin and Loss

A custom R script is provided for analyses run for this section (Supplementary file 10). To infer when gene families (as inferred from OrthoFinder) were gained and lost in vertebrate evolution, we mapped the origins and losses of gene families to the species tree parsimoniously, assuming that gene families have a single origin, but can be lost (Laetsch and Blaxter, 2017). We were then able to count the number of gene families present at the MRCA of nodes, the number of novel gene families that originated along each branch, and the number of gene families lost along each branch (including gene families uniquely lost along each branch). We also determined the number of gene families conserved in all descendants (core genes), and the number of novel gene families conserved in all descendants (novel core genes).

To confirm that the number of novel gene families in the jawed vertebrate ancestor was not inflated by artefactual oversplitting of ohnologs (gene duplicates that arose from two rounds of whole genome duplication early in vertebrate evolution). Singh *et al. (2015)* independently used a synteny-aware method to identify ohnologs in a subset of vertebrate genomes. We compared our assignment of human orthologs to gene families to the assignment of human orthologs to Singh *et al*. ohnolog families (downloaded 2020 June 2). We determined whether our gene families and Singh *et al*. ohnolog families matched and whether human orthologs were assigned to a single gene family or ohnolog. We replicated this for green anole, spotted gar, zebrafish, and possum ohnologs. To find the common genes between the Ensembl protein IDs we clustered and Ensembl gene IDs provided by Singh *et al*., we used the R package biomaRt v2.45.2 to translate identifiers (Durinck et al., 2005).

Based on the representative annotations for each orthogroup determined above, we then determined whether groups of gene families that were gained or lost along branches in the vertebrate phylogeny were enriched for certain functions using a Fisher’s Exact Test. Within each comparison, we adjusted the p-value to correct for multiple hypothesis testing by the Benjamin-Hochberg method using the p.adjust function in R (Wright, 1992). Corrected p-values under the BH method can be interpreted at a significance threshold that is equivalent to the false discovery rate. We considered functions enriched with an adjusted p-value of 0.05 and false discovery rate of 0.05.

### Innate Immunity Analyses

*Homology identification*: Sequence similarity searches were performed using BLAST+ v2.6.0 to identify putative homologs of TLRs, NLRs and RLRs (Altschul et al., 1990). An alternative approach using profile hidden Markov models, HMMER [version 3.1] (Eddy, 1998), was also tested for TLRs; the results obtained were identical, except that BLAST returned an additional putative TLR. Due to this HMMER results were not applied in subsequent analyses, and HMMER was not applied elsewhere(Eddy, 1998). Searches for whale shark TLR and RLR homologs were performed using all other sequences present in the TLR and RLR trees. Retention of sequences for further analyses was reliant on a reciprocal blast hit to a TLR or RLR in the Swissprot reviewed database or the NCBI non-redundant protein set (Boeckmann et al., 2003).

For NLRs, detection is more complicated, as some NLRs do not contain computationally detectable NACHT domains (i.e. some family members, even in humans, are false negatives in domain-based search tools and databases), despite the NACHT domain being the defining feature of NLR family members. Further, some of these genes contain other domains and are also included in other gene families where most members do not contain NACHT domains. As such, for the main analysis performed here, those sequences in the predicted proteome and translated transcriptome containing a predicted NACHT domain according to the NCBI CD- search webserver (Marchler-Bauer et al., 2015) are noted as such (and should be considered as the conservative set of whale shark NLR-like sequences). Additional sequences from the predicted protein set with a blast hit to known NLRs were also included to permit detection of potential orthologs of NLRs not found in the conservative set with definite/detectable NACHT domains. Proteins containing the closely related NB-ARC domain were also extracted from the whale shark proteome

In cases where a transcript matches the genomic location of a predicted protein, the predicted protein is the sequence reported. Where multiple predicted proteins refer to the same genomic location, only a single sequence is retained for further analysis.

*Phylogenetic Datasets:* For NLRs, we performed phylogenetic analyses of the whale shark putative NLRs to known NLRs from human and zebrafish, both of which are highly phylogenetically relevant and well-studied in this regard. Proteins containing the closely related NB-ARC domain were used as an outgroup in NLR analyses, along with Human APAF-1 which also harbors an NB-ARC domain (Urbach and Ausubel, 2017).

To better understand RLR and MAVS evolution we used two datasets. The first of these was based on the central DEAD-helicase domains (hence, excluding MAVS) to define which of the three RLR proteins could be found in whale shark, and infer the jawed vertebrate RLR repertoire, also following Mukherjee et al. (2014). For the RLR datasets, members of each of the three vertebrate RLR families, some invertebrate RLRs, and a selection of DICER proteins sequences as an outgroup (Mukherjee et al., 2014) were gathered to generate a phylogenetically informative dataset (i.e. aiming to include representatives of each of the major vertebrate classes for which genome data were available). Full length proteins were aligned for phylogenetic analysis of DEAD-Helicase domains, and trimmed to the start and end of these domains based on the three human RLR sequences (Mukherjee et al., 2014). The second dataset was based on individual CARD domains, as the presence of two CARD domains in RIG-1 and MDA5 is thought to have come about through independent domain duplication in each lineage, which would mislead phylogenetic analyses if ignored (Korithoski et al., 2015) The same process as for DEAD-Helicase domains was performed for the CARD domains (Korithoski et al., 2015).

For the TLR dataset, a large set of TLR nucleotide sequences were taken from a past study that densely sampled vertebrates (Wang et al., 2016). TLR sequence from grey bamboo shark (*Chiloscyllium griseum*) was also included (Krishnaswamy Gopalan et al., 2014). Following trimming, the alignment consisted almost entirely of sites from the TIR domain, so TIR domains were not specifically extracted for this analysis.

For the NLR analysis the described set of human NLRs and NACHT domain containing proteins, as well as the closely related NB-ARCs as an outgroup (Urbach and Ausubel, 2017), were downloaded from NCBI protein database. Sequences of zebrafish, where NLRs are massively expanded (Howe et al., 2016; Stein et al., 2007), were also included in this analysis, but these were downloaded from the InterPro website (i.e. all *Danio rerio* proteins containing a NACHT domain) (Hunter et al., 2009). A very large number of zebrafish sequences were obtained, so to reduce the prevalence of pseudo-replicate sequences (that are likely to be uninformative in the context of understanding the whale shark NLR repertoire), CD-HIT (Fu et al., 2012) was used to cluster zebrafish sequences with greater than 75% identity prior to phylogenetic analysis. An additional NLR analysis was performed focusing specifically on NOD1s, this employed NOD2 as an outgroup based on the larger NLR analysis and included NOD1s identified by BLAST in elephant shark. Notably, our NLR analsyis relies on poorer taxon sampling compared to that for the RLR and NLR datasets. This is due to a relative paucity of previously characterized NLR repertoires across vertebrate species. Importantly, although this does not lend itself well to understanding the tempo of lineage-specific gene family expansion/contraction it does not preclude detection of such events along the lineages leading to the species included in the analysis.

#### Multiple sequence alignment and phylogenetic analyses

Multiple sequence alignments were generated with MAFFT (version 7.313) (Katoh and Standley, 2013) using default parameters for the larger TLR and NLR datasets, but using the more intensive L-INS-i method for RLRs and the focused NOD1 NLR dataset. trimAl (version 1.2rev59) (Capella-Gutierrez et al., 2009) was applied to remove gap rich sites, which are often poorly aligned, from the alignments using the “gappyout” algorithm. BMGE (version 1.12) (Criscuolo and Gribaldo, 2010) was then used to help minimize the number of saturated sites in the remaining alignment (as identified using the BLOSUM30 matrix). The RLR analyses were not subjected to this BMGE analysis, as these were derived from conserved domains (meaning that alignments were based on relatively conserved sequence tracts and were already quite short). The NOD1 focused NLR alignments were judged to contain relatively similar sequences and were not subjected to either trimAl or BMGE analyses. Phylogenetic analyses were performed in IQ-TREE (version: omp- 1.5.4) (Nguyen et al., 2015) using 1000 ultrafast bootstrap replicates (Minh et al., 2013) and the best-fitting model of amino acid substitution. Best-fitting substitution models were determined according to the Bayesian information criterion with ModelFinder from the IQ-TREE package (Kalyaanamoorthy et al., 2017), and ultrafast bootstrap support was computed to assess branch support (Hoang et al., 2017). The following (best-fitting) models were applied for each dataset: LG+I+G for RLR CARD domains dataset, LG+I+F+G for RLR DEAD-Helicase domains dataset, JTT+I+F+G for the TLR dataset, JTT+F+G for the NLR dataset, and JTT+I+F+G for the NOD1 focused NLR dataset. The trees were rooted either by outgroup, or the TLR tree was rooted minimal ancestor deviation method (Tria et al., 2017). This is unlike many other TLR trees produced in previous studies which are unrooted (Roach et al., 2005; Wang et al., 2016)

### Phylogenomics

Orthogroups were filtered to single-copy orthologs for phylogenomic analyses. We determined orthologs from orthogroups by reconstructing orthogroup trees and used tree-based orthology determination using the UPhO pipeline (Ballesteros and Hormiga, 2016). The paMATRAX+ pipeline bundled with UPhO was used to perform alignment (mafft version 7.130b), mask gaps (trimAl version 1.2), remove sequences containing too few unambiguous sites, and check that at the minimum number of taxa are present (using the Al2Phylo script part of UPhO), and then reconstruct phylogenies (IQ-TREE v1.6.10) (Capella-Gutierrez et al., 2009; Katoh and Standley, 2013; Nguyen et al., 2015). Next, we used UPhO to extract orthologs by identifying all maximum inclusive subtrees from orthogroups with at least five species, with the allowance for in-paralogs (paralogs that arose after all species divergences in the phylogeny, and thus do not affect relative relationships in the phylogeny), and retained the longest in- paralogous sequence for each species within each ortholog. For each single-copy ortholog, we aligned, trimmed, and sanitized sequences using the paMATRAX+ pipeline.

To select the most reliable sequences for infer a phylogenomic time tree, we filtered for the most informative loci using MARE version 0.1.2-rc with default settings except -t (taxon weight) set to 10 to weight retaining taxa higher the alignment over retaining loci (Misof et al., 2013). Next, orthologs without lamprey, *Callorhinchus*, whale shark, *Branchiostoma*, and *Ciona* were excluded. We also filtered down to loci that supported the monophyly of vertebrate, gnathostome, chondrichthyan, and osteichthyan clades. After our filtering we were left with an alignment comprising 281 loci and 209,275 residues. We concatenated the sequences and selected the best model of amino acid substitution and partitioning scheme and inferred a maximum likelihood phylogeny using IQTREE v1.6.10 (Hoang et al., 2017; Kalyaanamoorthy et al., 2017; Nguyen et al., 2015) with the followings settings: -bb 1000 -bnni -m MFP+MERGE - rcluster 10. The tree was rooted using the amphioxus *Branchiostoma*. The phylogeny was largely consistent with consensus arising from phylogenomic studies. We also inferred a phylogeny accounting for incomplete lineage sorting using ASTRAL v5.7.1 (Zhang et al., 2018) based on gene trees (not shown), which was identical in topology except for the placement of armadillo (Xenarathra) sister to Afrotheria in the ASTRAL tree versus armadillos sister to Boreoeutheria in IQ-TREE. This relationship has historically been difficult to reconstruct and is consistent with prior conflicts between concatenated and coalescent-based analysis on the placement of Xenarthra with far more taxa (Esselstyn et al., 2017). In addition, none of our focal results are reported within mammals, where the relationships of Xenarthrans could be relevant.

Numerous fossil-based node calibrations were identified from the literature. Most node ages were derived from age ranges published in the Fossil Calibration Database (Benton et al., 2014) and are listed in Supplementary table 12. While previously the age of crown Chondrichthyes (here, the MRCA of Holocephali + Elasmobranchii) has been suggested to range from 333.56–422.4 Ma, the minimum age was recently pushed further back to 358 Ma based on multiple holocephalan fossils (Coates et al., 2017). To assess the concordance of the fossil calibrations, we used treePL version 1.0 to estimate divergence times from the ML tree with each fossil calibration using penalized likelihood, then performed cross-validation and evaluated the concordance of the fossils to the time tree to identify and exclude outliers (Near et al., 2005). After excluding fossils that were discordant with the others, we estimated divergence times using treePL with the remaining fossil calibrations. The final treePL config file is provided (Supplementary file 10).

### Tests for Rates of Substitution

Based on our aligned matrix from single-copy orthologs used for phylogenomics, we tested for differences in rates of molecular substitution between vertebrates by using the two-cluster test implemented in LINTRE (2010 Apr 17 version) (Takezaki et al., 1995), using amino acid p-distances between taxa to estimate branch lengths. The two-cluster test is designed to test if the rates in two clades are significantly different by comparison to an outgroup. We tested rates on the full tree, as well as focused on certain cluster pairs by subsetting the dataset to focus on specific clades for comparison. Sequences were converted to phylip format from fasta format using pxs2phy using phyx v1.01 (Brown et al., 2017).

We also compared rates of genomic evolution of four independent instances of vertebrate gigantism (whale shark, elephant, baleen whales, ocean sunfish) relative to the background rate of molecular evolution among vertebrates. To do this, we used PAML 4.9i to compute the likelihoods of the alignment of single-copy orthologs used for phylogenomics under two different models of molecular evolution (Yang, 2007). We computed the likelihood of the data under a strict clock model (single-rate model) and under a local clock model (two-rate model) where the clock rate differed on branches leading to vertebrate giants. We then determined significance using the likelihood ratio test. PAML control files are provided (Supplementary file 10).

### Rates of Gene Family Size Evolution

We estimated rates of gene family expansion and contraction across vertebrates among gene families. OrthoFinder output includes counts of the size of each orthogroup (i.e. gene families) for each species. We analyzed the evolution of gene family size under a birth-death process using CAFE version 4.2.1 (Han et al., 2013), with the gene family size evolutionary rate parameter λ. We focused on gene families present in the most recent common ancestor of vertebrates and filtered these only to gene families present in at least two species, and to exclude gene families that exceed 100 copies in any species, as large gene families have too large variance for consistent rate parameter estimation, resulting in 10,258 gene families. We used the caferror.py script to estimate species-by-species error rates in the annotation to improve the accuracy of rate estimation (Han et al., 2013). We used a time- calibrated phylogeny of vertebrates for this analysis (see above). We provide scripts used for running CAFE (Supplementary file 10).

We estimated rates of gene duplication and loss across vertebrates under a single λ model, and two multi-λ models: a two λ mode model where branches leading to gigantism had a second rate, and a five λ model where the rate categories were the background and a separate rate for each of the four independent origins of gigantism. However, the five λ model did not converge. To test for significance of the observed difference in likelihoods between the two λ model and the single λ model, we simulated gene family evolution with 500 replicates under these models and estimated the log-likelihood ratios from this null, simulated distribution. The p- value corresponds to the proportion of simulated replicates which had a smaller log-likelihood ratio than observed. When fitting the λ model, CAFE 4 additionally computes rates of duplication and loss along each branch for each gene family, and tests whether significant rate shifts occur along each branch (Supplementary file 10). P-values < 0.05 indicate a significant rate shift in gene family size evolution rate.

We identified gene families that had shifted and tested whether they were enriched for GO and Pfam terms (as above). We also tested for enrichment of gene families including human orthologs related to cancer. Cancer-related gene families were determined by downloading the gene families from the COSMIC Cancer Gene Census (Sondka et al., 2018) and determining which orthogroups included the human ortholog based on the Ensembl gene identifier provided by the CGC (database version 91 2020 April 07, accessed 2020 June 3). Ensembl gene ENSG identifiers were matched to the Ensembl protein ENSP identifiers (which we used for orthogroup determination) using biomaRt version 2.45.2, database accessed 2020 Sep 9 (Durinck et al., 2005).

We also tested whether gene families that shifted in expansion and contraction rate along branches leading to giant taxa were enriched for cancer genes. Six branches were tested in this set of focal branches relative to all other branches in the vertebrate phylogeny: the branch leading to whale shark, the branch leading to ocean sunfish, the branch leading to African elephant, and the branches corresponding to the clade of baleen whales (the clade sister to bottlenose dolphin). To confirm whether the results were more extreme than expected, we performed two tests. First, we drew the branches corresponding to the non-giant sister taxa of the vertebrate giants, then tested for whether these were enriched for cancer genes. Secondly, we tested for cancer gene enrichment on 100 permutations of selecting six random branches without replacement from across the vertebrate phylogeny. We then compared the observed odds ratio of enrichment for cancer genes to this null distribution.

We then compared the rate of gene family size evolution for gene families related to cancer to rates of gene families not related to cancer along branches leading to giant vertebrates and the remaining branches in phylogeny. Branch-wise rates of gene family size evolution were estimated by computing the difference in estimated ancestral and descendant gene family sizes of each branch and dividing by time. We also used the lme4 package and lmerTest packages to fit a linear mixed model and test for significant contribution on rate depends on whether it was estimated for a cancer gene or not, whether the rate was estimated on a branch leading to a giant taxon or not, the interaction of these variables, and with gene family as a random effect.

---

**Figure 2–figure supplement 1.**
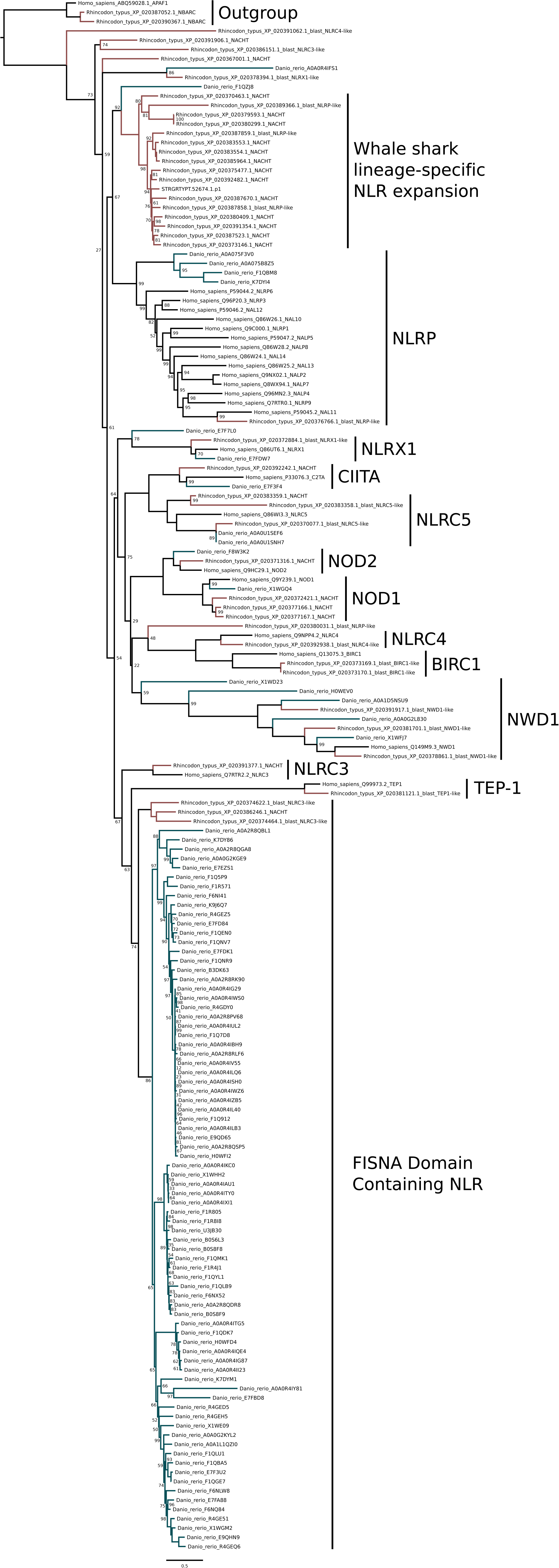
Phylogenetic analysis of NLRs from whale shark, zebrafish and human. Branches leading to human sequences are shown in black, to zebrafish in blue-green, and whale shark in red. Whale shark sequences with a detectable NACHT have ‘_NACHT’ at the end of the sequence identifier (except for transcriptome sequence which all contain NACHT domains; All NACHT domain containing sequences are also noted with ‘*’ in table 1).

**Figure 2–figure supplement 2.**
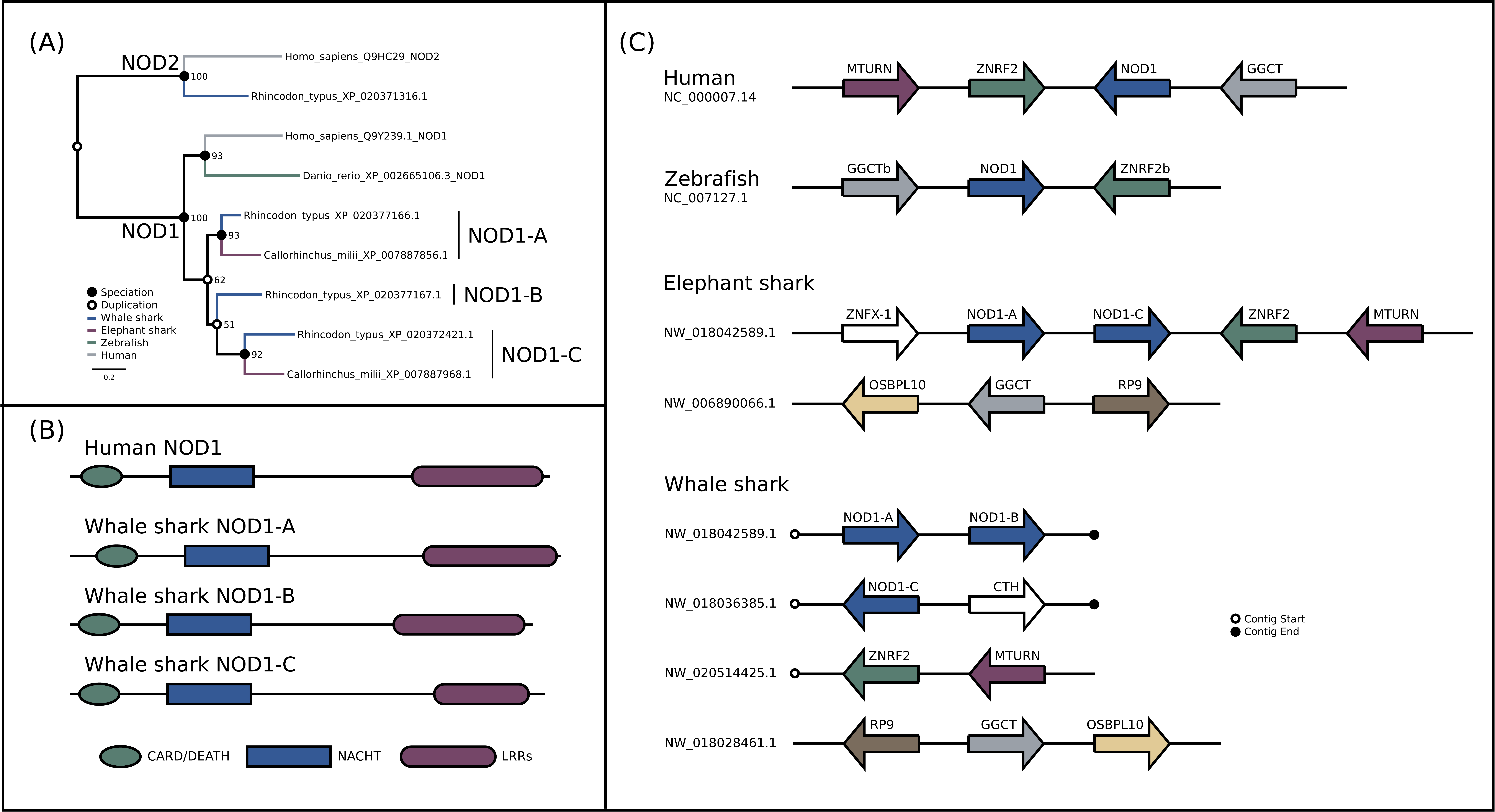
Detailed analysis of NOD1 evolution. (A) Phylogenetic analysis of NOD1 focused NLR dataset. (B) Domain structure of NOD1 duplicates in whale shark. (C) Synteny analysis of jawed vertebrate NOD1s.

**Figure 2–figure supplement 3.**
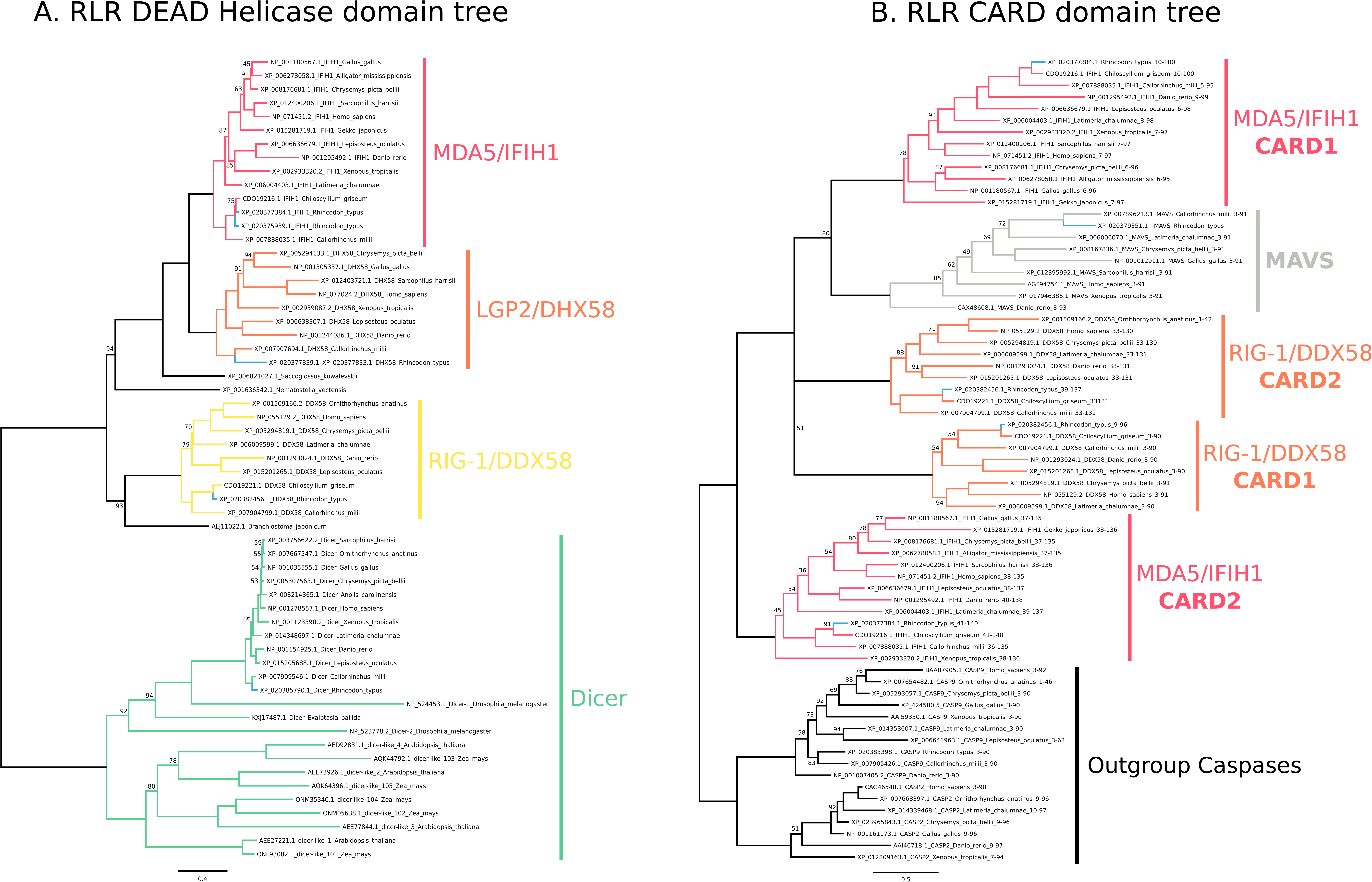
Phylogenetic Analyses of whale shark and jawed vertebrate RLRs, DICER and MAVS. (A) Tree of the RLR and DICER DEAD-Helicase domains. (B) Tree of the RLR and MAVS CARD domains. Whale shark sequences are highlighted in blue.

**Figure 2–figure supplement 4.**
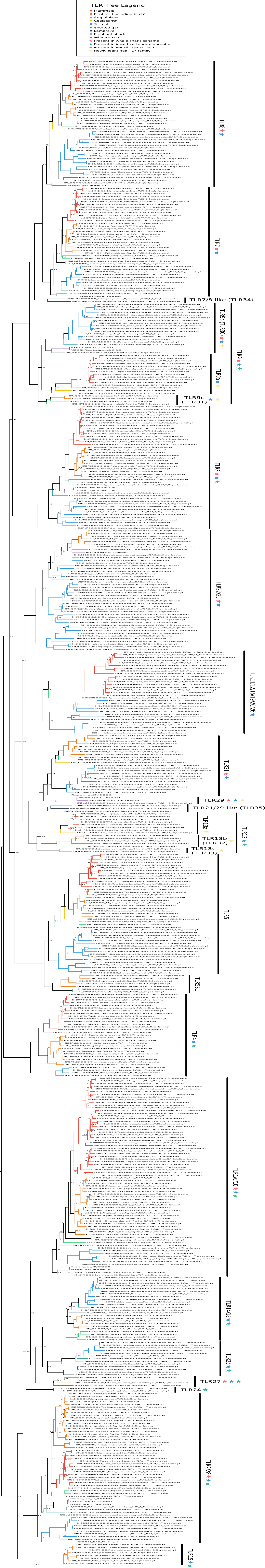
Phylogenetic tree of vertebrate TLRs, including new whale shark sequences. The tree is rooted according to the minimal ancestor deviation method (Kümmel Tria, Landan and Dagan, 2017).

## Notes

### Competing Interest Statement

The authors have declared no competing interest.

### Summary of Updates

The version of the manuscript has been revised to incorporate analysis of heterozygosity, further analysis of cancer-related gene evolution relative to gigantism, increased clarity of introduction and discussion of immune genes and other aspects of the research, and incorporation of some of the supplementary text into the main text.

## References

1. Abegglen LM, Caulin AF, Chan A, Lee K, Robinson R, Campbell MS, Kiso WK, Schmitt DL, Waddell PJ, Bhaskara S, Jensen ST, Maley CC, Schiffman JD. 2015. Potential Mechanisms for Cancer Resistance in Elephants and Comparative Cellular Response to DNA Damage in Humans. JAMA 314:1850–1860.

2. Alexopoulos H, Böttger A, Fischer S, Levin A, Wolf A, Fujisawa T, Hayakawa S, Gojobori T, Davies JA, David CN, Bacon JP. 2004. Evolution of gap junctions: the missing link? Curr Biol 14:R879–80.

3. Altschul SF, Gish W, Miller W, Myers EW, Lipman DJ. 1990. Basic local alignment search tool. J Mol Biol 215:403–410.

4. Ballesteros JA, Hormiga G. 2016. A New Orthology Assessment Method for Phylogenomic Data: Unrooted Phylogenetic Orthology. Mol Biol Evol 33:2481.

5. Benton MJ, Donoghue PCJ, Asher RJ, Friedman M, Near TJ, Vinther J. 2014. Constraints on the timescale of animal evolutionary history. Palaentologica Electronica 18.1.1FC:1–106.

6. Boeckmann B, Bairoch A, Apweiler R, Blatter M-C, Estreicher A, Gasteiger E, Martin MJ, Michoud K, O’Donovan C, Phan I, Pilbout S, Schneider M. 2003. The SWISS-PROT protein knowledgebase and its supplement TrEMBL in 2003. Nucleic Acids Res 31:365–370.

7. Boehm T. 2012. Evolution of vertebrate immunity. Curr Biol 22:R722–32.

8. Bolger AM, Lohse M, Usadel B. 2014. Trimmomatic: a flexible trimmer for Illumina sequence data. Bioinformatics 30:2114–2120.

9. Boot-Handford RP, Tuckwell DS. 2003. Fibrillar collagen: the key to vertebrate evolution? A tale of molecular incest. Bioessays 25:142–151.

10. Boudinot P, Zou J, Ota T, Buonocore F, Scapigliati G, Canapa A, Cannon J, Litman G, Hansen JD. 2014. A tetrapod-like repertoire of innate immune receptors and effectors for coelacanths. J Exp Zool B Mol Dev Evol 322:415–437.

11. Braasch I, Gehrke AR, Smith JJ, Kawasaki K, Manousaki T, Pasquier J, Amores A, Desvignes T, Batzel P, Catchen J, Berlin AM, Campbell MS, Barrell D, Martin KJ, Mulley JF, Ravi V, Lee AP, Nakamura T, Chalopin D, Fan S, Wcisel D, Cañestro C, Sydes J, Beaudry FEG, Sun Y, Hertel J, Beam MJ, Fasold M, Ishiyama M, Johnson J, Kehr S, Lara M, Letaw JH, Litman GW, Litman RT, Mikami M, Ota T, Saha NR, Williams L, Stadler PF, Wang H, Taylor JS, Fontenot Q, Ferrara A, Searle SMJ, Aken B, Yandell M, Schneider I, Yoder JA, Volff J-N, Meyer A, Amemiya CT, Venkatesh B, Holland PWH, Guiguen Y, Bobe J, Shubin NH, Di Palma F, Alföldi J, Lindblad-Toh K, Postlethwait JH. 2016. The spotted gar genome illuminates vertebrate evolution and facilitates human-teleost comparisons. Nat Genet 48:427–437.

12. Brayer KJ, Segal DJ. 2008. Keep your fingers off my DNA: protein--protein interactions mediated by C2H2 zinc finger domains. Cell Biochem Biophys 50:111–131.

13. Brown GD, Willment JA, Whitehead L. 2018. C-type lectins in immunity and homeostasis. Nat Rev Immunol 18:374–389.

14. Brown JW, Walker JF, Smith SA. 2017. Phyx: phylogenetic tools for unix. Bioinformatics 33:1886–1888.

15. Buckley KM, Rast JP. 2015. Diversity of animal immune receptors and the origins of recognition complexity in the deuterostomes. Dev Comp Immunol 49:179–189.

16. Capella-Gutierrez S, Silla-Martinez JM, Gabaldon T. 2009. trimAl: a tool for automated alignment trimming in large-scale phylogenetic analyses. Bioinformatics 25:1972–1973.

17. Chen H, Liang Y, Han Y, Liu T, Chen S. 2021. Genome-wide analysis of Toll-like receptors in zebrafish and the effect of rearing temperature on the receptors in response to stimulated pathogen infection. J Fish Dis 44:337–349.

18. Coates MI, Gess RW, Finarelli JA, Criswell KE, Tietjen K. 2017. A symmoriiform chondrichthyan braincase and the origin of chimaeroid fishes. Nature 541:208–211.

19. Criscuolo A, Gribaldo S. 2010. BMGE (Block Mapping and Gathering with Entropy): a new software for selection of phylogenetic informative regions from multiple sequence alignments. BMC Evol Biol 10:210.

20. Dooley H. 2014. Athena and the Evolution of Adaptive Immunity. Immunobiology of the Shark 29.

21. Durinck S, Moreau Y, Kasprzyk A, Davis S, De Moor B, Brazma A, Huber W. 2005. BioMart and Bioconductor: a powerful link between biological databases and microarray data analysis. Bioinformatics 21:3439–3440.

22. Eddy SR. 1998. Profile hidden Markov models. Bioinformatics 14:755–763.

23. Emms DM, Kelly S. 2015. OrthoFinder: solving fundamental biases in whole genome comparisons dramatically improves orthogroup inference accuracy. Genome Biol 16:157.

24. Esselstyn JA, Oliveros CH, Swanson MT, Faircloth BC. 2017. Investigating Difficult Nodes in the Placental Mammal Tree with Expanded Taxon Sampling and Thousands of Ultraconserved Elements. Genome Biol Evol 9:2308–2321.

25. Flajnik MF, Kasahara M. 2010. Origin and evolution of the adaptive immune system: genetic events and selective pressures. Nat Rev Genet 11:47–59.

26. Fritz JH, Kufer TA. 2015. Editorial: NLR-Protein Functions in Immunity. Front Immunol 6:306.

27. Fu L, Niu B, Zhu Z, Wu S, Li W. 2012. CD-HIT: accelerated for clustering the next-generation sequencing data. Bioinformatics 28:3150–3152.

28. Garrison E, Kronenberg ZN, Dawson ET, Pedersen BS, Prins P. 2021. Vcflib and tools for processing the VCF variant call format. bioRxiv. doi:10.1101/2021.05.21.445151

29. Garrison E, Marth G. 2012. Haplotype-based variant detection from short-read sequencing. arXiv *[q-bioGN]*.

30. Gurevich A, Saveliev V, Vyahhi N, Tesler G. 2013. QUAST: quality assessment tool for genome assemblies. Bioinformatics 29:1072–1075.

31. Han MV, Thomas GWC, Lugo-Martinez J, Hahn MW. 2013. Estimating gene gain and loss rates in the presence of error in genome assembly and annotation using CAFE 3. Mol Biol Evol 30:1987–1997.

32. Haq F, Ahmed N, Qasim M. 2019. Comparative genomic analysis of collagen gene diversity. 3 Biotech 9:83.

33. Hara Y, Tatsumi K, Yoshida M, Kajikawa E, Kiyonari H, Kuraku S. 2015. Optimizing and benchmarking de novo transcriptome sequencing: from library preparation to assembly evaluation. BMC Genomics 16:977.

34. Hara Y, Yamaguchi K, Onimaru K, Kadota M, Koyanagi M, Keeley SD, Tatsumi K, Tanaka K, Motone F, Kageyama Y, Nozu R, Adachi N, Nishimura O, Nakagawa R, Tanegashima C, Kiyatake I, Matsumoto R, Murakumo K, Nishida K, Terakita A, Kuratani S, Sato K, Hyodo S, Kuraku S. 2018. Shark genomes provide insights into elasmobranch evolution and the origin of vertebrates. Nat Ecol Evol 2:1761–1771.

35. Hoang DT, Chernomor O, von Haeseler A, Minh BQ, Le SV. 2017. UFBoot2: Improving the Ultrafast Bootstrap Approximation. Mol Biol Evol. doi:10.1093/molbev/msx281

36. Howe K, Schiffer PH, Zielinski J, Wiehe T, Laird GK, Marioni JC, Soylemez O, Kondrashov F, Leptin M. 2016. Structure and evolutionary history of a large family of NLR proteins in the zebrafish. Open Biol 6:160009.

37. Huang S, Yuan S, Guo L, Yu Y, Li J, Wu T, Liu T, Yang M, Wu K, Liu H, Ge J, Yu Y, Huang H, Dong M, Yu C, Chen S, Xu A. 2008. Genomic analysis of the immune gene repertoire of amphioxus reveals extraordinary innate complexity and diversity. Genome Res 18:1112– 1126.

38. Hunter S, Apweiler R, Attwood TK, Bairoch A, Bateman A, Binns D, Bork P, Das U, Daugherty L, Duquenne L, Finn RD, Gough J, Haft D, Hulo N, Kahn D, Kelly E, Laugraud A, Letunic I, Lonsdale D, Lopez R, Madera M, Maslen J, McAnulla C, McDowall J, Mistry J, Mitchell A, Mulder N, Natale D, Orengo C, Quinn AF, Selengut JD, Sigrist CJA, Thimma M, Thomas PD, Valentin F, Wilson D, Wu CH, Yeats C. 2009. InterPro: the integrative protein signature database. Nucleic Acids Res 37:D211–5.

39. Jones P, Binns D, Chang H-Y, Fraser M, Li W, McAnulla C, McWilliam H, Maslen J, Mitchell A, Nuka G, Pesseat S, Quinn AF, Sangrador-Vegas A, Scheremetjew M, Yong S-Y, Lopez R, Hunter S. 2014. InterProScan 5: genome-scale protein function classification. Bioinformatics 30:1236–1240.

40. Kaiser P, Rothwell L, Avery S, Balu S. 2004. Evolution of the interleukins. Dev Comp Immunol 28:375–394.

41. Kajitani R, Toshimoto K, Noguchi H, Toyoda A, Ogura Y, Okuno M, Yabana M, Harada M, Nagayasu E, Maruyama H, Kohara Y, Fujiyama A, Hayashi T, Itoh T. 2014. Efficient de novo assembly of highly heterozygous genomes from whole-genome shotgun short reads. Genome Res 24:1384–1395.

42. Kalyaanamoorthy S, Minh BQ, Wong TKF, von Haeseler A, Jermiin LS. 2017. ModelFinder: fast model selection for accurate phylogenetic estimates. Nat Methods 14:587–589.

43. Kasamatsu J, Oshiumi H, Matsumoto M, Kasahara M, Seya T. 2010. Phylogenetic and expression analysis of lamprey toll-like receptors. Dev Comp Immunol 34:855–865.

44. Katoh K, Standley DM. 2013. MAFFT multiple sequence alignment software version 7: improvements in performance and usability. Mol Biol Evol 30:772–780.

45. Kelley J, Walter L, Trowsdale J. 2005. Comparative genomics of natural killer cell receptor gene clusters. PLoS Genet 1:129–139.

46. Koren S, Walenz BP, Berlin K, Miller JR, Bergman NH, Phillippy AM. 2017. Canu: scalable and accurate long-read assembly via adaptive k-mer weighting and repeat separation. Genome Res. doi:10.1101/gr.215087.116

47. Korithoski B, Kolaczkowski O, Mukherjee K, Kola R, Earl C, Kolaczkowski B. 2015. Evolution of a Novel Antiviral Immune-Signaling Interaction by Partial-Gene Duplication. PLoS One 10:e0137276.

48. Krishnaswamy Gopalan T, Gururaj P, Gupta R, Gopal DR, Rajesh P, Chidambaram B, Kalyanasundaram A, Angamuthu R. 2014. Transcriptome profiling reveals higher vertebrate orthologous of intra-cytoplasmic pattern recognition receptors in grey bamboo shark. PLoS One 9:e100018.

49. Laetsch DR, Blaxter ML. 2017. KinFin: Software for Taxon-Aware Analysis of Clustered Protein Sequences. G3 7:3349–3357.

50. Li H. 2013. Aligning sequence reads, clone sequences and assembly contigs with BWA-MEM. arXiv preprint arXiv:13033997.

51. Li J, Mahajan A, Tsai M-D. 2006. Ankyrin repeat: a unique motif mediating protein- protein interactions. Biochemistry 45:15168–15178.

52. Litscher ES, Wassarman PM. 2014. Evolution, structure, and synthesis of vertebrate egg-coat proteins. Trends Dev Biol 8:65–76.

53. Loo Y-M, Gale M Jr. 2011. Immune signaling by RIG-I-like receptors. Immunity 34:680–692.

54. Mapleson D, Garcia Accinelli G, Kettleborough G, Wright J, Clavijo BJ. 2016. KAT: a K-mer analysis toolkit to quality control NGS datasets and genome assemblies. Bioinformatics. doi:10.1093/bioinformatics/btw663

55. Marçais G, Kingsford C. 2011. A fast, lock-free approach for efficient parallel counting of occurrences of k-mers. Bioinformatics 27:764–770.

56. Marchler-Bauer A, Derbyshire MK, Gonzales NR, Lu S, Chitsaz F, Geer LY, Geer RC, He J, Gwadz M, Hurwitz DI, Lanczycki CJ, Lu F, Marchler GH, Song JS, Thanki N, Wang Z, Yamashita RA, Zhang D, Zheng C, Bryant SH. 2015. CDD: NCBI’s conserved domain database. Nucleic Acids Res 43:D222–6.

57. Margolin JF, Friedman JR, Meyer WK, Vissing H, Thiesen HJ, Rauscher FJ 3rd. 1994. Krüppel-associated boxes are potent transcriptional repression domains. Proc Natl Acad Sci U S A 91:4509–4513.

58. Marra NJ, Stanhope MJ, Jue NK, Wang M, Sun Q, Bitar PP, Richards VP, Komissarov A, Rayko M, Kliver S, Stanhope BJ, Winkler C, O’Brien SJ, Antunes A, Jorgensen S, Shivji MS. 2019. White shark genome reveals ancient elasmobranch adaptations associated with wound healing and the maintenance of genome stability. Proc Natl Acad Sci U S A 201819778.

59. Martin AP, Palumbi SR. 1993. Body size, metabolic rate, generation time, and the molecular clock. Proc Natl Acad Sci U S A 90:4087–4091.

60. McClain CR, Balk MA, Benfield MC, Branch TA, Chen C, Cosgrove J, Dove ADM, Gaskins LC, Helm RR, Hochberg FG, Lee FB, Marshall A, McMurray SE, Schanche C, Stone SN, Thaler AD. 2015. Sizing ocean giants: patterns of intraspecific size variation in marine megafauna. PeerJ 2:e715.

61. Minh BQ, Nguyen MAT, von Haeseler A. 2013. Ultrafast approximation for phylogenetic bootstrap. Mol Biol Evol 30:1188–1195.

62. Misof B, Meyer B, von Reumont BM, Kück P, Misof K, Meusemann K. 2013. Selecting informative subsets of sparse supermatrices increases the chance to find correct trees. BMC Bioinformatics 14:348.

63. Mukherjee K, Korithoski B, Kolaczkowski B. 2014. Ancient origins of vertebrate-specific innate antiviral immunity. Mol Biol Evol 31:140–153.

64. Near TJ, Meylan PA, Shaffer HB. 2005. Assessing concordance of fossil calibration points in molecular clock studies: an example using turtles. Am Nat 165:137–146.

65. Nguyen L-T, Schmidt HA, von Haeseler A, Minh BQ. 2015. IQ-TREE: a fast and effective stochastic algorithm for estimating maximum-likelihood phylogenies. Mol Biol Evol 32:268– 274.

66. Nishimura O, Hara Y, Kuraku S. 2017. gVolante for standardizing completeness assessment of genome and transcriptome assemblies. Bioinformatics 33:3635–3637.

67. O’Connell J, Schulz-Trieglaff O, Carlson E, Hims MM, Gormley NA, Cox AJ. 2015. NxTrim: optimized trimming of Illumina mate pair reads. Bioinformatics 31:2035–2037.

68. Ohno S, Wolf U, Atkin NB. 1968. Evolution from fish to mammals by gene duplication. Hereditas 59:169–187.

69. O’Leary NA, Wright MW, Brister JR, Ciufo S, Haddad D, McVeigh R, Rajput B, Robbertse B, Smith-White B, Ako-Adjei D, Astashyn A, Badretdin A, Bao Y, Blinkova O, Brover V, Chetvernin V, Choi J, Cox E, Ermolaeva O, Farrell CM, Goldfarb T, Gupta T, Haft D, Hatcher E, Hlavina W, Joardar VS, Kodali VK, Li W, Maglott D, Masterson P, McGarvey KM, Murphy MR, O’Neill K, Pujar S, Rangwala SH, Rausch D, Riddick LD, Schoch C, Shkeda A, Storz SS, Sun H, Thibaud-Nissen F, Tolstoy I, Tully RE, Vatsan AR, Wallin C, Webb D, Wu W, Landrum MJ, Kimchi A, Tatusova T, DiCuccio M, Kitts P, Murphy TD, Pruitt KD. 2016. Reference sequence (RefSeq) database at NCBI: current status, taxonomic expansion, and functional annotation. Nucleic Acids Res 44:D733–45.

70. Peto R, Roe FJ, Lee PN, Levy L, Clack J. 1975. Cancer and ageing in mice and men. Br J Cancer 32:411–426.

71. Pierce KL, Premont RT, Lefkowitz RJ. 2002. Seven-transmembrane receptors. Nat Rev Mol Cell Biol 3:639–650.

72. Pimiento C, Cantalpiedra JL, Shimada K, Field DJ, Smaers JB. 2019. Evolutionary pathways toward gigantism in sharks and rays. Evolution. doi:10.1111/evo.13680

73. Popper AN, Platt C, Edds PL. 1992. Evolution of the Vertebrate Inner Ear: An Overview of Ideas In: Webster DB, Popper AN, Fay RR, editors. The Evolutionary Biology of Hearing. New York, NY: Springer New York. pp. 49–57.

74. Puttick MN, Thomas GH. 2015. Fossils and living taxa agree on patterns of body mass evolution: a case study with Afrotheria. Proc Biol Sci 282:20152023.

75. Rabosky DL, Santini F, Eastman J, Smith SA, Sidlauskas B, Chang J, Alfaro ME. 2013. Rates of speciation and morphological evolution are correlated across the largest vertebrate radiation. Nat Commun 4:1958.

76. Rast JP, Smith LC, Loza-Coll M, Hibino T, Litman GW. 2006. Genomic insights into the immune system of the sea urchin. Science 314:952–956.

77. Read TD, Petit RA 3rd, Joseph SJ, Alam MT, Weil MR, Ahmad M, Bhimani R, Vuong JS, Haase CP, Webb DH, Tan M, Dove ADM. 2017. Draft sequencing and assembly of the genome of the world’s largest fish, the whale shark: *Rhincodon typus* Smith 1828. BMC Genomics 18:532.

78. Redmond AK, Macqueen DJ, Dooley H. 2018. Phylotranscriptomics suggests the jawed vertebrate ancestor could generate diverse helper and regulatory T cell subsets. BMC Evol Biol 18:169.

79. Rhincodon typus Annotation Report. 2018. https://www.ncbi.nlm.nih.gov/genome/annotation_euk/Rhincodon_typus/100/

80. Richter DJ, Fozouni P, Eisen MB, King N. 2018. Gene family innovation, conservation and loss on the animal stem lineage. Elife 7. doi:10.7554/eLife.34226

81. Roach JC, Glusman G, Rowen L, Kaur A, Purcell MK, Smith KD, Hood LE, Aderem A. 2005. The evolution of vertebrate Toll-like receptors. Proc Natl Acad Sci U S A 102:9577–9582.

82. Schroder K, Tschopp J. 2010. The inflammasomes. Cell 140:821–832.

83. Shen W, Le S, Li Y, Hu F. 2016. SeqKit: A Cross-Platform and Ultrafast Toolkit for FASTA/Q File Manipulation. PLoS One 11:e0163962.

84. She R, Chu JS-C, Uyar B, Wang J, Wang K, Chen N. 2011. genBlastG: using BLAST searches to build homologous gene models. Bioinformatics 27:2141–2143.

85. Simão FA, Waterhouse RM, Ioannidis P, Kriventseva EV, Zdobnov EM. 2015. BUSCO: assessing genome assembly and annotation completeness with single-copy orthologs. Bioinformatics 31:3210–3212.

86. Singh PP, Arora J, Isambert H. 2015. Identification of Ohnolog Genes Originating from Whole Genome Duplication in Early Vertebrates, Based on Synteny Comparison across Multiple Genomes. PLoS Comput Biol 11:e1004394.

87. Slater GJ, Goldbogen JA, Pyenson ND. 2017. Independent evolution of baleen whale gigantism linked to Plio-Pleistocene ocean dynamics. Proc Biol Sci 284. doi:10.1098/rspb.2017.0546

88. Smith JJ, Kuraku S, Holt C, Sauka-Spengler T, Jiang N, Campbell MS, Yandell MD, Manousaki T, Meyer A, Bloom OE, Morgan JR, Buxbaum JD, Sachidanandam R, Sims C, Garruss AS, Cook M, Krumlauf R, Wiedemann LM, Sower SA, Decatur WA, Hall JA, Amemiya CT, Saha NR, Buckley KM, Rast JP, Das S, Hirano M, McCurley N, Guo P, Rohner N, Tabin CJ, Piccinelli P, Elgar G, Ruffier M, Aken BL, Searle SMJ, Muffato M, Pignatelli M, Herrero J, Jones M, Brown CT, Chung-Davidson Y-W, Nanlohy KG, Libants SV, Yeh C-Y, McCauley DW, Langeland JA, Pancer Z, Fritzsch B, de Jong PJ, Zhu B, Fulton LL, Theising B, Flicek P, Bronner ME, Warren WC, Clifton SW, Wilson RK, Li W. 2013. Sequencing of the sea lamprey (Petromyzon marinus) genome provides insights into vertebrate evolution. Nat Genet 45:415–21, 421e1–2.

89. Solbakken MH, Voje KL, Jakobsen KS, Jentoft S. 2017. Linking species habitat and past palaeoclimatic events to evolution of the teleost innate immune system. Proc Biol Sci 284. doi:10.1098/rspb.2016.2810

90. Sondka Z, Bamford S, Cole CG, Ward SA, Dunham I, Forbes SA. 2018. The COSMIC Cancer Gene Census: describing genetic dysfunction across all human cancers. Nat Rev Cancer 18:696–705.

91. Stein C, Caccamo M, Laird G, Leptin M. 2007. Conservation and divergence of gene families encoding components of innate immune response systems in zebrafish. Genome Biol 8:R251.

92. Sulak M, Fong L, Mika K, Chigurupati S, Yon L, Mongan NP, Emes RD, Lynch VJ. 2016. TP53 copy number expansion is associated with the evolution of increased body size and an enhanced DNA damage response in elephants. Elife 5. doi:10.7554/eLife.11994

93. Takei Y, Hasegawa Y, Watanabe TX, Nakajima K, Hazon N. 1993. A novel angiotensin I isolated from an elasmobranch fish. J Endocrinol 139:281–285.

94. Takezaki N, Rzhetsky A, Nei M. 1995. Phylogenetic test of the molecular clock and linearized trees. Mol Biol Evol 12:823–833.

95. Tassia MG, Whelan NV, Halanych KM. 2017. Toll-like receptor pathway evolution in deuterostomes. Proc Natl Acad Sci U S A 114:7055–7060.

96. Tollis M, Robbins J, Webb AE, Kuderna LFK, Caulin AF, Garcia JD, Bèrubè M, Pourmand N, Marques-Bonet T, O’Connell MJ, Palsbøll PJ, Maley CC. 2019. Return to the sea, get huge, beat cancer: an analysis of cetacean genomes including an assembly for the humpback whale (Megaptera novaeangliae). Mol Biol Evol. doi:10.1093/molbev/msz099

97. Tollis M, Schiffman JD, Boddy AM. 2017. Evolution of cancer suppression as revealed by mammalian comparative genomics. Curr Opin Genet Dev 42:40–47.

98. Tria FDK, Landan G, Dagan T. 2017. Phylogenetic rooting using minimal ancestor deviation. Nat Ecol Evol 1:193.

99. Urbach JM, Ausubel FM. 2017. The NBS-LRR architectures of plant R-proteins and metazoan NLRs evolved in independent events. Proc Natl Acad Sci U S A 114:1063–1068.

100. Venkatesh B, Lee AP, Ravi V, Maurya AK, Lian MM, Swann JB, Ohta Y, Flajnik MF, Sutoh Y, Kasahara M, Hoon S, Gangu V, Roy SW, Irimia M, Korzh V, Kondrychyn I, Lim ZW, Tay B- H, Tohari S, Kong KW, Ho S, Lorente-Galdos B, Quilez J, Marques-Bonet T, Raney BJ, Ingham PW, Tay A, Hillier LW, Minx P, Boehm T, Wilson RK, Brenner S, Warren WC. 2014. Elephant shark genome provides unique insights into gnathostome evolution. Nature 505:174–179.

101. Vurture GW, Sedlazeck FJ, Nattestad M, Underwood CJ, Fang H, Gurtowski J, Schatz MC. 2017. GenomeScope: fast reference-free genome profiling from short reads. Bioinformatics 33:2202–2204.

102. Wada H, Okuyama M, Satoh N, Zhang S. 2006. Molecular evolution of fibrillar collagen in chordates, with implications for the evolution of vertebrate skeletons and chordate phylogeny. Evol Dev 8:370–377.

103. Walker BJ, Abeel T, Shea T, Priest M, Abouelliel A, Sakthikumar S, Cuomo CA, Zeng Q, Wortman J, Young SK, Earl AM. 2014. Pilon: an integrated tool for comprehensive microbial variant detection and genome assembly improvement. PLoS One 9:e112963.

104. Wang J, Zhang Z, Liu J, Zhao J, Yin D. 2016. Ectodomain Architecture Affects Sequence and Functional Evolution of Vertebrate Toll-like Receptors. Sci Rep 6:26705.

105. Warren WC, Hillier LW, Tomlinson C, Minx P, Kremitzki M, Graves T, Markovic C, Bouk N, Pruitt KD, Thibaud-Nissen F, Schneider V, Mansour TA, Brown CT, Zimin A, Hawken R, Abrahamsen M, Pyrkosz AB, Morisson M, Fillon V, Vignal A, Chow W, Howe K, Fulton JE, Miller MM, Lovell P, Mello CV, Wirthlin M, Mason AS, Kuo R, Burt DW, Dodgson JB, Cheng HH. 2017. A New Chicken Genome Assembly Provides Insight into Avian Genome Structure. G3 7:109–117.

106. Weber JA, Park SG, Luria V, Jeon S, Kim H-M, Jeon Y, Bhak Y, Jun JH, Kim SW, Hong WH, Lee S, Cho YS, Karger A, Cain JW, Manica A, Kim S, Kim J-H, Edwards JS, Bhak J, Church GM. 2020. The whale shark genome reveals how genomic and physiological properties scale with body size. Proc Natl Acad Sci U S A 117:20662–20671.

107. Wolfe SA, Nekludova L, Pabo CO. 2000. DNA recognition by Cys2His2 zinc finger proteins. Annu Rev Biophys Biomol Struct 29:183–212.

108. Wright SP. 1992. Adjusted p-values for simultaneous inference. Biometrics 48:1005.

109. Yang Z. 2007. PAML 4: Phylogenetic Analysis by Maximum Likelihood. Mol Biol Evol 24:1586– 1591.

110. Yates AD, Achuthan P, Akanni W, Allen J, Allen J, Alvarez-Jarreta J, Amode MR, Armean IM, Azov AG, Bennett R, Bhai J, Billis K, Boddu S, Marugán JC, Cummins C, Davidson C, Dodiya K, Fatima R, Gall A, Giron CG, Gil L, Grego T, Haggerty L, Haskell E, Hourlier T, Izuogu OG, Janacek SH, Juettemann T, Kay M, Lavidas I, Le T, Lemos D, Martinez JG, Maurel T, McDowall M, McMahon A, Mohanan S, Moore B, Nuhn M, Oheh DN, Parker A, Parton A, Patricio M, Sakthivel MP, Abdul Salam AI, Schmitt BM, Schuilenburg H, Sheppard D, Sycheva M, Szuba M, Taylor K, Thormann A, Threadgold G, Vullo A, Walts B, Winterbottom A, Zadissa A, Chakiachvili M, Flint B, Frankish A, Hunt SE, IIsley G, Kostadima M, Langridge N, Loveland JE, Martin FJ, Morales J, Mudge JM, Muffato M, Perry E, Ruffier M, Trevanion SJ, Cunningham F, Howe KL, Zerbino DR, Flicek P. 2020. Ensembl 2020. Nucleic Acids Res 48:D682–D688.

111. Zhang C, Rabiee M, Sayyari E, Mirarab S. 2018. ASTRAL-III: polynomial time species tree reconstruction from partially resolved gene trees. BMC Bioinformatics 19:153.

112. Zhang Y, Gao H, Li H, Guo J, Ouyang B, Wang M, Xu Q, Wang J, Lv M, Guo X, Liu Q, Wei L, Ren H, Xi Y, Guo Y, Ren B, Pan S, Liu C, Ding X, Xiang H, Yu Y, Song Y, Meng L, Liu S, Wang J, Jiang Y, Shi J, Liu S, Sabir JSM, Sabir MJ, Khan M, Hajrah NH, Ming-Yuen Lee S, Xu X, Yang H, Wang J, Fan G, Yang N, Liu X. 2020. The White-Spotted Bamboo Shark Genome Reveals Chromosome Rearrangements and Fast-Evolving Immune Genes of Cartilaginous Fish. iScience 23:101754.

