## Appendix for "The whale shark genome reveals patterns of vertebrate gene family evolution"

Appendix 1. Genome content

Appendix 1A. Transposable Element Content.

The whale shark genome is relatively large compared to most (but not all) other fish genomes that have been sequenced. The larger genome size relative to *Callorhinchus* was likely driven in part by increases in content of repetitive elements such as transposable elements, as typical in eukaryotes (Gregory, 2005). Repetitive elements (Supplementary File 1) were annotated using the repeat library construction scripts used in MAKER-P (Campbell et al., 2014; Jiang et al., 2016). This pipeline includes using MITE-Hunter (version 11-2011) to identify MITEs (Han and Wessler, 2010), LTRharvest/LTRdigest (both installed through genometools version 1.5.8) to identify 99% similar and 85% similar LTRs (Ellinghaus et al., 2008; Steinbiss et al., 2009), and RepeatModeler v4.05 to identify other putative repeats (Smit and Hubley, 2008). To further classify repeats not initially identified by RepeatModeler, we used RepeatClassifier (which comes with RepeatModeler), and then searched the remaining unclassified repeats against the Dfam database, 2016 November 7 release downloaded 2017 Feb 27 (Hubley et al., 2016) using nhmmer (HMMER v3.1b2) with default settings (Hubley et al., 2016). We annotated that the proportion of repetitive elements to the genome length was 50.34% of the genome assembly (Supplementary table 2), only slightly higher than previous estimates: Hara et al. (Hara et al., 2018) annotated the proportion of repetitive elements as 46.63% of the genome, and Weber et al. (2020) annotated the proportion as 49.55% of the genome.

Based on annotation of repetitive elements, we found a much larger proportion of the whale shark genome (>50%) consists of transposable elements compared to *Callorhinchus* genome (28%) (Supplementary table 2). This is similar to the proportion found in zebrafish and higher than in human, which have genomes of roughly the same size (Braasch et al., 2016; Howe et al., 2013). This is also similar to the proportion of repetitive elements previously reported (Hara et al., 2018; Weber et al., 2020). By contrast, the assembly-free approach based on *k*-mers by GenomeScope estimated only approximately 789 Mbp repeat length (28%), which is likely an under-estimate. Given the draft nature of our assembly, we may still underestimate the proportion of the genome comprised of repetitive elements (Chalopin et al., 2015). A large proportion of repetitive elements was also found in the white shark genome (Marra et al., 2019).

Though the relationship between overall proportion of transposable elements and the whale shark genome size is consistent with patterns found in other vertebrates, proportions of transposable element classes differed (Supplementary table 2). The most well-represented class of repeats of DNA transposons, LINE, SINE, and LTR) in the whale shark genome were LINEs (33.25% of the genome), with the most well-represented superfamily of repeats being the CR1 LINEs (21.78% of the genome). CR1 LINEs comprise almost all the transposable element content in the whale shark genome (84.22% of genome content covered by transposable elements), which is high compared to the other vertebrate genomes considered. Compared to *Callorhinchus*, the whale shark has a greater proportion of LINEs (33.25% vs. 12.6%), which is primarily a difference in the proportion of CR1 repeats (21.78% vs. 4.0%) (Venkatesh et al., 2014), which has previously been noted to be in high proportion in the whale shark genome

(Marra et al., 2019; Weber et al., 2020). In addition, *Callorhinchus* has a much higher proportion of SINEs than whale shark (13.1% vs. 1.55%) (Venkatesh et al., 2014). The whale shark also has a higher proportion of LINEs and CR1s relative to the white shark (33.25% vs. 29.84%; 21.78% vs 18.75%). The previous assembly of the whale shark had a lower proportion of LINEs and CR1s than the white shark genome (Marra et al., 2019), and the increase is likely in part due to the use of long reads in the new assembly. The whale shark also has a relatively high proportion of unclassified repeats among vertebrates (10.8%, most vertebrates <5%) (Chalopin et al., 2015). Previous research also identified the large proportion of the genome comprising CR1 LINEs, particularly in introns (Weber et al., 2020).

The whale shark genome contains many of the transposable element superfamilies that are widespread in vertebrates, including the DNA transposons TcMariner, hAT, PIF-Harbinger, and Helitron, and many RNA transposons including the LTRs Gypsy, Copia, endogenous retroviruses, and the LINEs Penelope, RTE, CR1, and LINE2 (Supplementary File 1). We did not detect some transposable elements found in *Callorhinchus* including the DNA transposons PiggyBac, Sola, and Crypton and the RNA transposons Dong, and R2. By contrast, we found potential sequences of a number of transposons not found in *Callorhinchus*, including the DNA transposon Novosib and RNA transposons Ngaro, Rex-Babar, and Jockey (Chalopin et al., 2015; Shao et al., 2018). The presence of Novosib in whale shark extends the presence of this transposon in vertebrates to cartilaginous fishes, as it was previously only detected in teleosts. The whale shark genome appears to contain no full LINE1 repeats, which is consistent with this ancient repeat family being only present among Chondrichthyes as small fragments (Ivancevic et al., 2016), however both whale shark and *Callorhinchus* possess Tx1 L1-like repeats (Chalopin et al., 2015). Overall, the presences and absences of transposable elements in the whale shark is consistent with the patchy distribution of transposable element superfamilies among vertebrates (Chalopin et al., 2015).

### Appendix 1B. Gene completeness assessment using RNA-Seq data

We also assessed mapping of RNA-seq Illumina read data generated from another whale shark individual from blood that were previously generated (Hara et al., 2018) (Supplementary File 2). Sequences were first trimmed using Trimmomatic using the following options: `TruSeq3-PE-2.fa:2:30:10 HEADCROP:13 SLIDINGWINDOW:4:20 MINLEN:50`. We assembled the blood transcriptome reads by mapping to our genome assembly using HISAT v2.0.5 (Kim et al., 2015). The genome was indexed using the command `hisat2-build`. All trimmed reads (paired reads and unpaired reads after trimming) were aligned to the genome using the command hisat2 with the `--dta` flag to output alignments in a format for StringTie, and alignments were sorted using samtools sort. Sorted HISAT alignments were passed to StringTie v1.3.2b under default settings to assemble transcripts (Pertea et al., 2015). We used the `gffcompare` utility packaged with StringTie to match the StringTie reference-based assembly to the RefSeq annotation and quantify the number of exact matches. We assembled 10,873 transcripts (representing 9,606 genes) that exactly matched a RefSeq transcript's internal exon-intron boundaries, a similar number to the 10,990 transcripts (representing 8,938 genes) correctly assembled for human whole B cells in blood using the same software (Pertea et al., 2015). This is despite the transcriptome being fairly incomplete relative to the 2,586

BUSCO v2 conserved vertebrate_odb9 genes (likely because it derives from a single tissue type), with only 1,443 (1,199 single-copy, 244 duplicated) genes recovered as complete, 309 recovered as fragments, and 1,271 orthologs that were missing. This is consistent with core gene analysis (BUSCO, CVG) that suggest the gene completeness of the whale shark is relatively high.

Appendix 1C. Identifying potential missing gene annotations in whale shark and *Callorhinchus*.

Through preliminary orthology determination of chordate proteomes using OrthoFinder v2.2.6 (see Methods), we found numerous gene families conserved in vertebrates were absent in either or both whale shark and *Callorhinchus* RefSeq annotations. Some of these gene families were expected to be in these genomes *a priori* based on the conserved presence of orthologs in other vertebrates, including some that are known to be present in elasmobranchs based on previous research. In this preliminary orthology determination (not presented) 857 gene families were inferred to be lost in the MRCA of Chondrichthyes, 299 gene families were inferred to be gained in the MRCA of Osteichthyes, 1,057 gene families were lost specifically in *Callorhinchus*, and 757 gene families were lost specifically in whale shark. Hence, 1,913 of these gene families were absent from the whale shark genome while 2,213 were absent from the *Callorhinchus* genome, with an overlap of 1,156 gene families that were missing in both. The absence of these gene families in either or both chondrichthyan lineage would affect the inference of the origin and loss of gene families in the MRCA of gnathostomes, the MRCA of Chondrichthyes, and species-specific gains and losses in whale shark and *Callorhinchus*.

To identify putative members of these gene families in whale shark and *Callorhinchus*, we aligned orthologous sequences to both genomes. For whale shark, we aligned *Callorhinchus* proteins for the 757 gene families that would be inferred to be lost in whale shark, and for *Callorhinchus*, we aligned the 1,057 whale shark sequences for gene families that were inferred lost in *Callorhinchus*. We also aligned human, coelacanth, gar, and mouse protein sequences for the 1,156 gene families that were inferred missing in both chondrichthyan genomes to both *Callorhinchus* and whale shark genomes.

We used genBlast v1.39 to identify putative homologous protein sequences (She et al., 2011, 2009), which is a pipeline that utilizes TBLASTN (BLAST+ v2.6.0) to identify putative homologous sequences, then processes these alignments to identify putative orthologues and identify splicing sites. We aligned the orthologous protein sequences to each genome and conservatively selected only the top-ranking putative protein annotated by genBlast for each mapped sequence as a putative ortholog. To exclude proteins annotated by genBlast that were already annotated by RefSeq, we used gffcompare to determine the overlap among annotated regions for genes annotated by genBlast and RefSeq annotations, and we filtered down to only the genBlast annotated proteins that had no overlap with RefSeq annotations (class code of overlap between putative protein and reference sequences coded "u"). Finally, because there may be multiple sequences annotated for each locus (particularly for the bony vertebrate genes where multiple sequences representing the same gene were aligned to each genome), we filtered putative proteins down to the longest sequence per locus, resulting in 764 sequences annotated in whale shark and 376 sequences in *Callorhinchus*. Genes annotated by GenBlast in whale shark and *Callorhinchus* are provided in Supplementary Files 3 and 4, respectively. We then performed orthology assignment again including these additional annotations.

Appendix 2. Comments on Functional Enrichment of Gene Families Specific to Cartilaginous Fishes

In contrast to the enrichment of functions for gene families specific to bony vertebrates (gene families gained in bony vertebrates or ancestral genes lost in cartilaginous fishes), we find much less enrichment of terms for gene families specific to cartilaginous fishes. Firstly, we found no enrichment for function or domain terms in the 276 chondrichthyan-derived genes. While it is possible that novel cartilaginous fish genes are not enriched for any particular function, it is also possible the lack of enrichment may be partially explained by the difference in the number and proportion of genes with annotations: 142 of the 276 (51.4%) gene families that were novel to cartilaginous fishes were annotated, while 301 of the 414 (72.7%) gene families novel to bony vertebrates were annotated, indicating that the novel gene families in cartilaginous fishes were less likely to possess known protein domains or functions. This may have potentially been an effect of the greater divergence of chondrichthyans from well-characterized osteichthyans and the lower level of genomic study that chondrichthyans have received relative to osteichthyans. Therefore, the functional enrichment of gene families that osteichthyan-derived genes may be misled if there is a bias of genes of certain functions to be more difficult to annotate in cartilaginous fishes using InterProScan. This reinforces that some conclusions about the origin and evolution of immune genes in gnathostomes may require more focused study (Dijkstra, 2014; Redmond et al., 2018).

For the 208 gene families lost in osteichthyans and retained in chondricthyans, only one term was enriched: dynein complex. Given the gene families possessing this are not found in the relatively well-studied bony vertebrate genes and were annotated as genes with unknown names, these orthogroups may also be poorly known. Further genomic study and increased taxon sampling in cartilaginous fishes, chordates, and jawless fishes will help to provide further evidence of the distribution of these putative gene families across vertebrates lost in bony vertebrates.

Appendix 3. Additional Results and Discussion of innate immune pathogen receptors in whale shark

NLRs

Human NOD-like receptors (NLRs) are intracellular receptors for a wide array of pathogen-associated molecular patterns (PAMPs) and damage-associated molecular patterns (DAMPs)(Caruso et al., 2014; Fritz and Kufer, 2015; Keestra-Gounder and Tsolis, 2017; Proell et al., 2008; Ting et al., 2008). NLRs play key roles in innate immune defense, mainly through regulation of the inflammatory response through NF-κB and inflammasome (an intracellular multiprotein complex that activates inflammatory responses, and pyroptosis (inflammation-induced programmed cell death) (Caruso et al., 2014; Latz et al., 2013; Proell et al., 2008). NLRs are defined by the shared presence of a NACHT domain (although most also possess C-terminal leucine rich-repeats) and are grouped into three families: the NODs, NLRPs (NALPs), and IPAF. Studies of NLR evolution have revealed large scale-lineage specific expansion of NLRs in some invertebrate lineages (Huang et al., 2008; Rast et al., 2006; Yuen et al., 2014), as well as in teleost fishes (Howe et al., 2016; Laing et al., 2008). The whale shark genomic data were applied here to assess the cartilaginous fish, and ancestral jawed vertebrate, NLR repertoire.

Additional notes on triplicated NOD1: Most strikingly among the NLR results, the whale shark genome harbors three NOD1 genes (UFBOOT=100; Figure 2–Figure supplement 1). All three of the whale shark NOD1 sequences contain detectable NACHT domains and occupy unique genomic locations. This expansion may permit broader bacterial recognition or provide more nuanced responses to different pathogens. Our detailed analyses including elephant shark, for which we found two NOD1s, indicate that the three NOD1s in whale shark all encode detectable NACHT domains and originated in the ancestor of cartilaginous fishes (Figure 2–Figure supplement 1). We named the three whale shark genes NOD1-A, -B, and -C. To further confirm the orthology of whale shark NOD1s we employed synteny analysis which revealed that all three are located on short contigs. NOD1-A and NOD1-B are located on a single contig in the whale shark genome, and are the only genes annotated on this contig (Figure 2–Figure supplement 1). NOD1-C is located on a different two-gene contig, next to a Cystathionine gamma-lyase (CTH)-like gene (Figure 2–Figure supplement 1). Our analyses of human and zebrafish NOD1 regions do not support linkage with Cystathionine gamma-lyase (Figure 2–Figure supplement 2). However, we find that NOD1s in elephant shark are flanked by ZNRF2 and MTURN on one side and this is also the case for human and zebrafish (Figure 2–Figure supplement 2). Interestingly, whale shark ZNRF2 and MTURN are located at the start of a contig (Figure 2–Figure supplement 2), so it is possible that they also link to at least one of the NOD1 contigs. Finally, although NOD1-A and -B are located on different contigs than NOD1-C in whale shark, NOD1-A and NOD1-C orthologs in elephant shark (where NOD1-B appears to be lost) are located next to each other (Figure 2–Figure supplement 2) implying that the three cartilaginous fish NOD1s emerged via tandem duplication events. Human NOD1 plays an important role in intracellular detection of bacterial peptidoglycan amongst a variety of other agonists. As such, we hypothesized that the three copies of NOD1 in whale shark may permit broader ligand recognition or provide more nuanced response to different pathogens/commensals. To this end, we sought to identify key amino acid changes that might impact function. Sequence motifs previously reported as essential to the function of NOD1 are highly conserved in all three whale shark molecules, including the Walker A and B motifs in the NACHT domain necessary for nucleotide binding and hydrolysis, several LxxLL motifs thought to mediate protein-protein interaction, and residues crucial for binding to the RIP2 adaptor protein and downstream signaling (Boyle et al., 2013). Further, the pattern of residue conservation in the LRR domains indicate the whale shark NOD1 molecules, like those of other species, bind ligand on their concave surfaces. However, our attention was drawn to one residue in the LRR domain (E816 in human NOD1) which differed across the whale shark NOD1 molecules. Previous studies have shown this residue contributes to the preferential binding of different peptidoglycan fragments by mouse and human NOD1 (Girardin et al., 2005), supporting our suggestion that the whale shark duplicates have different recognition specificities. Thus, we hypothesize that all three whale shark NOD1 molecules are functional and share an ancestral mechanism of action but have different recognition specificities, potentiating broader and/or more nuanced responses to intracellular pathogens than species with a single NOD1 gene. Finally, examination of our orthogroups found that the three NOD1 orthologs are also conserved in white shark, brownbanded bamboo shark, and clouded catshark (OG0001251, Supplementary File 5).

NRLP-related expansion in the whale shark: As mentioned above, only a single NLRP-like sequence (for which a NACHT domain was not detected) was identified in whale shark, despite the fact that NLRPs are vital for inflammasome activation in studied species (Schroder and Tschopp, 2010). It therefore seems reasonable to suggest that the expanded repertoire of NLRP-related genes in whale shark provides the necessary 'NLRP' inflammasome activators, especially given that we also found a lack of one-to-one orthologs between human NLRP inflammasome activators and zebrafish NLRPs (UFBOOT>=99; Figure 2–Figure supplement 1). Further, lineage-specific clades of NLRPs are indicative of either concerted or rapid birth-death evolution, suggesting that inflammasome evolution (in the form of gene turnover) is highly influenced by environment (i.e. lineage/life-history/environment-specific immune challenges) (Nei and Rooney, 2005; Thomas, 2005). Such processes have also been observed for other immune genes, e.g. antiviral interferons (Redmond et al., 2019) and various adaptive immune genes (Nei et al., 1997).

### RLRs, DICER, and MAVS

MDA5 in whale shark: Although, we generally found one copy of RIG-1 and LGP2, we found two MDA5-like sequences in our analysis. Additional analyses do not provide sufficient support to indicate the presence of an additional MDA5 gene in whale shark. The two protein sequences do not overlap when aligned to other MDA5/RLRs and appear to be at the start and end of different scaffolds, implying that they may in fact be a single gene separated by a genome assembly gap. No sequence from the transcriptome assembly could bridge the gap between these sequences either, but given that no evidence of additional RLRs in vertebrates has been previously reported (even after another round of whole genome duplication in teleosts (Jaillon et al., 2004)) it seems most likely that whale shark possesses a single MDA5 gene.

MAVS in whale shark: Reciprocal BLAST searches between whale shark and the NCBI non-redundant protein database did not reveal a putative MAVS protein in whale shark. Upon more relaxed investigation not requiring reciprocal BLAST hit, one blast hit, from searches of whale shark with MAVS from other vertebrates, revealed a sequence with a CARD domain that appeared similar to those of other MAVS proteins, as well as the CARD domains of MDA5 and RIG-1. Inclusion of this sequence in a phylogenetic analysis of RLR and MAVS CARD domains (using caspase CARDs as outgroups, following Korithoski et al. (2015)) verified that this was in fact whale shark MAVS (UFBOOT=100; Figure 2–Figure supplement 3), and further verified the assignment of whale shark MDA5 and RIG-1 (UFBOOT=100 in all cases; Figure 2–Figure supplement 3). The phylogenetic analysis also placed a coelacanth sequence within the MAVS clade (UFBOOT=100; Figure 2–Figure supplement 3), despite the previous difficulty in identifying such a sequence (Boudinot et al., 2014), suggesting that MAVS is probably ubiquitous in jawed vertebrates. *Callorhinchus* orthologs of all three RLRs, MAVS, and DICER were also identified and/or verified.

TLRs

Toll-like receptors (TLRs) are probably the best known of all innate immune genes, and their functions and evolutionary history have been studied extensively, particularly in comparison to other PRRs (Leulier and Lemaitre, 2008; Roach et al., 2005; Vidya et al., 2018; Wang et al., 2016). TLRs are (typically) membrane spanning receptors that recognize disparate but specific conserved structures, for example TLR3 recognizes viral dsRNA, while TLR4 recognizes bacterial LPS, and TLR9 unmethylated CpG dinucleotides (Akira and Takeda, 2004; Barton and Medzhitov, 2002; Vidya et al., 2018). Vertebrate TLRs consist of a TIR domain involved in signal transduction and leucine-rich repeats (LRRs) that permit target recognition (Akira and Takeda, 2004; Vidya et al., 2018; Wang et al., 2016). Evolutionary studies to date suggest that the vertebrate TLR repertoire is highly conserved, with only small changes between species, whereas large-lineage specific expansions have been observed in invertebrates (Boudinot et al., 2014; Huang et al., 2008; Rast et al., 2006; Roach et al., 2005; Wang et al., 2016). However, a few differences are observed between the teleost and mammal TLR repertoires (partially due to whole genome duplication in teleosts). The spotted gar genome, a close relative of teleosts that did not share the teleost-specific genome duplication, possess a mosaic of mammalian and teleost like TLRs, while an expansion of TLRs in codfishes correlates with loss of CD4 and MHC class II in this lineage (Boudinot et al., 2014; Braasch et al., 2016; Malmstrøm et al., 2016; Roach et al., 2005; Star et al., 2011; Wang et al., 2016). TLRs have been predicted in the *Callorhinchus* genome but await orthology assignment (Venkatesh et al., 2014). Previously, we found a TLR similar to TLR13 and TLR21 in our previous whale shark genome draft assembly (Read et al., 2017); comparison of this with the sequences included in the vertebrate TLR tree below indicates that this previously-studied sequence corresponds to TLR21. The TLR repertoire of whale shark was assessed here to better understand the evolution of vertebrate TLRs and contextualize this with respect to invertebrate TLR expansions and the emergence of adaptive immunity.

<u>TLRs in *Callorhinchus* vs. whale shark:</u> In the TLR tree presented here, *Callorhinchus* does not possess orthologs of TLR21 or TLR29, but does possess orthologs of TLR14/18 and TLR25 (Boudinot et al., 2014; Wcisel et al., 2017). Thus, while the TLR repertoires of whale shark and elephant shark are quite well conserved, repertoire differences do exist; understanding the impact of which requires functional characterization of TLR14/18, TLR25, and TLR29. Extrapolating from our data, the most recent common ancestor of whale shark and elephant shark must have possessed at least 12 TLRs from different ancestral jawed vertebrate lineages. This implies that cartilaginous fishes possess a highly similar number of TLRs to mammals (approx. 10). However the whale shark TLR repertoire is novel in that it contains a mix of orthologs to the classical mammalian TLRs and 'fish-specific' TLRs (similar to spotted gar (Braasch et al., 2016; Wcisel et al., 2017)), as well as to TLR27 and the new TLR29. These last two appear to be absent from both mammals and teleosts, but present in the so-called 'living fossil' lineages (e.g. sharks, coelacanths, gars).

TLR9: Intriguingly, despite TLR9 evolution being well studied, in our analysis we observe two maximally supported TLR9 sister clades (Figure 2–Figure supplement 4), each of which contains an array of mutually exclusive vertebrate taxa except both contain spotted gar orthologs (Wcisel et al., 2017). We also found evidence that a third TLR9 group falls sister to these (UFBOOT=94; Figure 2–Figure supplement 4). For the newly discovered jawed vertebrate TLR9 lineages we suggest the following nomenclature: TLR9a (includes the originally characterized mammalian TLR9), TLR9b (or TLR30; includes the new whale shark TLR9), TLR9c (or TLR31) (Figure 2–Figure supplement 4). A similar scenario appears to have played out in TLR13 evolution, where three potential lineages are also detectable TLR13a (including mammalian TLR13), TLR13b (TLR32; including reptilian TLR13) and TLR13c (TLR33) (Figure 2–Figure supplement 4), although only a single lineage appears to have existed in the jawed vertebrate ancestor.

Ancestral reconstruction of TLR repertoires: Based on our rooted TLR tree, it is possible to estimate a minimal ancestral jawed vertebrate TLR repertoire, by inferring gene loss based on the species possessing a sister gene(Peterson and Sperling, 2007; Redmond et al., 2017). As an example, in the TLR tree presented here, TLR7 and TLR8 are sister clades, with both containing representatives of each of the major jawed vertebrate groups (Figure 2–Figure supplement 4). As such although cartilaginous fish orthologs of TLR7 and TLR8 exist (Figure 2–Figure supplement 4), if the tree contained no TLR8 orthologs, their existence could be inferred, because the presence of a cartilaginous fish TLR7 ortholog means that these genes split prior to speciation between cartilaginous fishes and bony vertebrates. Using this phylogenetic logic, it is apparent that at least 17 TLRs existed in the last common ancestor of jawed vertebrates, TLR1/6/10, TLR2/28, TLR3, TLR4, TLR5/5SL (possibly both), TLR7, TLR8, TLR9 (possibly three of these), TLR11/12/16/19/20/26, TLR13, TLR14/18, TLR15, TLR21, TLR22/23 (possibly both), TLR25, TLR27, TLR29 (inferred from Figure 2–Figure supplement 4). This number is much higher than what is commonly found in most jawed vertebrate lineages (excluding gar, teleosts, and particularly Atlantic cod, for the reasons stated above (Boudinot et al., 2014; Braasch et al., 2016; Roach et al., 2005; Star et al., 2011; Wcisel et al., 2017)). This result thus supports a scenario of convergently decreasing TLR repertoire complexity in the major jawed vertebrate lineages since their last common ancestor.

There are some caveats to this count, e.g. root placement is taken to be reasonably accurate, invertebrates were not included, and common vertebrate gene tree errors such as teleosts grouping sister to all other jawed vertebrates, or slowly evolving species grouping together were permitted. Additional potential errors may also exist, such as the placement of TLR15, which here is taken to be lost or not yet found in all non-reptile (incl. avian) lineages, but could conceivably be a highly divergent ortholog of another TLR group that is not easily phylogenetically placed due to extreme rate asymmetry (Holland et al., 2017; Manousaki et al., 2011; Redmond et al., 2018).

In addition to considering the ancestral jawed vertebrate TLR repertoire, the inclusion of lamprey TLRs in the tree meant that an estimate of the minimal ancestral vertebrate TLR repertoire could also be considered (Figure 2–Figure supplement 4). Using the same approach as above, and considering the same caveats, the last common ancestor of all vertebrates likely had at least 15 TLRs, spanning the full array of TLR superfamilies, and presumably their functions; TLR1/6/10, TLR2/28, TLR3, TLR4, TLR5/5SL, TLR7/8, TLR9,

TLR11/12/16/19/20/22/23/26, TLR13, TLR14/18, TLR15, TLR21/29, TLR24, TLR25, TLR27 (Figure 2–Figure supplement 4). This result implies an increase in TLR repertoire between the last common ancestors of all vertebrates and jawed vertebrates. Lineage-specific duplications in jawed vertebrates, e.g. TLR7 and TLR8, TLR21, and TLR29 (each of which are co-orthologs of lamprey lineages for which we suggest the names TLR7/8-like [or TLR34], and TLR21/29-like [or TLR35]; Figure 2–Figure supplement 4), have played an important role in this. However, the uniqueness of the lamprey genome (Smith et al., 2013), as well as the relatively poor sampling of jawless vertebrates could also contribute to this small discrepancy.

#### Adaptive immunity and the evolution of PRR repertoires

Early hypotheses to explain the stark differences between the relatively conserved PRR repertoires in vertebrates, and the remarkable PRR expansions in invertebrates (particularly in deuterostomes) suggested that the adaptive immune system negated the need for diversification of novel PRRs in vertebrates. Here, although we found evidence for a core set of PRRs that appear to have been 'locked in' during early vertebrate evolution alongside the appearance of the new adaptive immune system, we found that NLR expansions in jawed vertebrates are common and extensive, while jawed vertebrate RLRs (although major expansion of this family in invertebrates also seems not to have occurred) and TLRs appear to fit a model where expansions are constrained. These differences may be due to the complexity of interactions between adaptive and innate immune systems, as well as those between host and pathogen.

For example, the RLRs are the most conserved set of examined PRRs in our analyses, being nearly identical in repertoire in all jawed vertebrates. Experimental MDA5 duplication improves the immune response, but also accelerates autoimmunity (Crampton et al., 2012). MDA5 is capable of functionally replacing RIG-1 following loss, implying a certain level of redundancy (Xu et al., 2016). Together, this suggests that the RLR repertoire in vertebrates is constrained to maintain balance between a maximally robust response while avoiding autoimmunity. However, it is important to note that large expansion of this family in invertebrates has not been reported, and so it is possible that adaptive immunity has had an inconsequential role in orchestrating RLR repertoire evolution.

TLR evolution appears to occur more readily through gene loss than expansion in vertebrates, perhaps fitting a scenario where TLRs are lost if they become obsolete or are subverted by a pathogen. In general, TLRs fit well with the idea of adaptive immunity replacing or constraining the need for innate innovation through extensive gene duplication, however new TLRs have emerged independently of genome duplication during vertebrate evolution (e.g. TLR1/6/10 duplications, TLR11/12 duplication), suggesting that despite the presence of adaptive immunity there is still a place for new germline encoded TLRs.

NLR expansions have occurred multiple times during jawed vertebrate evolution and to independent NLR family members, firmly rejecting a situation where adaptive immunity relieves the need for PRR expansions. These expansions (assuming rapid turnover) may be required to maintain functional relevance for rapid detection of (new) lineage-specific pathogens or inflammasome activation (Latz et al., 2013; Schroder and Tschopp, 2010).

### Appendix 4. Body size evolution in Chondrichthyes.

Gigantism in vertebrates including elephants and whales are associated with a shift to a new rate regime of body size evolution (Puttick and Thomas, 2015; Slater et al., 2017). We tested if whale sharks are also unexpectedly large given background rates of body size evolution in cartilaginous fishes, which would imply that other aspects of its biological evolution may have also shifted in association with this gigantism.

Patterns of body size evolution were estimated using a published distribution of 500 time trees and log-transformed body mass data for chondrichthyans with 10 fossil calibrations(Stein et al., 2018). Analysis across a posterior distribution of trees allows for accounting for uncertainty in phylogenetic inference in estimating patterns of body size evolution. Body mass data for chondrichthyans from a previous study (Stein et al., 2018) were kindly provided by Chris Mull. Mass data were log-transformed for comparative analyses. Time-varying rates of body size evolution were estimated using BAMM 2.5.0 and BAMMtools 2.1.6 (Rabosky, 2014; Rabosky et al., 2014). For each of the trees, we used setBAMMpriors to calculate appropriate prior probabilities that were scaled for each tree. Each MCMC run was run with 4 chains for at least 30M generations, sampling once every 10K generations, and discarding the first 10% of samples as burn-in. We assessed convergence using the effectiveSize() function in coda v0.19-1 and determined if any parameters were below 200 effective samples. Runs with insufficient samples were re-run at 50M generations. All runs reached convergence (i.e. had over 200 effective samples) with 50M generations. After discarding burn-in, 2000 samples were extracted from the event data for each tree. We determined the marginal odds ratio of shifts for all branches, including the branch leading to the whale shark, to assess the support for a rate shift in body size evolution leading to the whale shark. To compute mean rates of body size evolution across branches for each posterior sample, we estimated the rate of evolution including the single branch representing the whale shark using getCladeRates(). To compute the mean rates of body size evolution in the background rate of body size evolution, for each posterior sample, we computed the rates of body size evolution across all branches that shared the background rate regime (the rate regime that originated at the root).

Numerous independent shifts in body size evolution were inferred in the chondrichthyans. including along the branch leading to the whale shark. We recovered a significant shift in the rate of body size evolution along the branch leading to the whale shark with a mean marginal odds ratio of 241.65, demonstrating strong support for a shift in the rate of body size evolution in the branch leading to the whale shark and demonstrating that the gigantism in the whale shark is not inferred to be the result of neutral, background rates of body size evolution. The mean rate of body size evolution along the whale shark branch across all posterior samples and all 500 tree samples (1M total samples) was 0.532 log-grams per million years, relative to 0.117 log-grams per million years estimated for the background rate in chondrichthyans (Supplementary figure 3). The difference in mean rate of body size evolution between the whale shark and the background across all samples is 4.31 times the background rate. These results support that the gigantic body size of the whale shark is due to a shift to a novel rate regime of body size evolution.

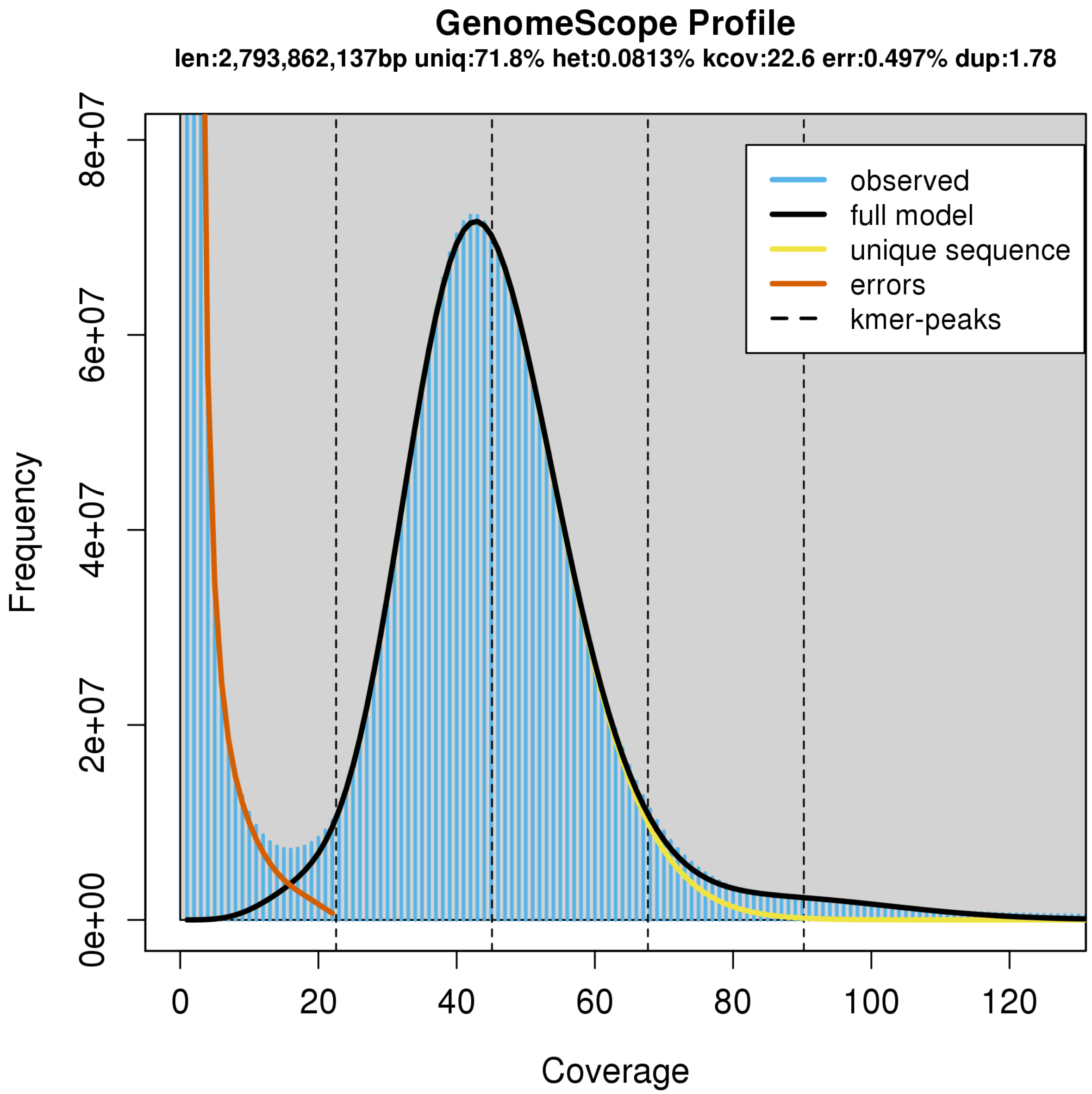


Supplementary figure 1. Characteristics of the whale shark genome assembly by *k*-mer profiling of raw Illumina reads by GenomeScope. GenomeScope fits a model to estimate genome parameters including heterozygosity (het), an estimated genome size (len), the unique proportion of the genome (uniq; as opposed to the remainder which would be repetitive genome length). Profiling of *k*-mers reveals high coverage sequencing as well as low heterozygosity. Consistent with low heterozygosity, most of the *k*-mers form one peak centered around roughly 40x coverage, and do not form another peak centered at roughly half the coverage that would represent *k*-mers arising from heterozygous alleles.

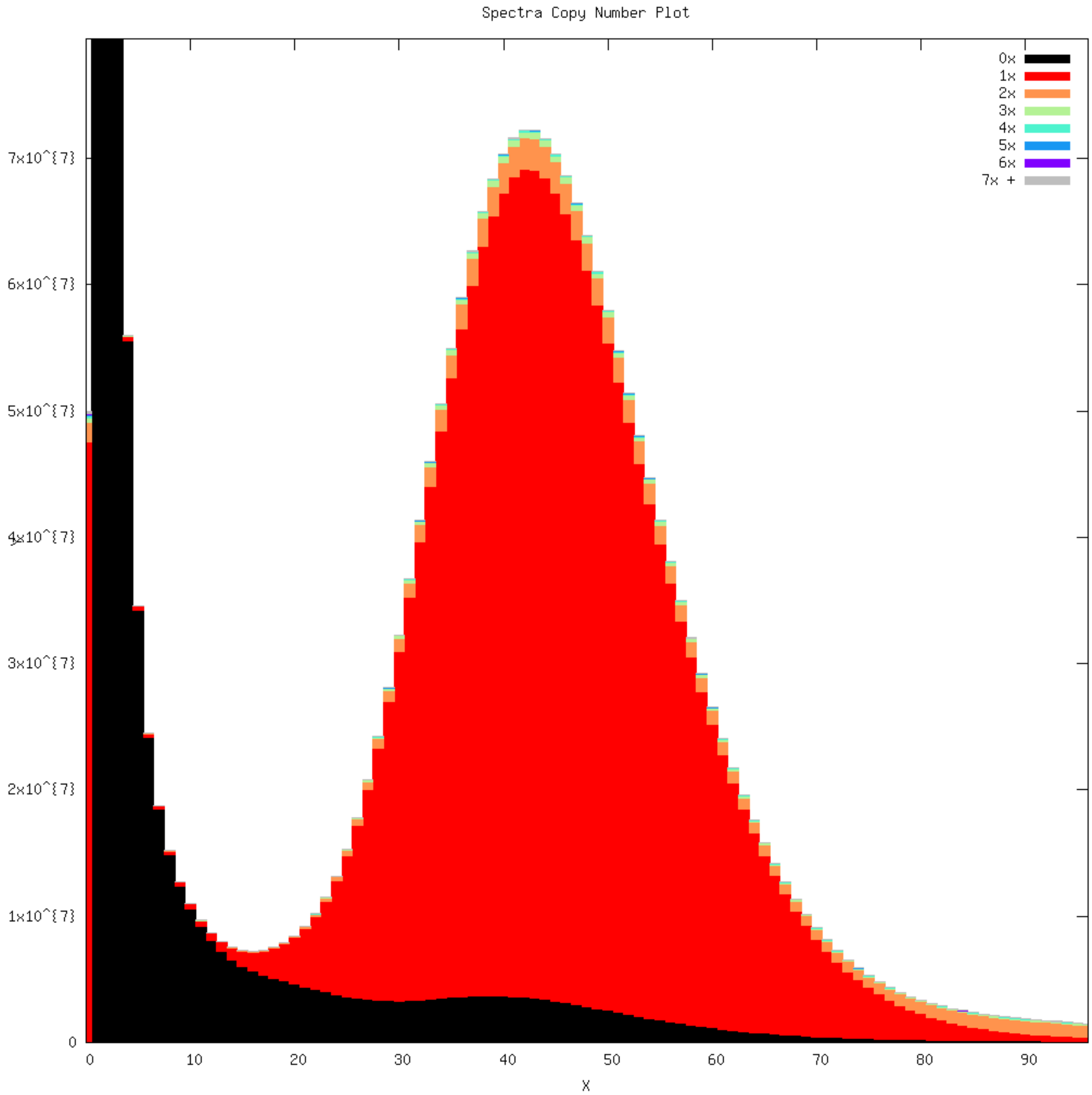


Supplementary figure 2. *k*-mer profile overlaid with copy number representation within the genome assembly as produced by KAT. *k*-mers arising from error in Illumina raw reads on the left part of the plot are not within the assembly (represented 0x). Most of the *k*-mers in the genome assembly are represented by a single copy (1x, red), suggesting an accurate haploid genome assembly with few diploid alleles assembled as separate contigs.

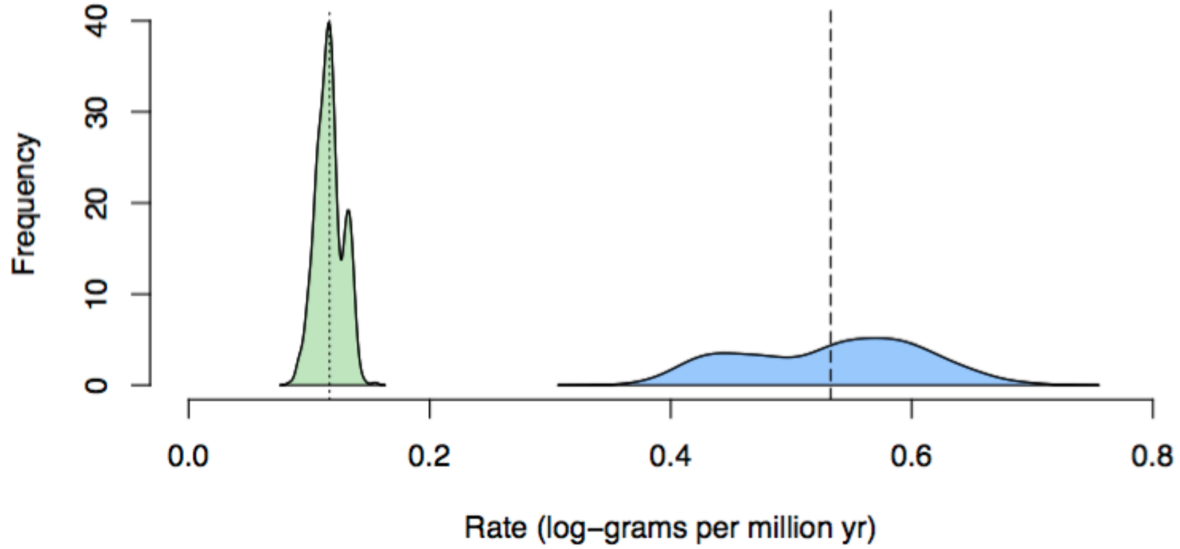


Supplementary figure 3. Distribution of mean estimated rate of body size evolution of the posterior distribution estimated for each tree sample (n = 500) for the background in Chondrichthyes (green) and for the whale shark (blue). Dotted line indicates mean estimated rate for Chondrichthyes across all tree samples, while the dashed line indicates mean estimated rate for the whale shark.

Supplementary table 1. Comparison of whale shark genome assemblies.

| | Read et al. 2017 | Hara et al. 2018 | Weber et al. 2020 (all sequences) | Current assembly |
|---|---|---|---|---|
| DNA source | male liver and spleen | male liver and spleen<br>(from Read et al. 2017) | male heart | male liver and spleen<br>(same as Read et al. 2017) |
| Illumina raw read data (Gbp) | 275.6 | 275.6<br>(from Read et al. 2017) | 526.0 | 275.6<br>(from Read et al. 2017) |
| PacBio raw read data (Gbp) | – | – | – | 61.8 |
| Total assembly size (Gbp) | 2.57 | 2.66 | 3.2 | 2.96 |
| Number of scaffolds | 997,976 | 155,537 | 3,305,708 | 39,176 |
| Scaffold N50 (Kbp) | 5.425 | 326 | 2,564 | 149.8 |
| Number of contigs | 1,213,000 | 15,923,929 | 3,497,228 | 57,333 |
| Contig N50 (bp) | 5,304 | 3,579 | 35,692 | 144,422 |

Supplementary table 2. Repetitive element content of the whale shark genome assembly (for methods see Appendix 1A).

| | Number of elements | Length occupied (bp) | Percentage of sequence |
|---|---|---|---|
| SINEs: | 238546 | 45540346 | 1.55 |
| ALUs | 0 | 0 | 0.00 |
| MIRs | 0 | 0 | 0.00 |
| LINEs: | 2198969 | 974884225 | 33.25 |
| LINE1 | 0 | 0 | 0.00 |
| LINE2 | 64356 | 18837437 | 0.64 |
| L3/CR1 | 1421231 | 638508824 | 21.78 |
| LTR elements: | 346587 | 116780647 | 3.98 |
| ERVL | 1342 | 1253585 | 0.04 |
| ERVL-MaLRs | 0 | 0 | 0.00 |
| ERV_classI | 4050 | 2057150 | 0.07 |
| Unclassified: | 1463974 | 318205896 | 10.85 |
| Total interspersed repeats: | | 1475831239 | 50.34% |
| small RNA: | 3192 | 7876017 | 0.27 |
| Satellites: | 6333 | 2523637 | 0.09 |
| Simple repeats: | 177884 | 14474709 | 0.49 |
| Low complexity: | 36180 | 3453271 | 0.12 |

Supplementary table 3. BUSCO v2 and CVG results. BUSCO v2 and CVG results for brownbanded bamboo shark and cloudy catshark were those reported by Hara et al. (2018) Supplementary figure 1d, who did not report Complete Single-copy and Complete Duplicate numbers and only reported percentages. *Callorhinchus* CVG scores are reported on the gVolante database (https://gvolante.riken.jp/script/database.cgi, accessed 2021 Jan 19). The BUSCO v2 set has 2,586 vertebrate orthologs, while the CVG has 233 total genes. Note CVG does not report if complete are single copy or duplicated. Percentages in parentheses. Note also that *Callorhinchus* was used in the ortholog design of both sets and therefore BUSCO and CVG overestimate its completeness.

| **BUSCO (2,586 total)** | **Complete** | **Complete Single copy** | **Complete Duplicate** | **Fragments/ Partial** | **Missing** |
|---|---|---|---|---|---|
| whale shark | 2033 (78.7) | 1967 (76.1) | 66 (2.6) | 323 (12.5) | 230 (8.8) |
| *Callorhinchus* | 2459 (95.1) | 1603 (62.0) | 856 (33.1) | 68 (2.6) | 59 (2.3) |
| bamboo shark | (92.6) | | | (5.5) | (1.9) |
| cloudy catshark | (81.8) | | | (14.1) | (4.1) |
| **CVG (233 total)** | **Complete** | | | **Fragments/ Partial** | **Missing** |
| whale shark | 199 (85.4) | | | 28 (12.0) | 6 (2.6) |
| *Callorhinchus* | 227 (97.4) | | | 4 (1.7) | 2 (0.86) |
| bamboo shark | (93.9) | | | (3.1) | (3.0) |
| cloudy catshark | (88.2) | | | (8.9) | (2.9) |

Supplementary table 4. Chordate species with whole genomic data included in comparative genomic analyses.

| Species | Common Name | Gene Model Source | Assembly Version |
|---|---|---|---|
| *Branchiostoma floridae* | Amphioxus | RefSeq | GCF_000003815.1_Version_2 |
| *Ciona intestinalis* | Sea squirt | Ensembl 89 | GCA_000224145.1 |
| *Petromyzon marinus* | Sea lamprey | Ensembl 89 | Pmarinus_7.0 |
| *Callorhinchus milii* | Elephant Shark, Ghost Shark* | RefSeq | GCF_000165045.1 |
| *Chiloscyllium punctatum* | Brownbanded Bamboo shark | https://transcriptome.riken.jp/squalomix/ | Cpunctatum_v1.0 |
| *Scyliorhinus torazame* | Cloudy catshark | https://transcriptome.riken.jp/squalomix/ | Storazame_v1.0 |
| *Carcharodon carcharias* | Great white shark | https://datadryad.org/stash/dataset/doi:10.5061/dryad.9r2p3ks | ASM360424v1 |
| *Lepisosteus oculatus* | Spotted gar | Ensembl 89 | LepOcu1 |
| *Scleropages formosus* | Asian bonytongue | Ensembl 99 | ASM162426v1 |
| *Danio rerio* | Zebrafish | Ensembl 89 | GRCz11 |
| *Esox lucius* | Northern pike | Ensembl 99 | Eluc_V3 |
| *Oreochromis niloticus* | Nile tilapia | Ensembl 89 | Orenil1.0 |
| *Gadus morhua* | Atlantic cod | Ensembl 89 | gadMor1 |
| *Gasterosteus aculeatus* | Three-spined stickleback | Ensembl 89 | BROAD S1 |
| *Dichotomyctere nigroviridis* | Green Spotted Pufferfish | Ensembl 89 | TETRAODON 8.0 |
| *Takifugu rubripes* | Fugu | Ensembl 89 | FUGU5 |
| *Mola mola* | Ocean sunfish | GigaDB: http://dx.doi.org/10.5524/100214 | ASM169857v1 |
| *Latimeria chalumnae* | Coelacanth | Ensembl 89 | LatCha1 |
| *Xenopus tropicalis* | Western clawed frog | Ensembl 89 | JGI 4.2 |

| | | | |
|---|---|---|---|
| *Anolis carolinensis* | Green anole | Ensembl 89 | AnoCar2.0 |
| *Alligator mississippiensis* | American alligator | RefSeq | GCF_000281125.3 |
| *Gallus gallus* | Chicken | Ensembl 89 | Gallus_gallus-5.0 |
| *Ornithorhynchus anatinus* | Platypus | Ensembl 89 | OANA5 |
| *Monodelphus domestica* | Gray short-tailed opposum | Ensembl 89 | monDom5 |
| *Dasypus novemcinctus* | Nine-banded armadillo | Ensembl 89 | Dasnov3.0 |
| *Loxodonta africana* | African elephant | Ensembl 89 | loxAfr3 |
| *Procavia capensis* | Rock hyrax | Ensembl 89 | proCap1 |
| *Homo sapiens* | Human | Ensembl 89 | GRCh38 |
| *Mus musculus* | Mouse | Ensembl 89 | GRCm38 |
| *Canis lupus familiaris* | Dog | Ensembl 89 | CanFam3.1 |
| *Sus scrofa* | Pig | Ensembl 89 | Sscrofa10.2 |
| *Bos taurus* | Cow | Ensembl 89 | UMD3.1 |
| *Tursiops truncatus* | Bottlenose dolphin | Ensembl 89 | turTru1 |
| *Balaenoptera acutorostrata scammoni* | Minke whale | RefSeq | GCF_000493695.1 |
| *Balaena mysticetus* | Bowhead whale | http://www.bowhead-whale.org/ | 1.0 |

Supplementary table 5. Summary of functional enrichment tests of gene families gained and lost throughout chordate evolution. We tested whether or not gene families in the foregrounds were enriched for functional terms and domains (GO, Pfam) relative to the background of what was present at a relevant ancestor.

| Foreground (n) | Background (n) | Num of significantly enriched terms | Tablesfor List of Enriched Functions/Domains |
|---|---|---|---|
| Novel genes for MRCA of Olfactores (989) | Present in MRCA of Olfactores (10255) | 13 | Supp Table 6 |
| Novel genes for MRCA of vertebrates (714) | Present in MRCA of vertebrates (10738) | 14 | Supp Table 7 |
| Novel genes for MRCA of gnathostomes (2106) | Present in MRCA of gnathostomes (12815) | 33 | Supp Table 8 |
| Novel genes for MRCA of Chondrichthyes (124) | Present in MRCA of gnathostomes (12815) | 0 | – |
| Novel genes for MRCA of Osteichthyes (298) | Present in MRCA of gnathostomes (12815) | 8 | Supp Table 9 |
| Gene lost in MRCA of Chondrichthyes (729) | Present in MRCA of gnathostomes (12815) | 22 | Supp Table 10 |
| Lost in MRCA of Osteichthyes (145) | Present in MRCA of gnathostomes (12815) | 0 | – |

Supplementary table 12. Fossil calibration age ranges, and the result of fossil concordance analysis. Discordant fossils were excluded from divergence time analysis. All age ranges are derived from Benton et al. (2014), except for the age of Chondricthyes, which were derived from Coatest et al. (2017).

Vertebrata = MRCA of *Petromyzon* + Gnathostomata (= tree height, 457.5–636.1 Ma), Gnathostomata = MRCA of Chondrichthyes + Osteichthyes (420.7–468.4 Ma), Osteichthyes = MRCA of Actinopterygii + Sarcopterygii (420.7–444.9 Ma)*, Sarcoptyergii = MRCA of *Latimeria* + Tetrapoda (408–427.9 Ma), Tetrapoda = MRCA of Lissamphibia + Amniota (337–351 Ma), Amniota = MRCA of Eureptilia + Diapsida (318–332.9 Ma)*, Mammalia = MRCA of Monotremata + Theria (164.9–201.5)*, Theria = MRCA of Marsupialia + Placentalia (156.3–169.6 Ma)*, Placentalia = MRCA of Atlantogenata + Boreutheria (61.6–164.6 Ma), and Artiodactyla = MRCA of Cetaceans + other artiodactyls (52.4–66 Ma).

| Crown node name | Min Age (Ma) | Max Age (Ma) | Concordant? |
|---|---|---|---|
| Vertebrata (root node) | 457.5 | 636.1 | Y |
| Gnathostomata | 420.7 | 468.4 | Y |
| Chondrichthyes | 358 | 422.4 | Y |
| Osteichthyes | 420.7 | 444.9 | Y |
| Sarcopterygii | 408 | 427.9 | Y |
| Tetrapoda | 337 | 351 | Y |
| Amniota | 318 | 332.9 | Y |
| Mammalia | 164.9 | 201.5 | N |
| Theria | 156.3 | 169.6 | N |
| Placentalia | 61.6 | 164.6 | N |
| Artiodactyla | 52.4 | 66 | N |

Supplementary table 13. Whale shark PRR gene accessions. Sequences that have identical or are isoforms of the same gene are indicated. TLR9 and TLR29 sequences that were not annotated are also indicated.

| | |
|---|---|
| TLRs | |
| TLR1/6/10 | XP_020390742.1<br>XP_020390749.1 (identical) |
| TLR2/28 | XP_020367827.1<br>XP_020367828.1 (identical)<br>XP_020379248.1 |
| TLR3 | XP_020375139.1<br>XP_020375140.1 (identical) |
| TLR7 | XP_020376237.1 |
| TLR8 | XP_020376247.1 |
| TLR9 | XP_020391373.1<br>Another on: tig000316855 |
| TLR21 | XP_020378687.1 |
| TLR22/23 | XP_020374252.1 |
| TLR27 | XP_020384134.1 |
| TLR29 | XP_020388546.1<br>Another on: tig000303350 |
| RLRs, MAVS, and DICER | |
| RIG-1 | XP_020382456.1 |
| MDA5 | XP_020377384.1<br>XP_020375939.1 (see text) |
| LGP2 | XP_020377833.1<br>XP_020377839.1 (identical) |
| MAVS | XP_020379351.1 |
| DICER | XP_020385790.1 |
| NLRs (*=detected NACHT domain) | |
| NOD1 | XP_020377166.1*<br>XP_020377167.1*<br>XP_020372421.1* |
| NOD2 | XP_020371316.1*<br>XP_020371320.1 (same gene) |

| | |
|---|---|
| | XP_020371321.1 (same gene) |
| NLRC3 (NOD3) | XP_020391377.1* |
| NLRC5 (NOD4) | XP_020383359.1*<br>XP_020383358.1<br>See also:<br>XP_020370077.1 |
| NLRX1 (NOD5) -like | XP_020372884.1 |
| CIITA | XP_020392242.1*<br>XP_020392245.1 (same gene) |
| NLRC4 (IPAF) -like | XP_020392928.1 |
| NWD1-like | XP_020378861.1<br>See also:<br>XP_020381701.1<br>XP_020391917.1 |
| NAIP (BIRC1) -like | XP_020373169.1<br>XP_020373170.1 |
| "Fish-specific" FISNA domain containing NLR | XP_020386246.1*<br>XP_020374464.1<br>XP_020374622.1 |
| NLRP | XP_020376766.1 |
| Whale Shark lineage-specific expansion | XP_020379593.1*<br>XP_020380299.1*<br>XP_020370463.1*<br>XP_020383553.1*<br>XP_020383554.1*<br>XP_020385964.1*<br>XP_020392482.1*<br>XP_020387670.1*<br>STRGRTYPT.52674*<br>XP_020380409.1*<br>XP_020387523.1*<br>XP_020373146.1*<br>XP_020375477.1*<br>XP_020391354.1*<br>XP_020389366.1<br>XP_020387859.1<br>XP_020387858.1 |
| TEP-1-like | XP_020381121.1 |
| NB-ARC domain containing proteins<br>(APAF-1) | XP_020387052.1<br>XP_020390367.1 |

| | |
|---|---|
| Other possible NLRs (unclear orthology/ novel lineages) | XP_020391906.1*<br>XP_020391908.1 (same gene)<br>XP_020391907.1<br>XP_020367001.1*<br>XP_020386151.1<br>XP_020391062.1<br>XP_020378394.1<br>XP_020380031.1 |

References:


Akira S, Takeda K. 2004. Toll-like receptor signalling. *Nat Rev Immunol* **4**:499–511.

Barton GM, Medzhitov R. 2002. Toll-like receptors and their ligands. *Curr Top Microbiol Immunol* **270**:81–92.

Boudinot P, Zou J, Ota T, Buonocore F, Scapigliati G, Canapa A, Cannon J, Litman G, Hansen JD. 2014. A tetrapod-like repertoire of innate immune receptors and effectors for coelacanths. *J Exp Zool B Mol Dev Evol* **322**:415–437.

Boyle JP, Mayle S, Parkhouse R, Monie TP. 2013. Comparative Genomic and Sequence Analysis Provides Insight into the Molecular Functionality of NOD1 and NOD2. *Front Immunol* **4**:317.

Braasch I, Gehrke AR, Smith JJ, Kawasaki K, Manousaki T, Pasquier J, Amores A, Desvignes T, Batzel P, Catchen J, Berlin AM, Campbell MS, Barrell D, Martin KJ, Mulley JF, Ravi V, Lee AP, Nakamura T, Chalopin D, Fan S, Wcisel D, Cañestro C, Sydes J, Beaudry FEG, Sun Y, Hertel J, Beam MJ, Fasold M, Ishiyama M, Johnson J, Kehr S, Lara M, Letaw JH, Litman GW, Litman RT, Mikami M, Ota T, Saha NR, Williams L, Stadler PF, Wang H, Taylor JS, Fontenot Q, Ferrara A, Searle SMJ, Aken B, Yandell M, Schneider I, Yoder JA, Volff J-N, Meyer A, Amemiya CT, Venkatesh B, Holland PWH, Guiguen Y, Bobe J, Shubin NH, Di Palma F, Alföldi J, Lindblad-Toh K, Postlethwait JH. 2016. The spotted gar genome illuminates vertebrate evolution and facilitates human-teleost comparisons. *Nat Genet* **48**:427–437.

Campbell MS, Law M, Holt C, Stein JC, Moghe GD, Hufnagel DE, Lei J, Achawanantakun R, Jiao D, Lawrence CJ, Ware D, Shiu S-H, Childs KL, Sun Y, Jiang N, Yandell M. 2014. MAKER-P: a tool kit for the rapid creation, management, and quality control of plant genome annotations. *Plant Physiol* **164**:513–524.

Caruso R, Warner N, Inohara N, Núñez G. 2014. NOD1 and NOD2: signaling, host defense, and inflammatory disease. *Immunity* **41**:898–908.

Chalopin D, Naville M, Plard F, Galiana D, Volff J-N. 2015. Comparative analysis of transposable elements highlights mobilome diversity and evolution in vertebrates. *Genome Biol Evol* **7**:567–580.

Crampton SP, Deane JA, Feigenbaum L, Bolland S. 2012. Ifih1 gene dose effect reveals MDA5-mediated chronic type I IFN gene signature, viral resistance, and accelerated autoimmunity. *J Immunol* **188**:1451–1459.

Dijkstra JM. 2014. TH2 and Treg candidate genes in elephant shark. *Nature* **511**:E7–9.

Ellinghaus D, Kurtz S, Willhoeft U. 2008. LTRharvest, an efficient and flexible software for de novo detection of LTR retrotransposons. *BMC Bioinformatics* **9**:18.

Fritz JH, Kufer TA. 2015. Editorial: NLR-Protein Functions in Immunity. *Front Immunol* **6**:306.

Girardin SE, Jéhanno M, Mengin-Lecreulx D, Sansonetti PJ, Alzari PM, Philpott DJ. 2005. Identification of the critical residues involved in peptidoglycan detection by Nod1. *J Biol Chem* **280**:38648–38656.

Gregory TR. 2005. Synergy between sequence and size in large-scale genomics. *Nat Rev Genet* **6**:699–708.

Han Y, Wessler SR. 2010. MITE-Hunter: a program for discovering miniature inverted-repeat transposable elements from genomic sequences. *Nucleic Acids Res* **38**:e199.

Hara Y, Yamaguchi K, Onimaru K, Kadota M, Koyanagi M, Keeley SD, Tatsumi K, Tanaka K, Motone F, Kageyama Y, Nozu R, Adachi N, Nishimura O, Nakagawa R, Tanegashima C, Kiyatake I, Matsumoto R, Murakumo K, Nishida K, Terakita A, Kuratani S, Sato K, Hyodo S, Kuraku S. 2018. Shark genomes provide insights into elasmobranch evolution and the origin of vertebrates. *Nat Ecol Evol*. doi:10.1038/s41559-018-0673-5

Holland PWH, Marlétaz F, Maeso I, Dunwell TL, Paps J. 2017. New genes from old: asymmetric divergence of gene duplicates and the evolution of development. *Philos Trans R Soc Lond*

*B Biol Sci* **372**. doi:10.1098/rstb.2015.0480
Howe K, Clark MD, Torroja CF, Torrance J, Berthelot C, Muffato M, Collins JE, Humphray S, McLaren K, Matthews L, McLaren S, Sealy I, Caccamo M, Churcher C, Scott C, Barrett JC, Koch R, Rauch G-J, White S, Chow W, Kilian B, Quintais LT, Guerra-Assunção JA, Zhou Y, Gu Y, Yen J, Vogel J-H, Eyre T, Redmond S, Banerjee R, Chi J, Fu B, Langley E, Maguire SF, Laird GK, Lloyd D, Kenyon E, Donaldson S, Sehra H, Almeida-King J, Loveland J, Trevanion S, Jones M, Quail M, Willey D, Hunt A, Burton J, Sims S, McLay K, Plumb B, Davis J, Clee C, Oliver K, Clark R, Riddle C, Eliott D, Threadgold G, Harden G, Ware D, Mortimer B, Kerry G, Heath P, Phillimore B, Tracey A, Corby N, Dunn M, Johnson C, Wood J, Clark S, Pelan S, Griffiths G, Smith M, Glithero R, Howden P, Barker N, Stevens C, Harley J, Holt K, Panagiotidis G, Lovell J, Beasley H, Henderson C, Gordon D, Auger K, Wright D, Collins J, Raisen C, Dyer L, Leung K, Robertson L, Ambridge K, Leongamornlert D, McGuire S, Gilderthorp R, Griffiths C, Manthravadi D, Nichol S, Barker G, Whitehead S, Kay M, Brown J, Murnane C, Gray E, Humphries M, Sycamore N, Barker D, Saunders D, Wallis J, Babbage A, Hammond S, Mashreghi-Mohammadi M, Barr L, Martin S, Wray P, Ellington A, Matthews N, Ellwood M, Woodmansey R, Clark G, Cooper J, Tromans A, Grafham D, Skuce C, Pandian R, Andrews R, Harrison E, Kimberley A, Garnett J, Fosker N, Hall R, Garner P, Kelly D, Bird C, Palmer S, Gehring I, Berger A, Dooley CM, Ersan-Ürün Z, Eser C, Geiger H, Geisler M, Karotki L, Kirn A, Konantz J, Konantz M, Oberländer M, Rudolph-Geiger S, Teucke M, Osoegawa K, Zhu B, Rapp A, Widaa S, Langford C, Yang F, Carter NP, Harrow J, Ning Z, Herrero J, Searle SMJ, Enright A, Geisler R, Plasterk RHA, Lee C, Westerfield M, de Jong PJ, Zon LI, Postlethwait JH, Nüsslein-Volhard C, Hubbard TJP, Roest Crollius H, Rogers J, Stemple DL. 2013. The zebrafish reference genome sequence and its relationship to the human genome. *Nature* **496**:498–503.
Howe K, Schiffer PH, Zielinski J, Wiehe T, Laird GK, Marioni JC, Soylemez O, Kondrashov F, Leptin M. 2016. Structure and evolutionary history of a large family of NLR proteins in the zebrafish. *Open Biol* **6**:160009.
Huang S, Yuan S, Guo L, Yu Y, Li J, Wu T, Liu T, Yang M, Wu K, Liu H, Ge J, Yu Y, Huang H, Dong M, Yu C, Chen S, Xu A. 2008. Genomic analysis of the immune gene repertoire of amphioxus reveals extraordinary innate complexity and diversity. *Genome Res* **18**:1112–1126.
Hubley R, Finn RD, Clements J, Eddy SR, Jones TA, Bao W, Smit AFA, Wheeler TJ. 2016. The Dfam database of repetitive DNA families. *Nucleic Acids Res* **44**:D81–9.
Ivancevic AM, Kortschak RD, Bertozzi T, Adelson DL. 2016. LINEs between Species: Evolutionary Dynamics of LINE-1 Retrotransposons across the Eukaryotic Tree of Life. *Genome Biol Evol* **8**:3301–3322.
Jaillon O, Aury J-M, Brunet F, Petit J-L, Stange-Thomann N, Mauceli E, Bouneau L, Fischer C, Ozouf-Costaz C, Bernot A, Nicaud S, Jaffe D, Fisher S, Lutfalla G, Dossat C, Segurens B, Dasilva C, Salanoubat M, Levy M, Boudet N, Castellano S, Anthouard V, Jubin C, Castelli V, Katinka M, Vacherie B, Biémont C, Skalli Z, Cattolico L, Poulain J, De Berardinis V, Cruaud C, Duprat S, Brottier P, Coutanceau J-P, Gouzy J, Parra G, Lardier G, Chapple C, McKernan KJ, McEwan P, Bosak S, Kellis M, Volff J-N, Guigó R, Zody MC, Mesirov J, Lindblad-Toh K, Birren B, Nusbaum C, Kahn D, Robinson-Rechavi M, Laudet V, Schachter V, Quétier F, Saurin W, Scarpelli C, Wincker P, Lander ES, Weissenbach J, Roest Crollius H. 2004. Genome duplication in the teleost fish Tetraodon nigroviridis reveals the early vertebrate proto-karyotype. *Nature* **431**:946–957.
Jiang N, Bowman M, Childs K. 2016. Repeat Library Construction-Advanced. *MAKER Wiki*. http://weatherby.genetics.utah.edu/MAKER/wiki/index.php/Repeat_Library_Construction-Advanced
Keestra-Gounder AM, Tsolis RM. 2017. NOD1 and NOD2: Beyond Peptidoglycan Sensing. *Trends Immunol* **38**:758–767.

Kim D, Langmead B, Salzberg SL. 2015. HISAT: a fast spliced aligner with low memory requirements. *Nat Methods* **12**:357–360.

Korithoski B, Kolaczkowski O, Mukherjee K, Kola R, Earl C, Kolaczkowski B. 2015. Evolution of a Novel Antiviral Immune-Signaling Interaction by Partial-Gene Duplication. *PLoS One* **10**:e0137276.

Laing KJ, Purcell MK, Winton JR, Hansen JD. 2008. A genomic view of the NOD-like receptor family in teleost fish: identification of a novel NLR subfamily in zebrafish. *BMC Evol Biol* **8**:42.

Latz E, Xiao TS, Stutz A. 2013. Activation and regulation of the inflammasomes. *Nat Rev Immunol* **13**:397–411.

Leulier F, Lemaitre B. 2008. Toll-like receptors--taking an evolutionary approach. *Nat Rev Genet* **9**:165–178.

Malmstrøm M, Matschiner M, Tørresen OK, Star B, Snipen LG, Hansen TF, Baalsrud HT, Nederbragt AJ, Hanel R, Salzburger W, Stenseth NC, Jakobsen KS, Jentoft S. 2016. Evolution of the immune system influences speciation rates in teleost fishes. *Nat Genet*. doi:10.1038/ng.3645

Manousaki T, Feiner N, Begemann G, Meyer A, Kuraku S. 2011. Co-orthology of P ax4 and P ax6 to the fly eyeless gene: molecular phylogenetic, comparative genomic, and embryological analyses. *Evol Dev* **13**:448–459.

Marra NJ, Stanhope MJ, Jue NK, Wang M, Sun Q, Bitar PP, Richards VP, Komissarov A, Rayko M, Kliver S, Stanhope BJ, Winkler C, O'Brien SJ, Antunes A, Jorgensen S, Shivji MS. 2019. White shark genome reveals ancient elasmobranch adaptations associated with wound healing and the maintenance of genome stability. *Proc Natl Acad Sci U S A* 201819778.

Nei M, Gu X, Sitnikova T. 1997. Evolution by the birth-and-death process in multigene families of the vertebrate immune system. *Proc Natl Acad Sci U S A* **94**:7799–7806.

Nei M, Rooney AP. 2005. Concerted and birth-and-death evolution of multigene families. *Annu Rev Genet* **39**:121–152.

Pertea M, Pertea GM, Antonescu CM, Chang T-C, Mendell JT, Salzberg SL. 2015. StringTie enables improved reconstruction of a transcriptome from RNA-seq reads. *Nat Biotechnol* **33**:290–295.

Peterson KJ, Sperling EA. 2007. Poriferan ANTP genes: primitively simple or secondarily reduced? *Evol Dev* **9**:405–408.

Proell M, Riedl SJ, Fritz JH, Rojas AM, Schwarzenbacher R. 2008. The Nod-like receptor (NLR) family: a tale of similarities and differences. *PLoS One* **3**:e2119.

Puttick MN, Thomas GH. 2015. Fossils and living taxa agree on patterns of body mass evolution: a case study with Afrotheria. *Proc Biol Sci* **282**:20152023.

Rabosky DL. 2014. Automatic Detection of Key Innovations, Rate Shifts, and Diversity-Dependence on Phylogenetic Trees. *PLoS One* **9**:e89543.

Rabosky DL, Grundler M, Anderson C, Title P, Shi JJ, Brown JW, Huang H, Larson JG. 2014. BAMMtools: an R package for the analysis of evolutionary dynamics on phylogenetic trees. *Methods Ecol Evol* **5**:701–707.

Rast JP, Smith LC, Loza-Coll M, Hibino T, Litman GW. 2006. Genomic insights into the immune system of the sea urchin. *Science* **314**:952–956.

Read TD, Petit RA 3rd, Joseph SJ, Alam MT, Weil MR, Ahmad M, Bhimani R, Vuong JS, Haase CP, Webb DH, Tan M, Dove ADM. 2017. Draft sequencing and assembly of the genome of the world's largest fish, the whale shark: *Rhincodon typus* Smith 1828. *BMC Genomics* **18**:532.

Redmond AK, Macqueen DJ, Dooley H. 2018. Phylotranscriptomics suggests the jawed vertebrate ancestor could generate diverse helper and regulatory T cell subsets. *BMC Evol Biol* **18**:169.

Redmond AK, Pettinello R, Dooley H. 2017. Outgroup, alignment and modelling improvements indicate that two TNFSF13-like genes existed in the vertebrate ancestor. *Immunogenetics* **69**:187–192.

Redmond AK, Zou J, Secombes CJ, Macqueen DJ, Dooley H. 2019. Discovery of All Three Types in Cartilaginous Fishes Enables Phylogenetic Resolution of the Origins and Evolution of Interferons. *Front Immunol* **10**:1558.

Roach JC, Glusman G, Rowen L, Kaur A, Purcell MK, Smith KD, Hood LE, Aderem A. 2005. The evolution of vertebrate Toll-like receptors. *Proc Natl Acad Sci U S A* **102**:9577–9582.

Schroder K, Tschopp J. 2010. The inflammasomes. *Cell* **140**:821–832.

Shao F, Wang J, Xu H, Peng Z. 2018. FishTEDB: a collective database of transposable elements identified in the complete genomes of fish. *Database* **2018**. doi:10.1093/database/bax106

She R, Chu JS-C, Uyar B, Wang J, Wang K, Chen N. 2011. genBlastG: using BLAST searches to build homologous gene models. *Bioinformatics* **27**:2141–2143.

She R, Chu JS-C, Wang K, Pei J, Chen N. 2009. GenBlastA: enabling BLAST to identify homologous gene sequences. *Genome Res* **19**:143–149.

Slater GJ, Goldbogen JA, Pyenson ND. 2017. Independent evolution of baleen whale gigantism linked to Plio-Pleistocene ocean dynamics. *Proc Biol Sci* **284**. doi:10.1098/rspb.2017.0546

Smit AFA, Hubley R. 2008. RepeatModeler Open-1.0. *Available fom http://www repeatmasker org*.

Smith JJ, Kuraku S, Holt C, Sauka-Spengler T, Jiang N, Campbell MS, Yandell MD, Manousaki T, Meyer A, Bloom OE, Morgan JR, Buxbaum JD, Sachidanandam R, Sims C, Garruss AS, Cook M, Krumlauf R, Wiedemann LM, Sower SA, Decatur WA, Hall JA, Amemiya CT, Saha NR, Buckley KM, Rast JP, Das S, Hirano M, McCurley N, Guo P, Rohner N, Tabin CJ, Piccinelli P, Elgar G, Ruffier M, Aken BL, Searle SMJ, Muffato M, Pignatelli M, Herrero J, Jones M, Brown CT, Chung-Davidson Y-W, Nanlohy KG, Libants SV, Yeh C-Y, McCauley DW, Langeland JA, Pancer Z, Fritzsch B, de Jong PJ, Zhu B, Fulton LL, Theising B, Flicek P, Bronner ME, Warren WC, Clifton SW, Wilson RK, Li W. 2013. Sequencing of the sea lamprey (Petromyzon marinus) genome provides insights into vertebrate evolution. *Nat Genet* **45**:415–21, 421e1–2.

Star B, Nederbragt AJ, Jentoft S, Grimholt U, Malmstrøm M, Gregers TF, Rounge TB, Paulsen J, Solbakken MH, Sharma A, Wetten OF, Lanzén A, Winer R, Knight J, Vogel J-H, Aken B, Andersen Ø, Lagesen K, Tooming-Klunderud A, Edvardsen RB, Tina KG, Espelund M, Nepal C, Previti C, Karlsen BO, Moum T, Skage M, Berg PR, Gjøen T, Kuhl H, Thorsen J, Malde K, Reinhardt R, Du L, Johansen SD, Searle S, Lien S, Nilsen F, Jonassen I, Omholt SW, Stenseth NC, Jakobsen KS. 2011. The genome sequence of Atlantic cod reveals a unique immune system. *Nature* **477**:207–210.

Steinbiss S, Willhoeft U, Gremme G, Kurtz S. 2009. Fine-grained annotation and classification of de novo predicted LTR retrotransposons. *Nucleic Acids Res* **37**:7002–7013.

Stein RW, Mull CG, Kuhn TS, Aschliman NC, Davidson LNK, Joy JB, Smith GJ, Dulvy NK, Mooers AO. 2018. Global priorities for conserving the evolutionary history of sharks, rays and chimaeras. *Nat Ecol Evol* **2**:288–298.

Thomas JH. 2005. Rapid Birth-Death Evolution Specific to Xenobiotic Cytochrome P450 Genes in Vertebrates. *PLoS Genet* **preprint**:e67.

Ting JP-Y, Lovering RC, Alnemri ES, Bertin J, Boss JM, Davis BK, Flavell RA, Girardin SE, Godzik A, Harton JA, Hoffman HM, Hugot J-P, Inohara N, Mackenzie A, Maltais LJ, Nunez G, Ogura Y, Otten LA, Philpott D, Reed JC, Reith W, Schreiber S, Steimle V, Ward PA. 2008. The NLR gene family: a standard nomenclature. *Immunity* **28**:285–287.

Venkatesh B, Lee AP, Ravi V, Maurya AK, Lian MM, Swann JB, Ohta Y, Flajnik MF, Sutoh Y, Kasahara M, Hoon S, Gangu V, Roy SW, Irimia M, Korzh V, Kondrychyn I, Lim ZW, Tay B-H, Tohari S, Kong KW, Ho S, Lorente-Galdos B, Quilez J, Marques-Bonet T, Raney BJ, Ingham PW, Tay A, Hillier LW, Minx P, Boehm T, Wilson RK, Brenner S, Warren WC.

2014. Elephant shark genome provides unique insights into gnathostome evolution. *Nature* **505**:174–179.
Vidya MK, Kumar VG, Sejian V, Bagath M, Krishnan G, Bhatta R. 2018. Toll-like receptors: Significance, ligands, signaling pathways, and functions in mammals. *Int Rev Immunol* **37**:20–36.
Wang J, Zhang Z, Liu J, Zhao J, Yin D. 2016. Ectodomain Architecture Affects Sequence and Functional Evolution of Vertebrate Toll-like Receptors. *Sci Rep* **6**:26705.
Wcisel DJ, Ota T, Litman GW, Yoder JA. 2017. Spotted Gar and the Evolution of Innate Immune Receptors. *J Exp Zool B Mol Dev Evol* **328**:666–684.
Weber JA, Park SG, Luria V, Jeon S, Kim H-M, Jeon Y, Bhak Y, Jun JH, Kim SW, Hong WH, Lee S, Cho YS, Karger A, Cain JW, Manica A, Kim S, Kim J-H, Edwards JS, Bhak J, Church GM. 2020. The whale shark genome reveals how genomic and physiological properties scale with body size. *Proc Natl Acad Sci U S A*. doi:10.1073/pnas.1922576117
Xu L, Yu D, Fan Y, Peng L, Wu Y, Yao Y-G. 2016. Loss of RIG-I leads to a functional replacement with MDA5 in the Chinese tree shrew. *Proc Natl Acad Sci U S A* **113**:10950–10955.
Yuen B, Bayes JM, Degnan SM. 2014. The characterization of sponge NLRs provides insight into the origin and evolution of this innate immune gene family in animals. *Mol Biol Evol* **31**:106–120.

Supplementary File 1. Repeat library annotated using the MAKER repeat annotation pipeline (FASTA). Repeat classification of each repeat sequence follows a "#" delimiter.
Supplementary File 2. Whale shark transcriptome annotation based on StringTie (GFF).
Supplementary File 3. Putative conserved vertebrate genes absent from the whale shark RefSeq annotation that were annotated using genBlast (GFF).
Supplementary File 4. Putative conserved vertebrate genes absent from the *Callorhinchus* RefSeq annotation that were annotated using genBlast. Annotations are for the GCF_000165045.1 genome assembly (GFF).
Supplementary File 5. Orthogroup assignment by OrthoFinder of chordate proteins (CSV).
Supplementary File 6. GO and PFfam annotations of orthogroups assigned by KinFin (tab-delimited table TSV).
Supplementary File 7. Human gene names of human orthologs assigned to each orthogroup (TXT).
Supplementary File 8. Species included and excluded for TLR analysis from Wang *et al.* 2016 dataset (XLSX).
Supplementary File 9. CAFE output for rates of gene duplication and loss of vertebrate orthogroups computed under a single global rate of gene duplication and loss for orthogroups (TXT).
Supplementary File 10. Scripts used for performing analyses to annotate repetitive sequences, assess gene family gain and loss and enrichment of gene family functional annotations, compare gene family assignment to known ohnologs, compare rates of substitution across the phylogeny using LINTRE and PAML, assess the rate of body size evolution across cartilaginous fishes compared to the whale shark, and summarize rates of gene family size evolution and enrichment of functional annotations and cancer-related function (ZIP).
