## Supplementary tables 6-11 for "The whale shark genome reveals patterns of vertebrate gene family evolution"

Supplementary table 6. Significantly-enriched functional and domain terms identified in novel gene families (orthogroups, Supplemental File 5) gained in the MRCA of Olfactores. n refers to the number of these gene families with that function gained. p refers to uncorrected p-values for Fisher's exact test, adj.p refers to the adjusted p-value for multiple testing (See Methods). See Supplementary File 7 for specific assignments of human gene names to each orthogroup.

| domain id | domain description | n | p | estimate odds.ratio | adj.p | orthogroups | human gene names |
|---|---|---|---|---|---|---|---|
| PF12796 | Ankyrin repeats (3 copies) | 34 | 5.21E-10 | 4.22448135 | 1.79E-06 | OG0000478, OG0000567, OG0000584, OG0001171, OG0001315, OG0001439, OG0001683, OG0001750, OG0001936, OG0002086, OG0002201, OG0002362, OG0002408, OG0002434, OG0002458, OG0003117, OG0003477, OG0004211, OG0005945, OG0006086, OG0006976, OG0007434, OG0007820, OG0008219, OG0008248, OG0008383, OG0009058, OG0009608, OG0010023, OG0010058, OG0010235, OG0010681, OG0011197, OG0011811 | ANK1, ANK3, ANK2, ANKRD52, ANKRD44, ANKRD28, ANKRD18B, POTED, POTEH, ANKRD20A4P, POTEC, ANKRD30B, CCDC144A, ANKRD20A2P, ANKRD18A, ANKRD30BL, ANKRD7, ANKRD36C, ANKRD26, AC098850.3, POTEM, POTEG, POTEB2, ANKRD62, ANKRD20A1, ANKRD36B, AC136352.4, BX072566.1, POTEB3, ANKRD20A3P, FAM95C, POTEA, ASB5, ASB9, ASB11, TANC1, TANC2, ANKS1A, ANKS1B, NFKBIE, EHMT2, EHMT1, ASB15, ASB14, BCORL1, BCOR, ANKS4B, USH1G, TNKS, TNKS2, ANKRD12, ANKRD11, NFKB1, NFKB2, ZDHHC13, ZDHHC17, ANKRA2, RFXANK, ANKRD17, ANKHD1, ANKHD1-EIF4EBP3, ANKRD23, ANKRD1, ANKEF1, ANKFY1, PSMD10, FEM1B, MPHOSPH8, ANKS3, ANKRD16, HECTD1, SNCAIP, ANKRD39, ANKRD31, BCL3, ANKRD42, ANKRD66 |
| PF13637 | Ankyrin repeats (many copies) | 7 | 6.75E-06 | 22.1412282 | 0.01383678 | OG0000478, OG0000567, OG0002362, OG0006086, OG0008248, OG0010235, OG0011197 | ANK1, ANK3, ANK2, ANKRD52, ANKRD44, ANKRD28, TNKS, TNKS2, ANKFY1, ANKRD16, ANKRD42 |
| PF00029 | Connexin | 5 | 8.03E-06 | Inf | 0.01383678 | OG0000128, OG0000926, OG0002122, OG0006602, OG0010862 | GJB6, GJB5, GJB4, GJB1, GJB3, GJB2, GJB7, GJC1, GJC2, GJD2, GJD4 |

Supplementary table 7. Significantly-enriched functional and domain terms identified in novel gene families (orthogroups) gained in the MRCA of vertebrates. n refers to the number of these gene families with that function gained. p.value refers to uncorrected p-values for Fisher's exact test, adj.p refers to the adjusted p-value for multiple testing (See Methods). See Supplementary File 7 for specific assignments of human gene names to each orthogroup.

| domain id | domain description | n | p | estimate odds.ratio | adj.p | orthogroups | human gene names |
|---|---|---|---|---|---|---|---|
| PF00001 | 7 | 34 | 3.23E-13 | 5.63990176 | 2.27E-09 | OG0000103, OG0000168, OG0000181, OG0000479, | CCR2, CCR5, CCR1, XCR1, CCR8, CCR4, CX3CR1, ACKR2, CCR3, F2RL3, |

| | | | | | | | |
|---|---|---|---|---|---|---|---|
| | transmembrane receptor (rhodopsin family) | | | | | OG0000650, OG0001472, OG0001557, OG0001638, OG0001664, OG0001826, OG0001908, OG0001960, OG0002235, OG0002331, OG0002573, OG0002711, OG0002969, OG0003696, OG0004011, OG0004310, OG0005633, OG0007575, OG0007780, OG0007904, OG0008315, OG0009052, OG0009906, OG0010481, OG0010743, OG0011476, OG0012139, OG0012756, OG0014852, OG0018208 | F2RL2, F2RL1, F2R, MC3R, MC4R, MC5R, MC2R, MC1R, FFAR2, FFAR1, FFAR3, GPR42, GPR65, GPR4, GPR68, TRHR, GPR85, GPR27, PTGER2, PTGIR, PTGDR, GPR37, GPR37L1, DRD3, DRD2, CYSLTR2, CYSLTR1, P2RY1, LPAR6, LPAR4, PTGER1, PTGFR, GPR26, GPR78, BDKRB1, BDKRB2, GPR83, GPR34, NPBWR1, NPBWR2, CXCR4, GPR146, GPR18, ACKR3, OXGR1, GPR176, GPR148, NPY5R |
| GO:0016021 | integral component of membrane | 71 | 2.64E-09 | 2.40370814 | 9.27E-06 | OG0000004, OG0000103, OG0000168, OG0000479, OG0000492, OG0000588, OG0000650, OG0000656, OG0000936, OG0001059, OG0001391, OG0001453, OG0001472, OG0001557, OG0001638, OG0001664, OG0001826, OG0001908, OG0001960, OG0002098, OG0002235, OG0002268, OG0002331, OG0002573, OG0002711, OG0002817, OG0002969, OG0003542, OG0003696, OG0003922, OG0004011, OG0004310, OG0004658, OG0004879, OG0005633, OG0007456, OG0007575, OG0007780, OG0007781, OG0007793, OG0007904, OG0008315, OG0008510, OG0008604, OG0008726, OG0008752, OG0009035, OG0009052, OG0009101, OG0009113, OG0009804, OG0009906, OG0010481, OG0010743, OG0010788, OG0012139, OG0012532, OG0012756, OG0013601, OG0014056, OG0014152, OG0014630, OG0014753, OG0014758, OG0014852, OG0017502, OG0018208, OG0018213, OG0018286, OG0018298, OG0018303 | OR51B5, OR52K1, OR52A5, OR52I2, OR52K2, OR52E4, OR51V1, OR52E2, OR51L1, OR51G2, OR52N2, OR52N1, OR52N5, OR52N4, OR52D1, OR52L1, OR51M1, OR52A1, OR52B6, OR51I2, OR51D1, OR52M1, OR52E6, OR51I1, OR51B6, OR51B4, OR52J3, OR51A2, OR51F1, OR51E2, OR51E1, OR52I1, OR52B2, OR52E8, OR52E1, OR52E5, OR52R1, OR51B2, OR52B4, OR51G1, CCR2, CCR5, CCR1, XCR1, CCR8, CCR4, CX3CR1, ACKR2, CCR3, F2RL3, F2RL2, F2RL1, F2R, MC3R, MC4R, MC5R, MC2R, MC1R, ADGRF5, ADGRF2, ADGRF3, ADGRF4, ADGRF1, TM4SF5, TM4SF1, TM4SF4, FFAR2, FFAR1, FFAR3, GPR42, GABRG1, GABRE, GABRG2, GABRG3, PRPH2, ROM1, SLC4A5, SLC4A4, SLC4A9, CHRND, SYNDIG1L, SYNDIG1, TMEM91, GPR65, GPR4, GPR68, TRHR, GPR85, GPR27, PTGER2, PTGIR, PTGDR, GPR37, GPR37L1, DRD3, DRD2, CYSLTR2, CYSLTR1, SLC1A4, SLC1A5, P2RY1, CACNG1, CACNG6, LPAR6, LPAR4, PTGER1, PTGFR, GPR26, GPR78, CHRNB3, CHRNA5, BDKRB1, BDKRB2, MGST2, LTC4S, GPR83, UPK1B, UPK1A, GPR34, NPBWR1, NPBWR2, XKR8, CLDN10, CXCR4, GPR146, GPR18, CFTR, OCSTAMP, ACKR3, OXGR1, CHST10, KCNJ13, NIPA1, XKR9, GPR176, CLDN18, SLC29A3, MMGT1, GPR148, NPY5R |
| PF07686 | Immunoglobulin V-set domain | 9 | 1.65E-08 | 31.2006635 | 3.85E-05 | OG0001162, OG0001753, OG0003241, OG0004367, OG0004811, OG0006579, OG0009319, OG0010863, OG0014243 | CXADR, CLMP, AC136352.5, SCN3B, SCN1B, SIGLEC15, HEPACAM, MXRA8, VSTM2L, SCN2B |
| PF01391 | Collagen triple helix repeat (20 copies) | 14 | 2.41E-08 | 9.29967479 | 4.22E-05 | OG0001408, OG0001971, OG0003316, OG0003349, OG0003947, OG0004089, OG0004578, OG0004744, OG0004785, OG0004923, OG0008036, OG0008695, OG0012951, OG0015390 | COL23A1, COL13A1, COL25A1, C1QB, C1QC, C1QA, C1QTNF7, C1QTNF2, ADIPOQ, C1QTNF9, C1QTNF9B, COL7A1, COL14A1, COL6A2, COL12A1, EMILIN1, COL6A1, C1QTNF5, SCARA3 |
| PF00100 | Zona pellucida-like domain | 7 | 1.47E-05 | 13.8275909 | 0.02057389 | OG0000407, OG0000458, OG0001540, OG0001842, OG0005017, OG0010090, OG0014060 | ZP1, ZP4, ZP2, ZP3, ZPLD1, TECTA, TECTB |
| PF00096 | Zinc finger, C2H2 type | 27 | 2.48E-05 | 2.74829863 | 0.02483901 | OG0001984, OG0002987, OG0003154, OG0003192, OG0003554, OG0004208, OG0004368, OG0004461, OG0004939, OG0006971, OG0007186, OG0007392, | ZNF281, ZNF148, ZNF837, HIC1, HIC2, SP5, ZBTB16, SP3, ZXDA, ZXDB, ZXDC, GLIS2, ZBTB49, PRDM4, ZBTB40, ZBTB2, PRDM15, KLF3, ZBTB14, ZNF516, ZNF653, AC008481.3, ZBTB21, GLI4, ZNF672, SALL2, ZNF275, |

| | | | | | | | |
|---|---|---|---|---|---|---|---|
| | | | | | | OG0007519, OG0007545, OG0007594, OG0008196, OG0009061, OG0009170, OG0009438, OG0010626, OG0011065, OG0011584, OG0011821, OG0011965, OG0012017, OG0013057, OG0014518 | ZNF771, ZFP3 |
| PF00092 | von Willebrand factor type A domain | 9 | 2.24E-05 | 7.794886 | 0.02483901 | OG0000383, OG0002543, OG0003947, OG0004089, OG0004578, OG0004744, OG0004923, OG0005486, OG0012771 | ITGA1, ITGA2, ITGA10, ITGA11, VIT, COL7A1, COL14A1, COL6A2, COL12A1, COL6A1, VWA2 |
| GO:0007186 | G protein-coupled receptor signaling pathway | 14 | 2.99E-05 | 4.32999648 | 0.02619907 | OG0000004, OG0000492, OG0001638, OG0002083, OG0002946, OG0007389, OG0007904, OG0009052, OG0010743, OG0010869, OG0014051, OG0014753, OG0014852, OG0018208 | OR51B5, OR52K1, OR52A5, OR52I2, OR52K2, OR52E4, OR51V1, OR52E2, OR51L1, OR51G2, OR52N2, OR52N1, OR52N5, OR52N4, OR52D1, OR52L1, OR51M1, OR52A1, OR52B6, OR51I2, OR51D1, OR52M1, OR52E6, OR51I1, OR51B6, OR51B4, OR52J3, OR51A2, OR51F1, OR51E2, OR51E1, OR52I1, OR52B2, OR52E8, OR52E1, OR52E5, OR52R1, OR51B2, OR52B4, OR51G1, ADGRF5, ADGRF2, ADGRF3, ADGRF4, ADGRF1, GPR85, GPR27, RGS9, RGS11, GNG11, GNGT1, GNGT2, GNG13, ACKR3, GPR176, NPY5R |

Supplementary table 8. Significantly-enriched functional and domain terms identified in novel gene families (orthogroups) gained in the MRCA of gnathostomes. n refers to the number of these gene families with that function gained. p.value refers to uncorrected p-values for Fisher's exact test, adj.p refers to the adjusted p-value for multiple testing (See Methods). See Supplementary File 7 for specific assignments of human gene names to each orthogroup.

| domain id | domain description | n | p | estimate odds.ratio | adj.p | orthogroups | human gene names |
|---|---|---|---|---|---|---|---|
| PF07686 | Immunoglobulin V-set domain | 65 | 3.69E-31 | 18.489222 | 9.55E-28 | OG0000002, OG0000006, OG0000013, OG0000021, OG0000063, OG0000080, OG0000087, OG0000686, OG0001164, OG0001304, OG0001378, OG0002194, OG0002715, OG0002882, OG0003641, OG0003774, OG0003966, OG0004153, OG0004170, OG0004171, OG0004548, OG0004595, OG0004620, OG0004813, OG0004900, OG0005006, OG0005524, OG0006062, OG0006725, OG0008000, OG0009114, OG0009269, OG0009334, OG0009420, OG0009442, OG0009915, OG0010028, OG0010095, OG0010266, OG0010296, OG0010309, OG0010333, OG0010364, OG0010387, OG0010437, OG0010613, OG0010773, OG0010792, OG0010825, OG0010921, OG0011010, OG0011128, | VPREB3, IGKV4-1, IGKV5-2, IGKV3-7, IGKV3-15, IGKV6-21, IGKV1D-33, IGKV2D-26, IGKV3D-20, IGKV6D-41, IGKV3D-11, IGLV4-69, IGLV8-61, IGLV4-60, IGLV6-57, IGLV11-55, IGLV5-52, IGLV1-51, IGLV1-50, IGLV5-48, IGLV1-47, IGLV7-46, IGLV5-45, IGLV1-44, IGLV7-43, IGLV1-40, IGLV5-37, IGLV1-36, IGLV2-33, IGLV3-32, IGLV3-27, IGLV3-25, IGLV2-23, IGLV3-22, IGLV3-21, IGLV3-19, IGLV2-14, IGLV2-11, IGLV3-9, IGLV4-3, IGLV3-1, VPREB1, IGKV2D-28, IGKV3D-7, IGKV3D-15, IGKV1D-39, IGKV2D-40, IGKV6D-21, IGLV9-49, IGKV2D-24, IGKV1-16, IGKV1-37, IGKV2D-29, IGKV1D-43, IGKV2-30, IGKV1D-16, IGKV1D-17, IGKV1-17, IGKV3-20, IGKV1-27, IGKV2D-30, IGKV1-39, IGKV1-8, IGKV2-24, IGKV2-28, IGKV1-9, IGKV1-33, IGKV3-11, IGKV1D-8, IGKV1-6, IGKV1-5, IGKV1-12, IGKV1D-37, IGKV3OR2-268, IGKV1OR2-108, IGKV2-40, IGKV1D-13, IGKV1D-12, IGLV2-8, AC145029.2, AC009286.3, IGHV6-1, IGHV1-2, |

| | | | | | | | |
|---|---|---|---|---|---|---|---|
| | | | | | | OG0011163, OG0011212, OG0011561, OG0012291, OG0012389, OG0012395, OG0013259, OG0013411, OG0013573, OG0013686, OG0014061, OG0014552, OG0015823 | IGHV1-3, IGHV2-5, IGHV3-7, IGHV3-11, IGHV3-13, IGHV3-15, IGHV3-16, IGHV1-18, IGHV3-20, IGHV3-21, IGHV3-23, IGHV1-24, IGHV2-26, IGHV4-28, IGHV3-33, IGHV4-34, IGHV3-35, IGHV3-38, IGHV4-39, IGHV1-45, IGHV1-46, IGHV3-48, IGHV3-49, IGHV5-51, IGHV3-53, IGHV1-58, IGHV4-59, IGHV4-61, IGHV3-66, IGHV1-69, IGHV2-70D, IGHV3-73, IGHV7-81, IGHV3-74, IGHV4-31, IGHV3-43, IGHV3-64, IGHV4-4, IGHV3-72, IGHV4OR15-8, IGHV3OR16-12, AC136428.1, IGHV1OR15-1, IGHV3OR16-9, IGHV2OR16-5, IGHV3OR15-7, IGHV3-30, IGHV1OR15-9, IGHV3OR16-13, IGHV3OR16-10, IGHV3OR16-8, AC233755.1, AC141272.1, IGHV1-69-2, AC136616.2, AC135068.2, AC233755.2, IGHV1OR21-1, IGHV1-69D, IGHV2-70, AC135068.8, AC234301.4, IGHV7-4-1, AC234301.1, IGHV5-10-1, IGHV3-64D, AC244226.1, AC247036.3, AC247036.4, AC234301.3, BTN1A1, BTN3A3, MOG, BTN3A1, BTN2A1, BTNL9, BTNL3, BTNL8, BTN3A2, BTN2A2, HHLA2, VTCN1, BTNL2, AC226007.1, TRAV2, TRAV9-1, TRAV8-2, TRAV8-3, TRAV8-1, TRAV3, TRAV8-4, TRAV14DV4, TRAV8-7, TRAV8-6, TRAV38-1, TRAV26-2, TRAV16, TRAV4, TRAV18, TRDV3, TRAV9-2, TRDV1, TRAV19, TRAV38-2DV8, TRAV26-1, TRAV40, CD300LG, CD300C, TREML4, CD300A, TMIGD3, NCR2, CD300LD, CD300E, CD300LB, CD300LF, TRBV6-1, TRBV4-1, TRBV6-4, TRBV7-3, TRBV5-3, TRBV9, TRBV10-1, TRBV11-1, TRBV6-5, TRBV7-4, TRBV6-6, TRBV5-5, TRBV6-7, TRBV7-6, TRBV5-6, TRBV6-8, TRBV7-7, TRBV5-7, TRBV5-1, TRBV3-1, TRBV4-2, TRBV19, TRBV23-1, TRBV24-1, TRBV25-1, TRBV27, TRBV28, TRBV2, TRBV10-2, TRBV5-4, TRBV7-1, TRBV13, TRBV14, TRBV12-3, TRBV7-9, TRBV16, TRBV10-3, TRBV12-5, TRBV11-3, TRBV12-4, TRBV15, TRBV17, TRBV18, AC229888.1, AC234635.1, TRBV7-2, AC233282.1, AC233282.2, AC234635.3, TRBV6-2, TRAV12-2, TRAV6, TRAV13-2, TRAV10, TRAV13-1, TRAV7, TRAV12-1, TRAV5, TRAV36DV7, TRAV39, TRAV41, TRAV23DV6, TRAV30, TRAV27, TRAV12-3, TRAV24, TRAV34, TRAV20, TRAV17, TRAV25, TRAV29DV5, TRAV22, TRAV21, CD33, SIGLEC12, SIGLEC8, SIGLEC7, SIGLEC10, SIGLEC14, AC011452.2, SIGLEC6, SIGLEC11, SIGLEC9, AC018755.2, SIGLEC5, TIMD4, HAVCR2, HAVCR1, MPZL2, MPZL3, MPZL1, TRGV11, TRGV10, TRGV8, TRGV5, TRGV4, TRGV3, TRGV1, TRGV9, TRGV2, AC242528.1, CD200, CD101, IGSF3, CD274, PDCD1LG2, NCR3LG1, NCR3, TAPBPL, CD8A, TRBV30, CD276, CD200R1, CD200R1L, VSTM4, SCN4B, ESAM, PTGFRN, VSIR, TRBV20OR9-2, TRBV20-1, TRBV29-1, FAM187A, CD80, VSTM2A, CD226, SECTM1, CD7, IGSF8, CRTAM, ICOSLG, FP565260.3, MPZ, VSTM2B, VSIG4, PILRA, IGSF6, TIGIT, TMIGD2, CD8B, CD86, JAML |
| PF07654 | Immunoglob ulin C1-set | 16 | 2.25E-11 | Inf | 2.91E-08 | OG0000019, OG0000091, OG0000143, OG0000284, OG0000319, OG0003641, OG0004153, OG0004171, | HLA-F, AZGP1, HLA-B, MR1, HLA-E, HLA-A, HLA-G, HLA-C, HFE, IGLL1, IGKC, IGLC1, IGLC2, IGLC3, IGLC7, IGLJ1, IGLL5, HLA-DRB3, HLA-DQB2, |

| | | | | | | | |
|---|---|---|---|---|---|---|---|
| | domain | | | | | OG0004412, OG0004971, OG0005377, OG0008912, OG0011212, OG0011236, OG0011551, OG0013594 | HLA-DRB1, HLA-DQB1, HLA-DRB5, HLA-DPB1, HLA-DOB, HLA-DRB4, HLA-DOA, HLA-DQA2, HLA-DQA1, HLA-DPA1, HLA-DRA, AL662796.1, IGHA2, IGHE, IGHG4, IGHG2, IGHA1, IGHG1, IGHG3, IGHM, NCR3LG1, TAPBPL, B2M, TRBC2, TRBC1, TAPBP, TRGC1, TRGC2, TRDC |
| GO:0005576 | extracellular region | 52 | 1.68E-08 | 3.024175 | 1.45E-05 | OG0000050, OG0000094, OG0000095, OG0000266, OG0001580, OG0001667, OG0002374, OG0002625, OG0002696, OG0002893, OG0003159, OG0003363, OG0003573, OG0003885, OG0004351, OG0004652, OG0004775, OG0005227, OG0006012, OG0006200, OG0006436, OG0006890, OG0007402, OG0008260, OG0008878, OG0009042, OG0009491, OG0009542, OG0009678, OG0009714, OG0009774, OG0010261, OG0010339, OG0010419, OG0010448, OG0010742, OG0010837, OG0010904, OG0010911, OG0011002, OG0011097, OG0011125, OG0011213, OG0011242, OG0012322, OG0012386, OG0013115, OG0013326, OG0013403, OG0016060, OG0016519, OG0017497 | CCL26, CCL22, CCL17, CCL24, CCL2, CCL13, CCL21, CCL11, CCL20, CCL7, CCL19, CCL8, CCL15, CCL3, CCL16, CCL4, CCL4L2, CCL18, CCL3L3, CCL3L1, CCL23, CCL15-CCL14, CCL14, CCL5, IFNA7, IFNA6, IFNA1, IFNK, IFNA8, IFNA2, IFNA16, IFNA10, IFNA14, IFNA21, IFNW1, IFNB1, IFNA17, IFNA4, IFNE, IFNA13, IFNA5, CXCL6, PF4V1, CXCL13, CXCL3, CXCL5, PPBP, PF4, CXCL10, CXCL8, CXCL11, CXCL9, CXCL1, CXCL2, APOL5, APOL1, APOLD1, APOL4, APOL3, APOL6, APOL2, APOE, APOA4, PRL, IGFBP4, IGFBP1, NPPC, SST, PRRG3, PRRG1, CRH, UCN, ASIP, PRRG2, PRRG4, PTHLH, SPP2, CXCL12, GUCA2A, GUCA2B, ADCYAP1, CCK, UCN3, EPO, C5, ADM, IAPP, IL12A, ECM1, VIP, CCL25, CCL27, CCL28, CGA, CXCL14, LIF, IL22, PTH2, NTS, IL15, LEP, GNRH1, IFNG, MLN, HAMP, APOA1 |
| GO:0006955 | immune response | 21 | 4.66E-08 | 7.64906736 | 3.62E-05 | OG0000050, OG0000095, OG0000143, OG0000284, OG0004652, OG0007971, OG0008878, OG0009542, OG0009678, OG0009774, OG0010261, OG0010536, OG0010695, OG0010721, OG0010722, OG0010742, OG0011002, OG0011242, OG0011317, OG0011472, OG0012322 | CCL26, CCL22, CCL17, CCL24, CCL2, CCL13, CCL21, CCL11, CCL20, CCL7, CCL19, CCL8, CCL15, CCL3, CCL16, CCL4, CCL4L2, CCL18, CCL3L3, CCL3L1, CCL23, CCL15-CCL14, CCL14, CCL5, CXCL6, PF4V1, CXCL13, CXCL3, CXCL5, PPBP, PF4, CXCL10, CXCL8, CXCL11, CXCL9, CXCL1, CXCL2, HLA-DRB3, HLA-DQB2, HLA-DRB1, HLA-DQB1, HLA-DRB5, HLA-DPB1, HLA-DOB, HLA-DRB4, HLA-DOA, HLA-DQA2, HLA-DQA1, HLA-DPA1, HLA-DRA, AL662796.1, CXCL12, IL1B, IL12A, CCL25, CCL27, CCL28, CXCL14, LIF, IL18, LTB, CD40LG, IL15, IFNG, LTBR |
| GO:0005179 | hormone activity | 19 | 9.98E-07 | 6.28657492 | 0.00070484 | OG0001667, OG0002625, OG0002696, OG0003159, OG0003885, OG0005227, OG0006012, OG0006200, OG0006436, OG0007402, OG0008260, OG0009491, OG0009714, OG0009769, OG0010837, OG0010911, OG0011097, OG0013115, OG0013326 | PRL, NPPC, SST, CRH, UCN, PTHLH, ADCYAP1, CCK, UCN3, EPO, ADM, IAPP, VIP, CGA, PRLH, LEP, GNRH1, MLN |
| GO:0008009 | chemokine activity | 8 | 4.78E-06 | Inf | 0.00247506 | OG0000050, OG0000095, OG0004652, OG0009542, OG0009678, OG0009774, OG0011242, OG0012322 | CCL26, CCL22, CCL17, CCL24, CCL2, CCL13, CCL21, CCL11, CCL20, CCL7, CCL19, CCL8, CCL15, CCL3, CCL16, CCL4, CCL4L2, CCL18, CCL3L3, CCL3L1, CCL23, CCL15-CCL14, CCL14, CCL5, CXCL6, PF4V1, CXCL13, CXCL3, CXCL5, PPBP, PF4, CXCL10, CXCL8, CXCL11, CXCL9, CXCL1, CXCL2, CXCL12, CCL25, CCL27, CCL28, CXCL14 |
| PF00048 | Small cytokines (intecrine/chemokine), | 8 | 4.78E-06 | Inf | 0.00247506 | OG0000050, OG0000095, OG0004652, OG0009542, OG0009678, OG0009757, OG0011242, OG0012322 | CCL26, CCL22, CCL17, CCL24, CCL2, CCL13, CCL21, CCL11, CCL20, CCL7, CCL19, CCL8, CCL15, CCL3, CCL16, CCL4, CCL4L2, CCL18, CCL3L3, CCL3L1, CCL23, CCL15-CCL14, CCL14, CCL5, CXCL6, PF4V1, CXCL13, CXCL3, CXCL5, PPBP, PF4, CXCL10, CXCL8, CXCL11, CXCL9, CXCL1, |

| | interleukin-8 like | | | | | | CXCL2, CXCL12, CCL25, CCL27, CCL28 |
|---|---|---|---|---|---|---|---|

Supplementary table 9. Significantly-enriched functional and domain terms identified in novel gene families (orthogroups) gained in the MRCA of Osteichthyes. n refers to the number of these gene families with that function gained. p.value refers to uncorrected p-values for Fisher's exact test, adj.p refers to the adjusted p-value for multiple testing (See Methods). See Supplementary File 7 for specific assignments of human gene names to each orthogroup.

| domain id | domain description | n | p | estimate odds.ratio | adj.p | orthogroups | human gene names |
|---|---|---|---|---|---|---|---|
| PF00096 | Zinc finger, C2H2 type | 26 | 2.84E-08 | 3.98407764 | 0.00011148 | OG0000670, OG0003551, OG0010444, OG0010637, OG0010664, OG0010793, OG0010841, OG0010872, OG0011424, OG0011486, OG0011496, OG0011579, OG0011969, OG0012403, OG0012612, OG0012834, OG0013091, OG0013829, OG0014215, OG0014809, OG0014856, OG0015934, OG0016027, OG0016061, OG0016083, OG0017505 | ZNF787, ZNF581, ZNF524, ZNF408, ZNF629, ZNF697, ZNF770, REPIN1, ZNF597, ZNF691, ZNF229 |
| GO:0004930 | G protein-coupled receptor activity | 11 | 1.71E-06 | 7.27772534 | 0.00334684 | OG0000023, OG0000210, OG0000260, OG0003307, OG0013404, OG0013735, OG0014847, OG0015147, OG0015786, OG0018003, OG0018005 | OR6M1, OR6S1, OR2AP1, OR6C2, OR6T1, OR6C76, OR6C70, OR6X1, OR6C74, OR6C6, OR6C65, OR6C75, OR6C3, OR6C1, OR6C4, OR6V1, OR6J1, OR6C68, OR13C9, OR13C4, OR13D1, OR13C8, OR13F1, OR2S2, OR2K2, OR13C5, OR13C3, OR13J1, OR13C2, MAS1, MRGPRX1, MRGPRF, MRGPRD, MRGPRX4, MRGPRG, MRGPRX2, MAS1L, MRGPRE, MRGPRX3 |
| GO:0003676 | nucleic acid binding | 24 | 2.05E-05 | 2.83216665 | 0.02229267 | OG0003551, OG0010444, OG0010664, OG0010793, OG0011286, OG0011384, OG0011424, OG0011486, OG0011496, OG0011579, OG0012403, OG0012612, OG0012834, OG0013075, OG0013793, OG0014215, OG0014809, OG0016027, OG0016061, OG0016083, OG0016446, OG0017477, OG0017503, OG0017505 | ZNF787, ZNF581, ZNF524, ZNF408, ZNF629, THAP9, ZBTB9, ZNF770, REPIN1, ZNF597, ZNF691, ZNF229 |
| GO:0007186 | G protein-coupled receptor signaling pathway | 11 | 2.67E-05 | 5.23802374 | 0.02229267 | OG0000023, OG0000210, OG0000260, OG0003307, OG0013404, OG0013735, OG0014847, OG0015147, OG0015786, OG0018003, OG0018005 | OR6M1, OR6S1, OR2AP1, OR6C2, OR6T1, OR6C76, OR6C70, OR6X1, OR6C74, OR6C6, OR6C65, OR6C75, OR6C3, OR6C1, OR6C4, OR6V1, OR6J1, OR6C68, OR13C9, OR13C4, OR13D1, OR13C8, OR13F1, OR2S2, OR2K2, OR13C5, OR13C3, OR13J1, OR13C2, MAS1, MRGPRX1, MRGPRF, MRGPRD, MRGPRX4, MRGPRG, MRGPRX2, MAS1L, MRGPRE, MRGPRX3 |
| PF05699 | hAT family | 4 | 5.48E-05 | 31.9512668 | 3.91E-02 | OG0012361, OG0012385, OG0013080, OG0013383 | |

| | |
|---|---|
| | C-terminal dimerisation region |

Supplementary table 10. Significantly-enriched functional and domain terms identified in gene families (orthogroups) lost in the MRCA of Chondrichthyes. n refers to the number of these gene families with that function gained. p.value refers to uncorrected p-values for Fisher's exact test, adj.p refers to the adjusted p-value for multiple testing (See Methods). See Supplementary File 7 for specific assignments of human gene names to each orthogroup.

| domain id | domain description | n | p | estimate odds.ratio | adj.p | orthogroups | human gene names |
|---|---|---|---|---|---|---|---|
| PF00096 | Zinc finger, C2H2 type | 46 | 6.86E-28 | 10.6206996 | 5.33E-24 | OG0000225, OG0000956, OG0002249, OG0002987, OG0003779, OG0004959, OG0009283, OG0010416, OG0010626, OG0010851, OG0010982, OG0011022, OG0011065, OG0011067, OG0011222, OG0011453, OG0011584, OG0011599, OG0011742, OG0011821, OG0011865, OG0011944, OG0011965, OG0012017, OG0012140, OG0012762, OG0013057, OG0013171, OG0013175, OG0013220, OG0013514, OG0013893, OG0013896, OG0014096, OG0014287, OG0014339, OG0014359, OG0014363, OG0014386, OG0014947, OG0015058, OG0015104, OG0016274, OG0016327, OG0016335, OG0016357 | ZNF416, ZNF132, ZNF773, ZNF551, ZNF256, ZNF211, ZNF17, ZNF417, ZNF776, ZNF671, ZNF530, ZNF749, ZNF548, ZNF772, ZNF587, ZNF549, ZNF552, ZNF586, ZNF134, ZNF792, ZNF419, ZNF304, ZNF418, ZNF814, ZNF154, ZNF587B, AC004076.1, AC010522.1, AC003005.1, AC003006.1, AC003002.3, ZIK1, AC003002.1, ZNF777, ZNF212, ZNF746, ZNF783, ZNF398, ZNF282, NA, ZNF837, ZNF774, ZNF646, SNAI3, GLI4, ZNF672, ZNF32, ZFP69B, ZFP69, SALL2, ZNF775, ZNF775, ZNF683, ZNF275, ZNF589, AL049650.2, ZNF343, ZNF771, ZNF146, ZFP3 |
| PF00059 | Lectin C-type domain | 10 | 8.12E-09 | 16.5125796 | 3.15E-05 | OG0000432, OG0001053, OG0001906, OG0003408, OG0012083, OG0012148, OG0012681, OG0013467, OG0013495, OG0014331 | CD69, CLEC2B, CLEC2D, KLRG1, CLEC2L, CLEC2A, SFTPD, SFTPA2, MBL2, SFTPA1, REG1A, REG4, REG3G, REG1B, REG3A, KLRC4, KLRC1, KLRC2, KLRC3, AC068775.1, KLRC4-KLRK1, CLEC5A |
| PF13912 | C2H2-type zinc finger | 7 | 6.14E-06 | 12.6103152 | 1.56E-02 | OG0004959, OG0011065, OG0011453, OG0012762, OG0014096, OG0014359, OG0014947 | ZNF646, SALL2 |
| PF01352 | KRAB box | 4 | 8.03E-06 | 71.4512335 | 1.56E-02 | OG0000225, OG0000956, OG0011022, OG0011742 | ZNF416, ZNF132, ZNF773, ZNF551, ZNF256, ZNF211, ZNF17, ZNF417, ZNF776, ZNF671, ZNF530, ZNF749, ZNF548, ZNF772, ZNF587, ZNF549, ZNF552, ZNF586, ZNF134, ZNF792, ZNF419, ZNF304, ZNF418, ZNF814, ZNF154, ZNF587B, AC004076.1, AC010522.1, AC003005.1, AC003006.1, AC003002.3, ZIK1, AC003002.1, ZNF777, ZNF212, ZNF746, ZNF783, ZNF398, ZNF282, ZFP69B, ZFP69, ZNF589, AL049650.2, ZNF343 |

Supplementary table 11. Significantly-enriched functional and domain terms identified in gene families (orthogroups) with a rate shift in gene family size in any part of the vertebrate phylogeny. n refers to the number of these gene families with that function gained. p.value refers to uncorrected p-values for Fisher's exact test, adj.p refers to the adjusted p-value for multiple testing (See Methods). See Supplementary File 7 for specific assignments of human gene names to each orthogroup.

| domain id | domain description | n | p | estimate odds.ratio | adj.p | orthogroups | human gene names |
|---|---|---|---|---|---|---|---|
| GO:0005840 | ribosome | 43 | 5.92E-15 | 5.87923953 | 3.48E-11 | OG0000627, OG0001253, OG0001296, OG0001778, OG0001872, OG0001911, OG0002023, OG0002040, OG0002382, OG0002504, OG0002728, OG0002888, OG0002896, OG0003081, OG0003102, OG0003220, OG0003345, OG0003528, OG0003538, OG0003738, OG0003748, OG0003757, OG0003881, OG0003893, OG0003939, OG0004038, OG0004135, OG0004191, OG0004254, OG0004270, OG0004348, OG0004738, OG0004764, OG0004820, OG0004891, OG0005109, OG0005437, OG0006163, OG0006834, OG0007395, OG0009978, OG0010750, OG0010807 | RPL23A, RPL17, RPL17-C18orf32, RPSA, RPL10L, RPL10, RPL22, RPL22L1, RPS27L, RPL31, RPL21, RPL9, RPS4Y1, RPS4X, RPS4Y2, RPL32, RPL26L1, AC135178.3, RPS15A, RPL36AL, RPL36A-HNRNPH2, RPL36A, RPS6, RPL12, RPS26, RPS24, RPS3A, FAU, RPL23, RPL37, RPL6, RPS23, RPS17, RPS2, RPS11, RPL29, RPL4, RPL36, RPS14, RPL35, RPL37A, RPS21, RPS13, RPS18, RPL39L, RPL39, RPS29, RPS28, MRPS18C, MRPL36 |
| GO:0006412 | translation | 43 | 2.52E-14 | 5.53161738 | 9.89E-11 | OG0000627, OG0001253, OG0001296, OG0001778, OG0001872, OG0001911, OG0002023, OG0002040, OG0002382, OG0002504, OG0002728, OG0002888, OG0002896, OG0003081, OG0003102, OG0003220, OG0003345, OG0003528, OG0003538, OG0003738, OG0003748, OG0003757, OG0003881, OG0003893, OG0003939, OG0004038, OG0004135, OG0004191, OG0004254, OG0004270, OG0004348, OG0004738, OG0004764, OG0004820, OG0004891, OG0005109, OG0005437, OG0006163, OG0006834, OG0007395, OG0009978, OG0010750, OG0010807 | RPL23A, RPL17, RPL17-C18orf32, RPSA, RPL10L, RPL10, RPL22, RPL22L1, RPS27L, RPL31, RPL21, RPL9, RPS4Y1, RPS4X, RPS4Y2, RPL32, RPL26L1, AC135178.3, RPS15A, RPL36AL, RPL36A-HNRNPH2, RPL36A, RPS6, RPL12, RPS26, RPS24, RPS3A, FAU, RPL23, RPL37, RPL6, RPS23, RPS17, RPS2, RPS11, RPL29, RPL4, RPL36, RPS14, RPL35, RPL37A, RPS21, RPS13, RPS18, RPL39L, RPL39, RPS29, RPS28, MRPS18C, MRPL36 |
| GO:0003735 | structural constituent of ribosome | 43 | 3.63E-13 | 4.94618578 | 1.07E-09 | OG0000627, OG0001253, OG0001296, OG0001778, OG0001872, OG0001911, OG0002023, OG0002040, OG0002382, OG0002504, OG0002728, OG0002888, OG0002896, OG0003081, OG0003102, OG0003220, OG0003345, OG0003528, OG0003538, OG0003738, OG0003748, OG0003757, OG0003881, OG0003893, OG0003939, OG0004038, OG0004135, OG0004191, OG0004254, OG0004270, OG0004348, OG0004738, | RPL23A, RPL17, RPL17-C18orf32, RPSA, RPL10L, RPL10, RPL22, RPL22L1, RPS27L, RPL31, RPL21, RPL9, RPS4Y1, RPS4X, RPS4Y2, RPL32, RPL26L1, AC135178.3, RPS15A, RPL36AL, RPL36A-HNRNPH2, RPL36A, RPS6, RPL12, RPS26, RPS24, RPS3A, FAU, RPL23, RPL37, RPL6, RPS23, RPS17, RPS2, RPS11, RPL29, RPL4, RPL36, RPS14, RPL35, RPL37A, RPS21, RPS13, RPS18, RPL39L, RPL39, RPS29, RPS28, MRPS18C, MRPL36 |

| | | | | | | | |
|---|---|---|---|---|---|---|---|
| | | | | | | OG0004764, OG0004820, OG0004891, OG0005109, OG0005437, OG0006163, OG0006834, OG0007395, OG0009978, OG0010750, OG0010807 | |
| PF12775 | P-loop containing dynein motor region | 6 | 3.75E-05 | 38.4900634 | 0.04010636 | OG0000048, OG0000290, OG0000794, OG0003524, OG0011461, OG0016374 | DNAH3, DNAH7, DNAH2, DNAH14, DNAH1, DNAH12, DNAH9, DNAH11, DNAH17, DNAH5, DNAH8, DNAH10 |
| PF12777 | Microtubule-binding stalk of dynein motor | 6 | 3.75E-05 | 38.4900634 | 0.04010636 | OG0000048, OG0000290, OG0000794, OG0003483, OG0003524, OG0012835 | DNAH3, DNAH7, DNAH2, DNAH14, DNAH1, DNAH12, DNAH9, DNAH11, DNAH17, DNAH5, DNAH8, DNAH10 |
| PF12780 | P-loop containing dynein motor region D4 | 6 | 3.75E-05 | 38.4900634 | 0.04010636 | OG0000048, OG0000290, OG0000794, OG0003483, OG0003524, OG0016374 | DNAH3, DNAH7, DNAH2, DNAH14, DNAH1, DNAH12, DNAH9, DNAH11, DNAH17, DNAH5, DNAH8, DNAH10 |
| PF12781 | ATP-binding dynein motor region | 6 | 3.75E-05 | 38.4900634 | 0.04010636 | OG0000048, OG0000290, OG0000794, OG0003483, OG0003524, OG0012287 | DNAH3, DNAH7, DNAH2, DNAH14, DNAH1, DNAH12, DNAH9, DNAH11, DNAH17, DNAH5, DNAH8, DNAH10 |
| PF17857 | AAA+ lid domain | 5 | 4.49E-05 | Inf | 0.04405071 | OG0000290, OG0000794, OG0003524, OG0011461, OG0016374 | DNAH9, DNAH11, DNAH17, DNAH5, DNAH8, DNAH10 |
